# Somatic DNA-repeat expansions initiate distinct yet analogous changes in Huntington’s disease cortical and striatal neurons

**DOI:** 10.64898/2025.12.09.688862

**Authors:** Seva Kashin, Won-Seok Lee, Tara M. McDonald, Kiely Morris, Robert E. Handsaker, Curtis J. Mello, Liv Spina, Nora M. Reed, Heather de Rivera, Sri Havya Jana, Marina Hogan, Sabina Berretta, Steven A. McCarroll

## Abstract

Huntington’s disease (HD) affects the neocortex and striatum with different timing, severity, and gene-expression changes. These pathologies have long been proposed to involve different pathogenic mechanisms, with one brain area’s degeneration leading to the other’s. Striatal neurons degenerate in HD as their huntingtin (*HTT*) CAG-repeat tracts expand somatically beyond 150 CAGs. Here we find that cortical pyramidal neurons in HD undergo biological changes analogous to those in striatal neurons – the same sequence of pathological phases, with toxicity also commencing at ∼150 CAGs yet then changing their gene expression in distinct ways. Pyramidal neurons’ precise molecular identities, rather than their areal or laminar locations, appeared to determine the extent of this somatic DNA-repeat expansion, which varied up to 20-fold among pyramidal neuron subtypes. Cortical and striatal neurons also shared a key pathological feature: nuclear inclusions of aggregated HTT protein in those nuclei with expansions well beyond 150 CAGs.

## Introduction

Huntington’s disease (HD) is a neurodegenerative disease of progressive loss of neurons in the striatum and neocortex, leading to escalating motor, cognitive and psychiatric impairments. The most prominent disease-associated change is atrophy of the striatum (caudate and putamen) due to loss of striatal spiny projection neurons (SPNs, also called medium spiny neurons or MSNs). HD also affects the neocortex, causing loss of pyramidal projection neurons, especially in deep cortical layers ^1–6^, which harbor a variety of pyramidal neuron types, including neurons whose corticostriatal-projecting axons contact and synapse onto SPNs.

Striatal and cortical pathology in HD appear different in most details. In striatum, HD causes loss of inhibitory (GABAergic) SPNs; in the neocortex, HD causes loss of excitatory (glutamatergic) pyramidal neurons ^5^. While most-to-all SPNs are lost in advanced HD, only a minority of cortical pyramidal neurons are lost. Profiling gene expression in striatum and cortex in persons with HD and in HD mouse models has not identified strong correlations between the gene-expression changes in the two brain areas, and studies suggest distinct mechanisms for the two ^7–9^. These and earlier observations have long been interpreted to suggest that striatal and cortical pathology in HD may have different mechanisms and moreover that, due to the axonal contacts between striatal and cortical neurons, pathology in one brain area elicits secondary pathology in the other brain area – for example, due to excitotoxicity ^10–12^, corticostriatal disconnection ^7,8,13,14^, or spreading proteins with prion-like properties ^15,16^.

HD is a genetically dominant condition caused by an inherited expansion of a DNA-repeat tract ([CAG]_n_) in the first exon of the huntingtin (*HTT*) gene. This inherited CAG-repeat tract is generally 15 to 30 CAGs long in unaffected individuals but contains 36 or more CAGs (most frequently 40-50) in all persons with HD ^17^. Age at onset of clinical HD symptoms is a more complex trait: in addition to being shaped by inherited CAG-repeat tract length (longer inherited CAG-repeat tracts associate with earlier onset), age at clinical motor symptom onset is also shaped by common genetic variants at many loci. Several of the implicated loci contain genes that affect somatic DNA repeat instability ^18–30^.

We recently described experimental results nominating a model for striatal HD pathology in which particularly long somatic expansions (>150 CAGs) of this inherited CAG repeat tract, attained via many length increases that stochastically occur over decades, eventually – after expansion beyond about 150 CAGs – lead to a striatal neuron’s pathology and then death ^31^. First, the CAG-repeat tract expands somatically over decades via many small length-change mutations (phase A). Upon expanding beyond about 90 CAGs, expansion accelerates, passing 150 CAGs in just a few more years (phase B). When a neuron’s CAG-repeat tract expands beyond about 150 CAGs, the neuron begins to exhibit very specific changes in gene expression, affecting expression of hundreds of genes, with changes increasing in magnitude alongside further repeat expansion (phase C). Individual SPNs later enter a “de-repression crisis” (phase D) in which scores of normally-silent genes are de-repressed. These SPNs are then lost (phase E).

It is not known whether such a dynamic could also explain cortical neurodegeneration in HD. Key challenges would be to explain the many ways in which cortical and striatal neurodegeneration differ – in timing, severity, cell-type specificity, and gene-expression changes.

To address these and other questions, we used an approach we invented in the earlier HD study for analyzing *HTT* CAG-repeat length and genome-wide RNA expression together at single-cell resolution ^31^. We analyzed more than 125,000 cell nuclei sampled *post mortem* from the neocortex of eleven persons with HD.

We find that cortical and striatal neurodegeneration actually do arise from analogous cell-autonomous events, each driven by somatic expansion of a cell’s own *HTT* CAG repeat; that somatic expansion beyond about 150 CAGs also initiates the cortical-neuron pathology; that this 150-CAG threshold is shared by pyramidal neurons of all types; and that cortical area, laminar locations, and axonal projections play only incidental roles. Instead, a pyramidal neuron’s precise molecular identity profoundly affects its somatic-expansion kinetics, determining the likelihood that the high threshold for toxicity is reached in a human lifetime. We also find that a long-mysterious pathological feature of HD – nuclear inclusions of aggregated HTT protein – arises in precisely those neurons with CAG-repeat tracts longer than 150 CAGs.

## Results

### Long repeat expansions, beyond 100 CAGs, in glutamatergic neurons

In the striatum, HD causes loss of SPNs while sparing other cells; this cell-type specificity corresponds to the cell-type specificity of long somatic expansions (beyond 150 CAGs), which manifest almost exclusively in SPNs ^31^. (Cholinergic interneurons exhibit moderate somatic expansion^7^ but relatively few of them reach ∼150 CAGs, whereas large fractions of SPNs do so ^31^.)

In the neocortex in HD, the degenerating cells are pyramidal projection neurons ^2–4^, which are excitatory (glutamatergic) and differ markedly in gene expression and physiology from SPNs, which are inhibitory GABAergic projection neurons.

To assess the cell-type specificity of somatic expansion in the neocortex, we measured CAG-repeat length and genome-wide RNA expression together at single-cell resolution, using a laboratory approach we recently developed ^31^. We generated such paired measurements for 70,760 nuclei from 11 persons with HD – sampling motor cortex (Brodmann Area 4, BA4) and anterior cingulate cortex (BA32) – as well as 56,889 nuclei sampled from 10 additional cortical areas (the 12 cortical areas are described in **Suppl. Table 1**) of one of these brain donors, a 48yo person with HD (Vonsattel grade HD-1, “Donor B” in **Fig. 1**) who had inherited a CAG-repeat tract of 42 CAGs.

**Figure 1.**
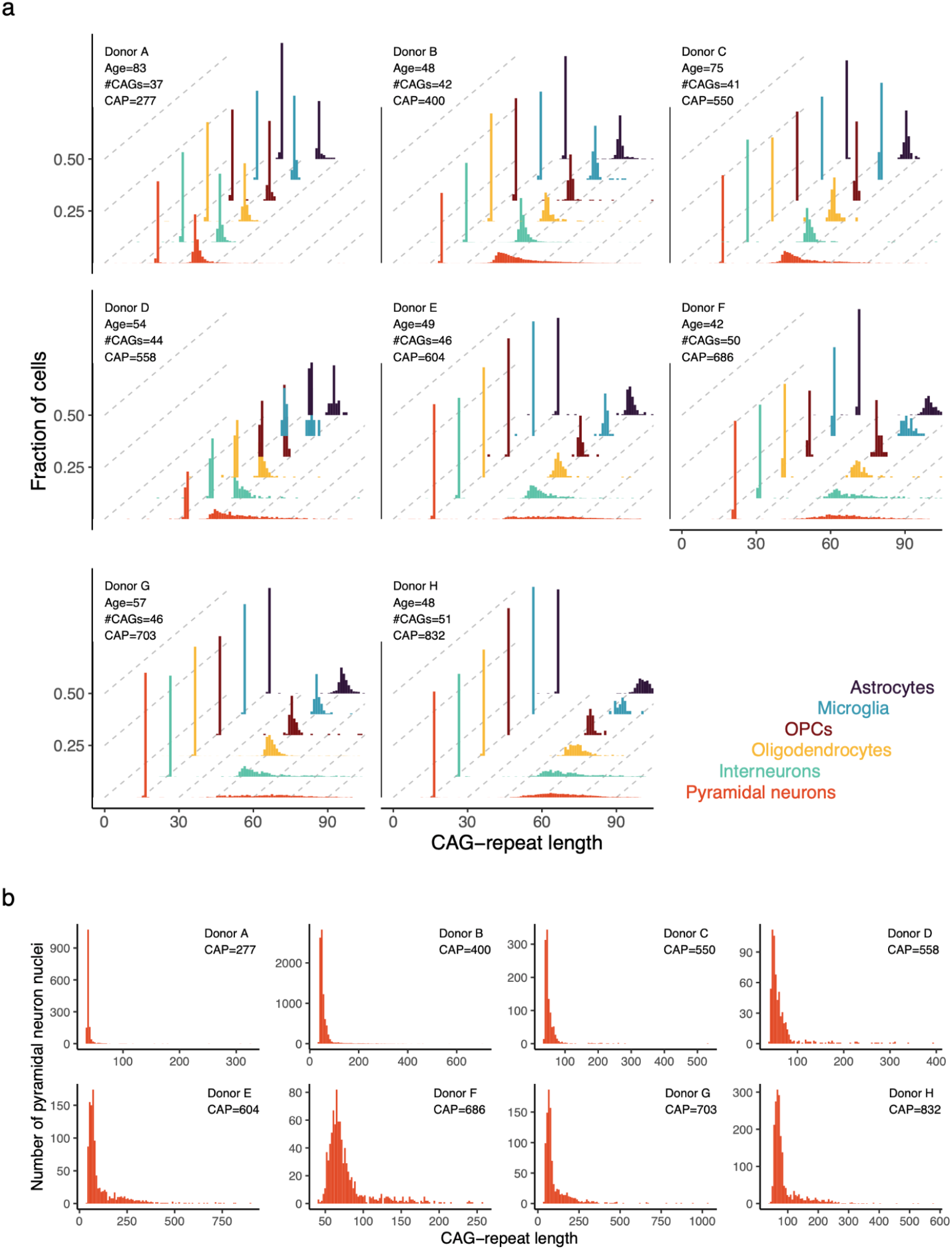
Somatic CAG-repeat instability in the neocortex, by cell type. (**a**) Cell-type-specific distributions of CAG-repeat length in cell types of the neocortex (BA4) from eight brain donors with HD. The donors are ordered (A..H) based on increasing CAG-age-product (CAP) score, an estimate of HD prognosis/progression based on age and inherited CAG-repeat length. Length distributions for both alleles are shown, though the shorter (non-HD-causing) allele appears to have been stable in each donor. (**b**) As in panel a, but focusing on pyramidal neurons and on the longer (HD-causing) allele, and extending the x-axis to capture the full distribution of observed CAG-tract lengths in each donor.

As in the striatum, somatic expansion had affected the HD-causing allele but not the other (shorter) allele, which appeared to have been stable throughout life (**Fig. 1a**).

The HD-causing allele exhibited some CAG-repeat instability in all cell types, but expansion was by far the highest in pyramidal neurons (**Fig. 1**). This pattern replicated across all donors (**Fig. 1**) and across all cortical areas sampled (**Fig. 2a,b**). (Cortical interneurons exhibited the next most somatic expansion – mainly in donors who had inherited CAG-repeat tracts longer than 45 CAGs – but such neurons seldom reached even 100 CAGs (**Fig. 1a and Fig. S1**).)

**Figure 2.**
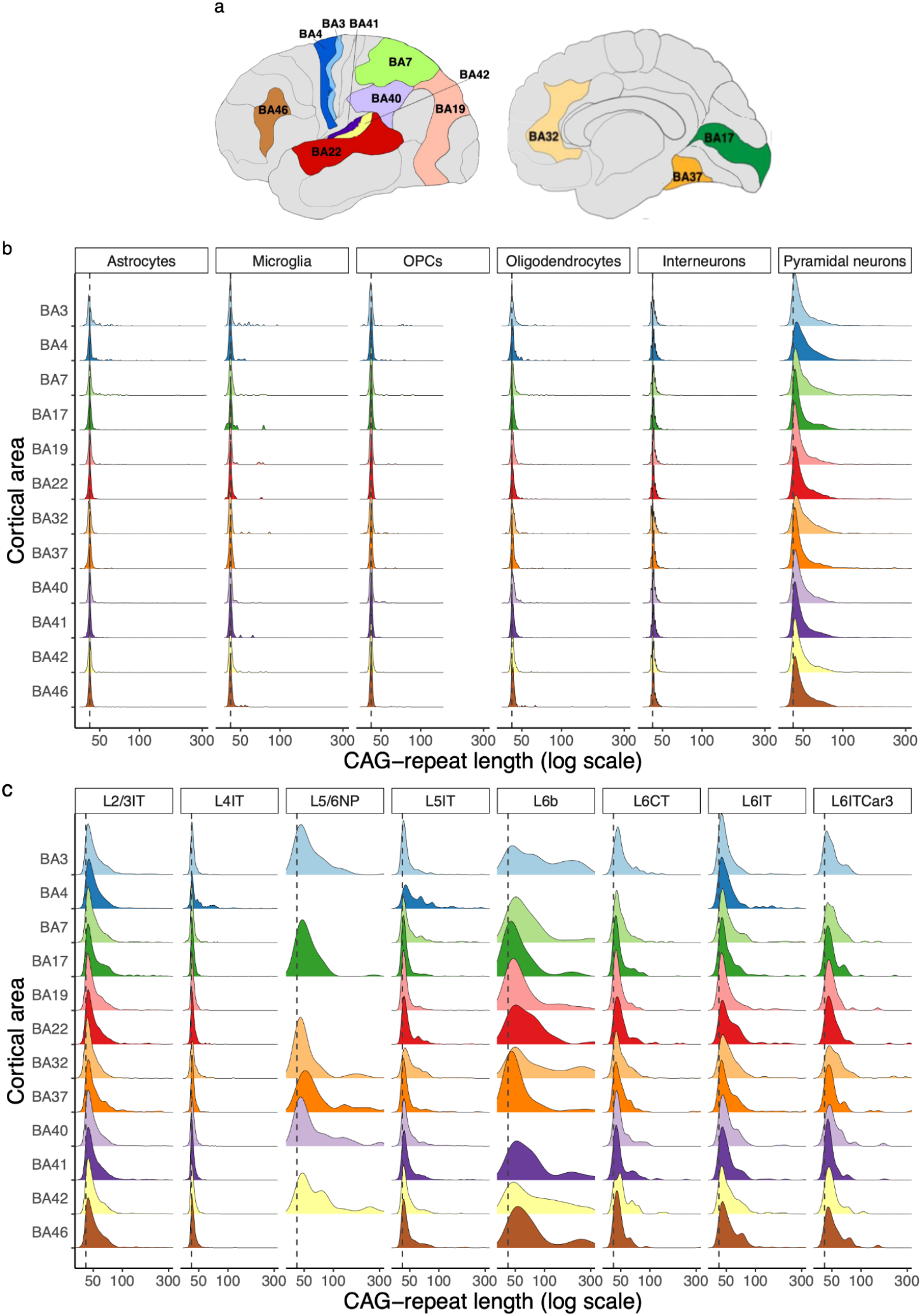
Somatic CAG-repeat expansion across cortical areas and pyramidal neuron types (**a**) The twelve cortical areas analyzed in one of the above donors (donor B). BA3, primary somatosensory; BA4, primary motor; BA7, multisensory visual/somatosensory associative; BA17, primary visual; BA19, visual associative; BA22, auditory associative; BA32, anterior cingulate; BA37, visual high level associative; BA40, somatosensory associative; BA41, primary auditory; BA42, auditory associative; BA46, dorsolateral prefrontal. (**b**) CAG-repeat length distributions (focusing on the longer, HD-causing allele) for six cortical cell types (columns) in 12 cortical areas (rows). Colors correspond to the areas in panel a. Note that the areas under the curves are non-identical, as each curve’s highest point defines the top of the plotting area. (**c**) CAG-repeat length distributions for specific types of cortical pyramidal neurons, focusing on the longer, HD-causing allele. Density curves as in b. Curves are shown for those region/celltype combinations for which we obtained measurements of the HD-causing allele in at least 20 cells.

Thus, long somatic expansions – exceeding 100 CAGs in length – arise primarily among the cortical neurons that are lost in large numbers in HD (**Fig. 1**), analogous to the pattern we have described in the striatum ^31^.

### Somatic expansion varies profoundly across pyramidal neuron types

Neuroimaging and stereological studies have found that deeper cortical layers (5 and 6) exhibit more neuronal loss in HD than upper layers (2,3,4) do ^4^, and that severity of atrophy also varies by cortical area. The appearance of areal and laminar differences in vulnerability could indeed be due to direct effects of cortical area or layer; alternatively, these differences might instead reflect much-stronger effects of precise neuronal identity – pyramidal-neuron types and subtypes – since each cortical layer and area contains pyramidal-neuron types and subtypes in different quantities .

Cortical neurons are increasingly classified, based on RNA-expression patterns, into taxonomies of types and subtypes at ever-finer levels of resolution ^32–35^. The early snRNA-seq taxonomies (such as Hodge, e.g. refs^32,34^) have been annotated with anatomic information on laminar locations and axonal projections; a more recent taxonomy ^33^ has defined a more fine-grained hierarchy of types and subtypes, but with less annotation information so far.

We first used the Hodge taxonomy^32^ as a scaffold for analysis of somatic expansion in eight major pyramidal neuron types (those that were sufficiently well-represented for statistically well-powered analysis). These types of pyramidal neurons varied greatly in the amounts of somatic CAG-repeat expansion that they had acquired over a lifetime (**Fig. 2c**). For example, in the anterior cingulate cortex (BA32), 43% of layer 6b (L6b) neurons had expanded their CAG-repeat tract beyond 100 CAGs, but only 0.5% and 0.9% of layer 4 and layer 2/3 intratelencephalic-projecting neurons (L4IT and L2/3IT, respectively) had done so (**Fig. 2c**).

The pyramidal neuron types that were high-expansion or low-expansion in one cortical area had this same property in all cortical areas analyzed (**Fig. 2c**). For example, layer 6b (L6b) pyramidal neurons (fifth column in **Fig. 2c**) were the most repeat-expanded pyramidal type in all cortical areas in which they were sufficiently numerous for conclusive analysis, whereas layer 4 intratelencephalic-projecting (L4IT) neurons (second column in **Fig. 2c**) were the least repeat-expanded type in all cortical areas.

These results reveal that somatic expansion rates are intrinsic features of pyramidal neurons’ molecular identities, shared across cortical areas.

Across cortical areas, two types of pyramidal neurons – layer 6b (L6b) and layer 5/6 near-projecting (L5/6NP) neurons – exhibited far more somatic CAG-repeat expansion than other neurons did (**Fig. 2c**). These high-somatic-expansion pyramidal types (L6b, L5/6NP) were characterized not only by high mean expansion but also by high variance (in expansion) across individual cells (**Fig. 2c**), likely reflecting a dynamic (described and modeled previously ^31^) in which incremental length-change mutations are stochastic (random) and initially infrequent, but increase in frequency with increasing length

The axonal projections of pyramidal neuron types have been systematically characterized by retrograde labeling experiments in mice ^34^. Such experiments have found that L6b neurons project intracortically and to the thalamus, and that L5/6NP neurons project intracortically. Corticostriatal projections have been found among L5IT and L6IT neurons ^34^, which exhibited just intermediate amounts of somatic expansion (**Fig. 2c**).

The above results (**Fig. 2c**) suggest a strong effect of a pyramidal neuron’s precise molecular identity, relative to the effect of cortical area. To rigorously measure such effects, and to expand analysis to a more fine-grained set of pyramidal-neuron molecular identities (subtypes), we performed multiple negative binomial regression analyses, modelling the CAG-repeat tract’s somatic length increase in each individual neuron against independent variables for that neuron’s cortical area, donor and pyramidal neuron subtype. We used the larger number of fine-grained pyramidal-neuron molecular identities (subtypes or “cluster names”) in the Siletti taxonomy ^33^. (We also mapped each of the Siletti clusters to its corresponding supertype (L2/3IT, L5/6NP, etc.) in the earlier Hodge taxonomy by matching RNA-expression patterns (see Methods), to benefit from spatial and connectivity characterizations of the Hodge taxonomy.)

We first analyzed 46,665 neuronal nuclei sampled from 12 cortical areas of a single donor (donor B in **Fig 1**). Analysis revealed that cortical area had at most a modest effect on a neuron’s propensity for repeat expansion, with cortical areas varying by only up to 1.3-fold in somatic expansion when controlling for the molecular identities of the neurons they contained (**Fig. 3a**).

**Figure 3.**
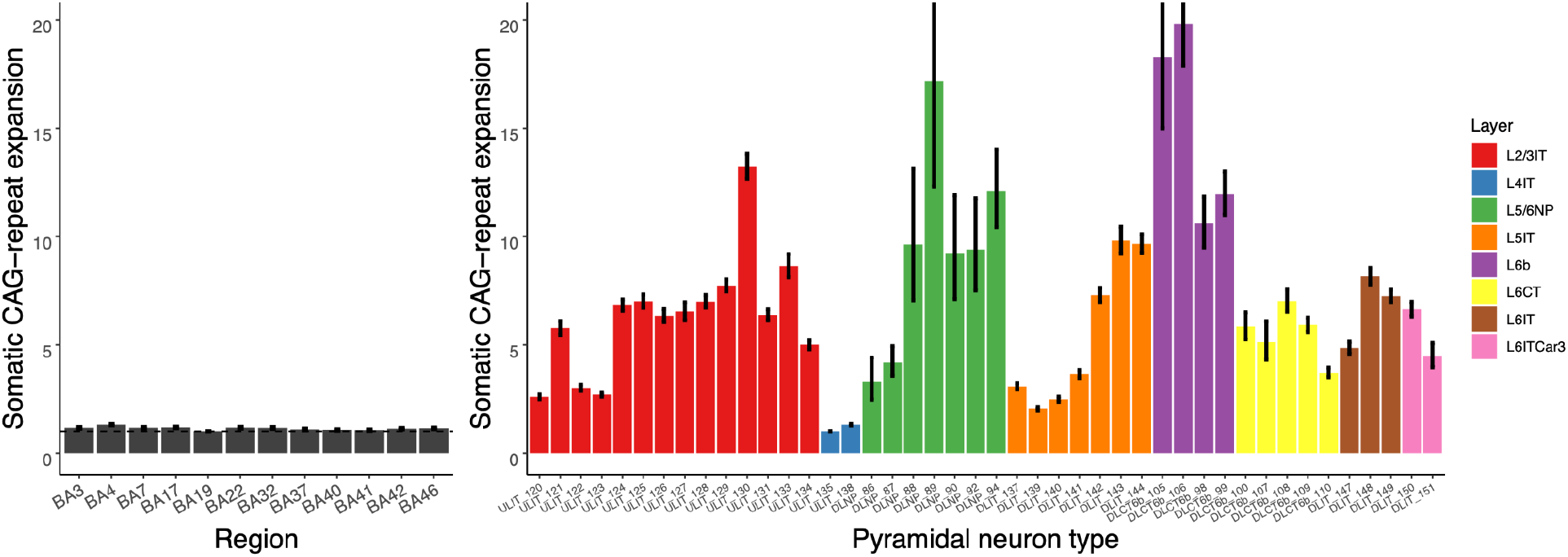
Effects of cortical area/region, molecular identity (pyramidal neuron type), and laminar location (layer) upon cortical glutamatergic neurons’ somatic CAG-repeat expansion. The heights of the bars represent the exponentials of the model coefficients inferred from a regression model (fitted to data from 46,665 neurons) in which a neuron’s somatic expansion (the log of its change in CAG-tract length relative to its initial/inherited length) is regressed against explanatory variables for (**a**) the cortical area from which it was sampled and (**b**) its molecular identity. The 44 molecular identities in (**b**) correspond to clusters in the Siletti taxonomy, each of which we also connected (via RNA expression patterns) to its corresponding glutamatergic subclass (L2/3IT, L5/6NP, etc., represented by colors) in the earlier Hodge taxonomy.

In contrast to the modest effect of cortical area, a neuron’s molecular identity (RNA-expression-defined subtype) had a profound effect, with pyramidal subtypes exhibiting up to 20-fold variation in the amounts of somatic expansion their CAG-repeat tracts had undergone (**Fig. 3b**) (This appears to explain their still-greater variation in tendency to have expanded beyond 100 CAGs, described above.)

To assess whether these relative expansion rates were intrinsic and general (as opposed to donor-specific) features of these neuronal subtypes, we also performed two separate Negative Binomial Regression analyses of 11,953 neuronal nuclei sampled from motor cortex (BA4) of ten additional brain donors with HD as well as 18,991 neuronal nuclei sampled from cingulate cortex (BA32) of seven additional brain donors with HD. The resulting estimates of the neuronal types’ relative expansion propensities were highly correlated across these analyses, regardless of the donor(s) or cortical area(s) in which they were measured (**Fig. S2**), reinforcing the conclusion that somatic-expansion rates are persistent features of pyramidal neurons’ precise molecular identities.

The laminar locations of molecularly defined pyramidal-neuron types have been characterized by earlier spatial transcriptomics analyses ^35^, making it possible to ask about the relationship of somatic expansion to cortical layer. The pyramidal-neuron types with the most somatic CAG-repeat expansion (L6b, L5/6NP) are among the types found in deep layers 5 and 6, aligning with longstanding observations that HD pathology affects deep layers earlier and more severely than it affects upper layers ^4,6^. However, all cortical layers contain mixtures of pyramidal neuron subtypes that our analyses found to have a variety of expansion rates. For example, upper-layer (L2/3IT) projection neurons include L2/3IT subtypes with diverse amounts of expansion (red bars in **Fig. 3b**, see also **Fig. S4**), and deep-layer L5/6NP neurons include some L5/6NP subtypes (shortest green bars in **Fig. 3b**, see also **Fig. S4**) with little somatic expansion.

Our experimental results indicate that molecularly defined types and subtypes of pyramidal neurons have diverse intrinsic propensities for somatic expansion that are consistent across cortical areas, consistent across people, that correlate only partially with laminar locations, and that do not distinguish corticostriatal projection neurons from other types of pyramidal neurons.

Somatic expansion could in principle be better explained by one or more features of pyramidal neurons’ fine-grained molecular identities, such as activity patterns or metabolic properties, which generally have yet to be characterized ^36^. We tested one simple possibility – that the nuclear RNA expression levels of *HTT* or one of several DNA-repair genes might substantially explain this variation. Nuclear RNA expression levels of most of these genes exhibited just subtle variation across pyramidal neuron types; this variation did not appear to explain the large variation in somatic DNA-repeat expansion among these neuronal types (**Fig. S5**). (Of course, it remains possible that complex and non-linear combinations of these variables are more predictive, but this is challenging to test in a rigorous way.)

### Somatic expansion occurs in two phases (A and B), as in SPNs

In striatum, SPN CAG-repeat lengths exhibit a mathematical distribution that visually resembles the profile of an armadillo, with a large majority of neurons having expanded to 45-90 CAGs (the body of the distribution), and a small fraction having attained far-higher CAG-repeat lengths of 100 to 500+ CAGs (the tail) at any one time ^31^. These distributions appear to arise from a two-stage somatic-expansion process in which somatic expansion is initially slow (phase A, requiring decades to expand from 40 to 90 CAGs) but accelerates – beyond its general pattern of continuous increase with repeat length – after expansion past about 90 CAGs (phase B) ^31^.

CAG-repeat-length distributions in cortical pyramidal neurons of all types with substantial somatic expansion exhibited similar shapes (**Fig. S6**), suggesting that this two-phase expansion regime is a shared property of cortical and striatal projection neurons.

### Phase C: gene expression pathology commences at about 150 CAGs

The key question for understanding HD pathology in the neocortex involves whether, when and how the length of the disease-causing CAG repeat affects the biology of cortical neurons. We thus sought to recognize relationships between the length of this CAG-repeat tract in individual neurons and the same neurons’ patterns of gene expression.

We performed negative binomial regression (NBR) analysis of each gene’s expression levels across all the individual neurons in the analysis, controlling for the effects of fine-grained molecular identities (of the neurons) and of donor-specific effects by including molecular identity and donor as additional independent variables. This analysis identified more than 400 genes whose expression levels were associated with the lengths of pyramidal neurons’ CAG-repeat tracts (**Data S1**).

As in the striatum, gene-expression changes were apparent in those neurons that had expanded their CAG-repeat tracts beyond about 150 CAGs (**Fig. 4a**; **Fig. S7,8**). This was true regardless of a neuron’s molecular identity, laminar location or reported connectivity (**Fig. 4a**). The magnitude of these gene-expression changes appeared to escalate alongside further CAG-repeat expansion (beyond 150 CAGs): Initially subtle (at 150-180 CAGs), these gene expression changes steadily escalated in magnitude alongside additional repeat expansion (**Fig. 4a**). Neurons with shorter repeat expansions (less than 150 CAGs) exhibited a more-random pattern consistent with statistical sampling noise and/or random biological fluctuations (**Fig. 4a**).

**Figure 4.**
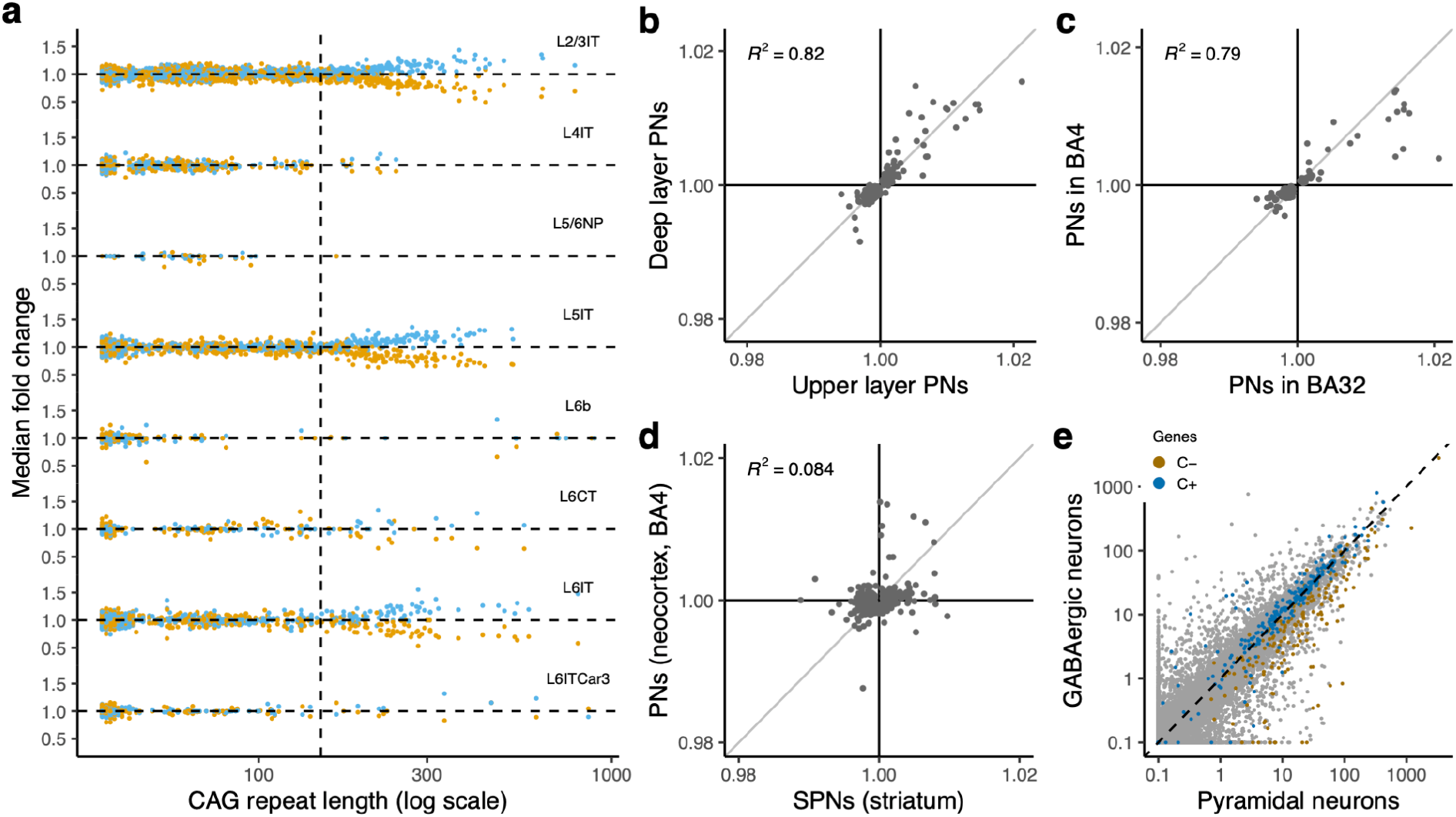
Phase C gene-expression changes in cortical pyramidal neurons. (**a**) Relationship of phase C gene-expression changes (y-axis) to a neuron’s own CAG-repeat length (x-axis), across eight types of pyramidal neurons (L2/3IT, L4IT, etc.; Hodge taxonomy). Each cortical pyramidal neuron is represented by both a blue point and an orange point: orange points show the median fold-change of 144 genes that decrease in expression with CAG-repeat expansion (C- genes); blue points show the median fold-change of 179 genes that increase in expression (C+ genes). The data are from analysis of anterior cingulate cortex (BA32) in five brain donors; analysis of a distinct donor (including 12 cortical areas) is in **Fig. S8**. (**b-d**) Comparisons of phase C gene-expression changes between (**b**) deep-layer vs. upper-layer pyramidal neurons, (**c**) pyramidal neurons in BA32 vs. BA4, and (**d**) cortical pyramidal neurons vs. striatal projection neurons. Each point corresponds to a gene that exhibited significant (p < 10^-10^) evidence of phase C gene-expression change in at least one of the two sets of cells being compared. The quantities plotted on the x- and y-axes are the fold-changes per CAG-repeat unit expansion (beyond 150 CAGs). (**e-f**) Genes that decline in expression during Phase C (C- genes, orange) are genes that, under normal circumstances, are much more strongly expressed in cortical pyramidal neurons (x-axis) than in cortical interneurons (y-axis). Gray points: all genes (cell-type-specific expression levels in unaffected individuals). Colored circles: Genes whose expression levels decline (C- genes, orange) or increase (C+ genes, blue) in neurons with *HTT* CAG-repeat expansion beyond 150 units (in phase C).

Critically, these gene-expression changes had affected pyramidal neurons *of all types*: at the single-neuron level, the critical variable involved was not molecular identity (or factors implicit in molecular identity – such as laminar location and axonal projections) but rather the extent to which a neuron’s CAG-repeat tract exceeded the 150-CAG threshold (**Fig. 4a**, **Fig. S7,8**). The effect of molecular identity was seen mainly in the fraction of neurons that had crossed this threshold (**Fig. 4a**; **Fig. S7,8**).

The ensuing gene-expression changes (that commenced upon a neuron’s crossing of the 150-CAG threshold) were also essentially identical across cortical layers, as could be seen when comparing fold-change estimates from NBR models fitted separately on the upper-layer and deep-layer pyramidal neurons (**Fig. 4b**). The changes were also essentially identical across cortical areas (**Fig. 4c**).

We call this period of continuously escalating transcriptional changes “phase C”. Its progressive dynamic could reflect either intensifying toxicity (with further expansion beyond 150 CAGs) or the amount of time since the toxicity threshold was crossed.

These results indicate that, despite the long-noted differences between cortical and striatal pathology in HD, cortical and striatal projection neurons in fact share a common CAG-repeat length toxicity threshold and pathological phases A, B, and C.

### Distinct Phase C gene-expression changes in cortical and striatal neurons

One of the mysteries that has long suggested the potential for distinct cellular mechanisms for cortical and striatal neuropathology in HD is that the pathological gene-expression changes in the two brain areas have been reported to exhibit little correlation with each other ^7–9^.

Although phase C gene-expression changes in cortical neurons were highly similar across cortical areas, and between deep-layer and upper-layer pyramidal neurons (*r*^2^ = 0.8, **Fig. 4bc**), and although phase C changes appeared to commence at the same CAG-length threshold in cortical neurons (∼150 CAGs, **Fig. 4a**) as in striatal neurons, phase C gene-expression changes in cortical neurons were quite different from phase C changes in striatal neurons (*r*^2^ = 0.084, **Fig. 4d**). Thus, although striatal and cortical neurons share a similar CAG-repeat-length threshold for the commencement of phase C changes, phase C perturbs their gene expression in quite different ways – with potentially quite-distinct effects on their physiologies and, downstream, on the other cells with which they interact.

### Phase C changes in cortical and striatal neurons involve identity erosion

In striatum, the hundreds of genes that decline in expression in SPNs during phase C share a specific property in common: almost all are genes whose expression distinguishes SPNs from other kinds of striatal neurons ^31^ (see also analysis of a mouse HD model ^37^).

The distinct set of genes that declined in expression in cortical pyramidal neurons during their phase C exhibited this same property: these genes (“C- genes”, orange points in **Fig. 4ef**) were genes whose expression distinguishes pyramidal neurons from neurons of other types. These genes encoded a wide variety of molecular functions. They included genes that normally exhibit categorical, on/off differences between pyramidal neurons and interneurons – such as *SV2B*, *ENC1*, and *LMO4* – as well as genes with large quantitative differences in expression levels (consistently higher in pyramidal neurons than in interneurons), such as *PRKCB*, *NR3C1*, *KALRN*, and *ATP2B1*. However, phase C appeared to spare expression of many genes that distinguish types of pyramidal neurons from one another – such as *CUX2* (expressed in upper-layer pyramidal neurons), *FEZF2* (expressed in deep-layer subcerebrally projecting pyramidal neurons), and *THEMIS* (expressed in deep-layer intratelencephalically projecting neurons). In fact, phase C gene-expression changes appeared to be shared almost identically between upper-layer and deep-layer pyramidal neurons, even when inferred in separate and independent analyses (**Fig. 4b**).

Notably, this strong relationship to pyramidal-neuron identity did not characterize the genes that increased in expression during phase C (“C+ genes”, blue points in **Fig. 4ef**), though C+ and C- genes had similarly high expression levels in pyramidal neurons (**Fig. 4ef**).

Thus, phase C gene-expression declines in striatal and cortical neurons are conceptually analogous and involve the same biological theme – erosion of gene-expression features of the neurons’ respective identities – while affecting different genes.

### Cortical neurons undergo a de-repression crisis (phase D)

In striatum, SPN pathology culminates, after still-further somatic CAG-repeat expansion, in de-repression of scores of genes that are normally silent in adult neurons ^31^. This “phase D” appears to follow the onset of a cell’s phase C by at least several months, as neurons with 150-250 CAGs have seldom entered phase D, whereas, with further repeat expansion (to 250+ CAGs) and progression of the phase C gene expression changes, ever-larger fractions of SPNs acquire this phase D de-repression phenotype ^31^.

Cortical pyramidal neurons with particularly long CAG-repeat expansions appeared to enter an analogous phase D, expressing very many genes that are usually silent in cortical pyramidal neurons (we call these D++ genes, due to their large fold-changes) (**Fig. 5a, Fig. S9a**). As in striatum, phase D was infrequent among neurons with repeat tracts of 150-250 CAGs, but was found with increasing frequency among neurons with still-longer CAG-repeat tracts (**Fig. 5b, Fig. S9b**). (Its frequency among neurons with <150 CAGs was below the well-recognized frequency of cell-barcode switching in the single-cell technology and thus considered negligible.)

**Figure 5.**
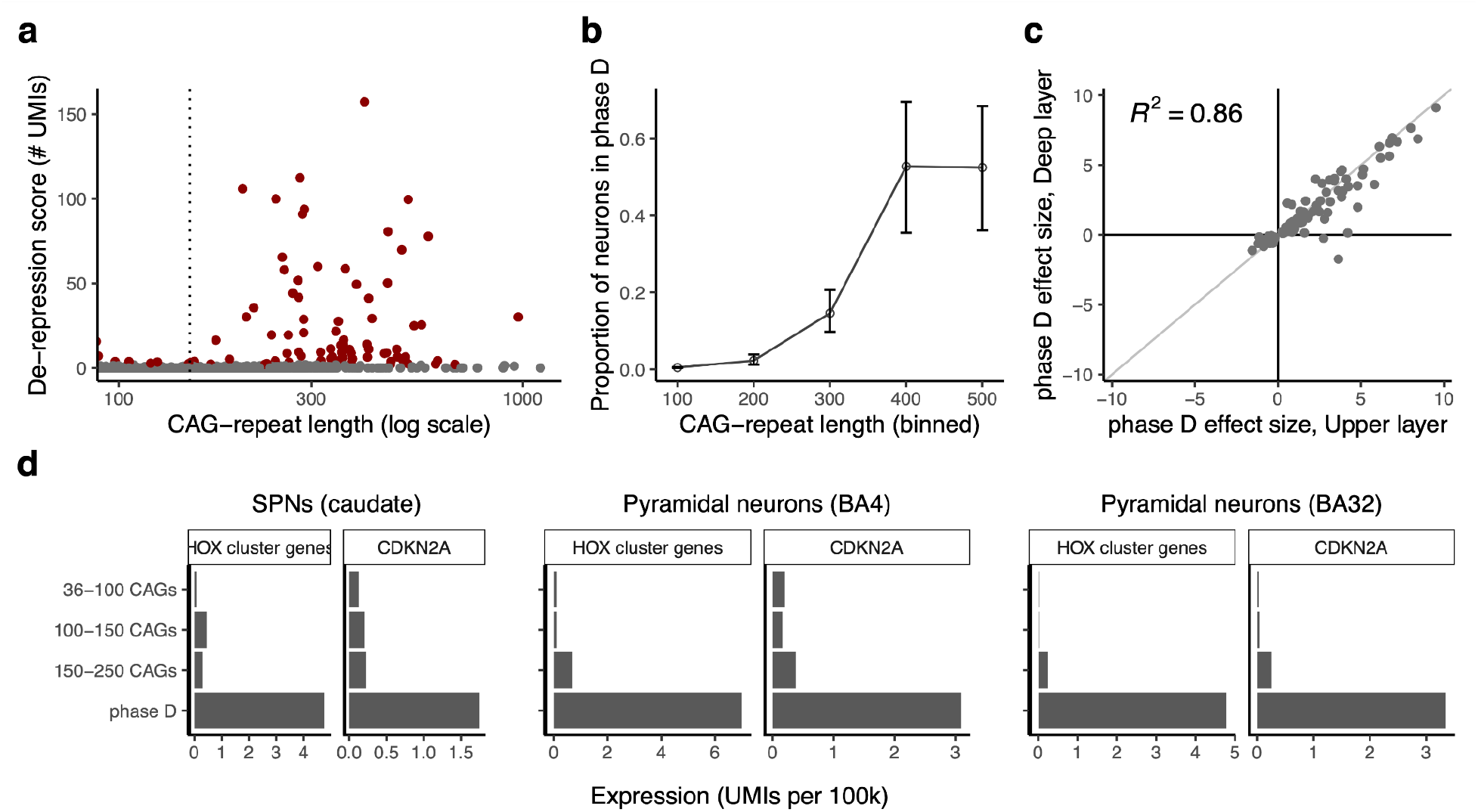
De-repression crisis (phase D). (**a**) De-repression of genes that are normally silent in cortical projection neurons. Points represent individual neuronal nuclei. y-axis: de-repression score, the number of transcripts (UMIs, normalized to 100k transcripts) detected from 59 phase D++ genes that are normally silent in cortical projection neurons. Replication analyses of data from additional persons with HD are in **Fig. S9**. (**b**) Proportion of cortical glutamatergic neurons that have entered phase D (y-axis), in relationship to the lengths of their *HTT* CAG-repeat tracts (x-axis), which is shown in bins. (**c**) Correlation of phase D gene-expression changes between upper-layer and deep-layer pyramidal neurons. Each point represents one gene; the point’s x-axis value represents that gene’s phase D expression log_2_-fold-change in upper-layer pyramidal neurons; the y-axis reports the same gene’s phase D log_2_-fold-change in deep-layer pyramidal neurons. Genes with significant phase D changes in at least one analysis are shown. Note the preponderance of de-repression events (large positive fold-changes). (**d**) Expression of HOX cluster genes and *CDKN2A* in SPNs, cortical pyramidal neurons from BA4, and cortical pyramidal neurons from BA32.

As in striatal neurons, phase D resembled a discrete crisis that contrasted with the gradually progressing changes of phase C. Most neurons with >150 CAGs continued to have minimal expression of phase D++ genes (more than 85% of such neurons had only 0-1 unique RNA molecules (unique molecular indices or UMIs, per 100k UMIs) detected across those 59 genes together. On the other hand, among neurons with more than 2 UMIs from phase D genes, more than 40% had at least 10 such UMIs and expressed up to 37 distinct phase D++ genes (**Fig. 5a, Fig. S9a**).

Moreover, unlike the phase C changes – whose magnitude was strongly correlated with the extent to which a neuron’s CAG-repeat tract length had extended beyond 150 CAGs (**Fig. 4a**) – the magnitude of the phase D changes did not associate with CAG-repeat length (**Fig. 5a, Fig. S9a**); rather, a neuron’s CAG-repeat length was associated with its probability of having entered phase D at all (**Fig. 5b, Fig. S9b**). All these results suggest that phase D involves a discrete crisis that cells enter rather suddenly, in contrast to the slowly, predictably escalating changes of phase C.

The intensity of the phase D++ changes – involving expression of very many genes that are usually silent – made it possible to identify a clear subset of nuclei as “phase D nuclei” (**Fig. 5a**), and then in turn to identify a larger set of phase D gene-expression changes (including more modest, quantitative changes, D+ and D- genes) that distinguished these nuclei from others (**Data S1**, **Methods**).

Like phase C changes, phase D gene-expression changes were shared almost identically between upper-layer and deep-layer pyramidal neurons (**Fig. 5c**). Unlike phase C changes, however, phase D++ changes were also largely shared with phase D++ changes in striatal neurons (**Fig. S9cd**).

Phase D pyramidal neurons greatly upregulated dozens of genes encoding transcription factors that are classic regulators of embryonic development, most of which had been silent in pyramidal neurons from unaffected individuals or (in HD) in neurons with 40-150 CAGs. These included more than 15 HOX cluster transcription factors and non-coding RNAs (Fig. 7d), which are normally involved in embryonic patterning; paired-domain developmental transcription factors (PAX1, PAX2, PAX3, PAX9), which regulate nervous system patterning and regional fate; LIM homeodomain transcription factors (LHX1, LHX4, MNX1), which regulate cell fate and migration during development; the homeodomain transcription factors ONECUT1, ONECUT2, SIX1, SIM2, SHOX2, PITX1, ZIC1, and OTP; and the non-homeobox transcription factors TBX3, TBX5 and FOXF1.

**Figure 6.**
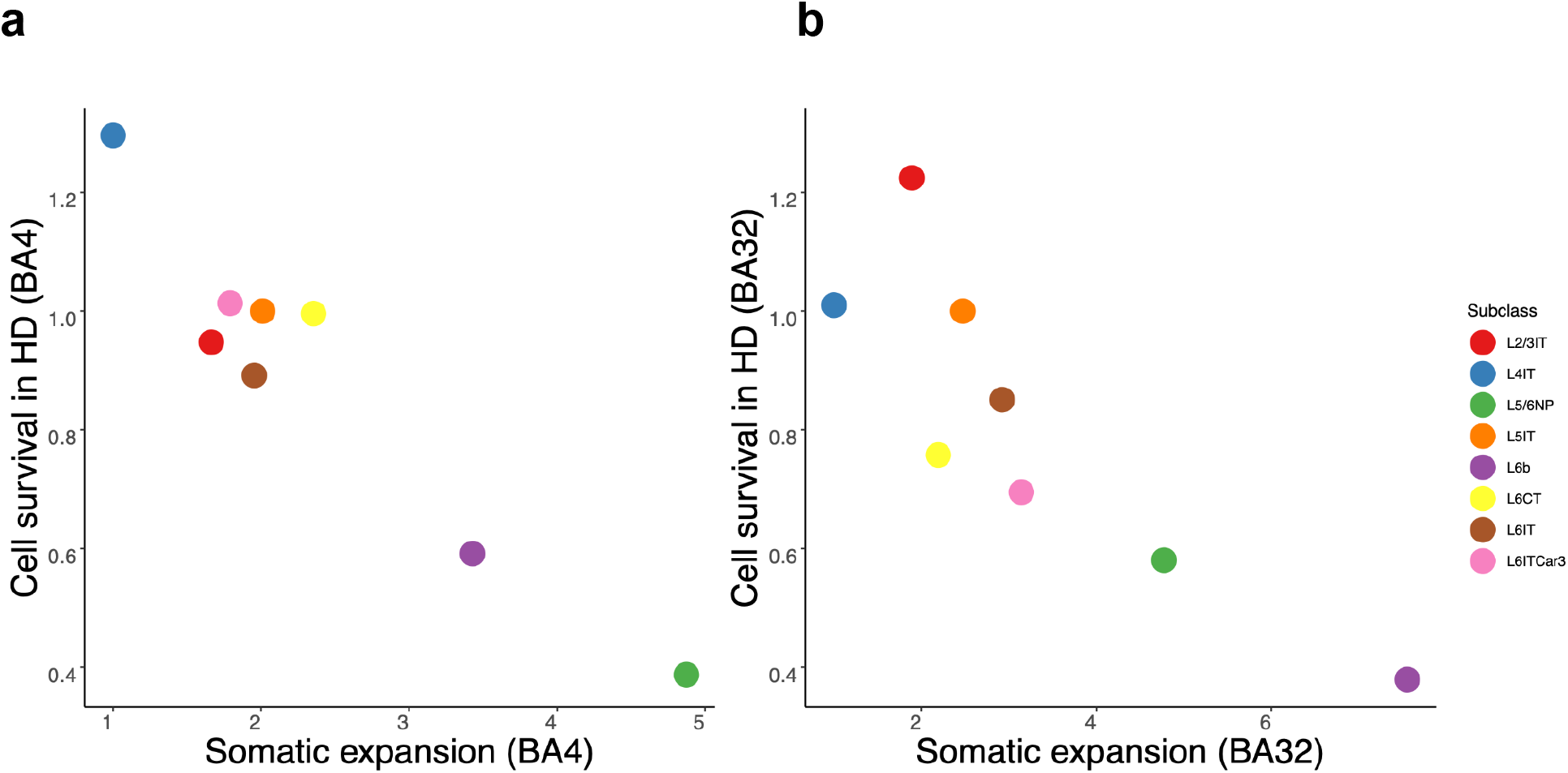
Loss of specific types of pyramidal neurons in HD, in two cortical areas (BA4 and BA32), in relation to these pyramidal types’ relative amounts of somatic expansion. Cell survival rates (y-axis) were calculated from the ratio of the group medians (median abundance in cortical samples among 50+ persons with HD relative to median abundance among 50+ unaffected control brain donors, where, for each donor, abundance was computed as a ratio of the number of neurons of that subtype to the number of L5IT neurons). Pyramidal subtypes’ relative amounts of somatic expansion (x-axis) were estimated from regression of the log of the CAG-repeat length changes of the individual pyramidal neurons (n = 20,872 from BA4 and n = 23,375 from BA32) against independent variables for the cell identities (pyramidal “subtypes”) (Methods). We used the resulting regression coefficients for the cell identities as estimates of the pyramidal subtypes’ relative propensities for somatic expansion (x-axis). These coefficients have been rescaled so that the value for the least expansive pyramidal subtype (L4IT) is 1.0. **Fig. S8** shows the replication of such estimates across brain donors. (**a**) Data from BA4. (**b**) Data from BA32.

**Figure 7.**
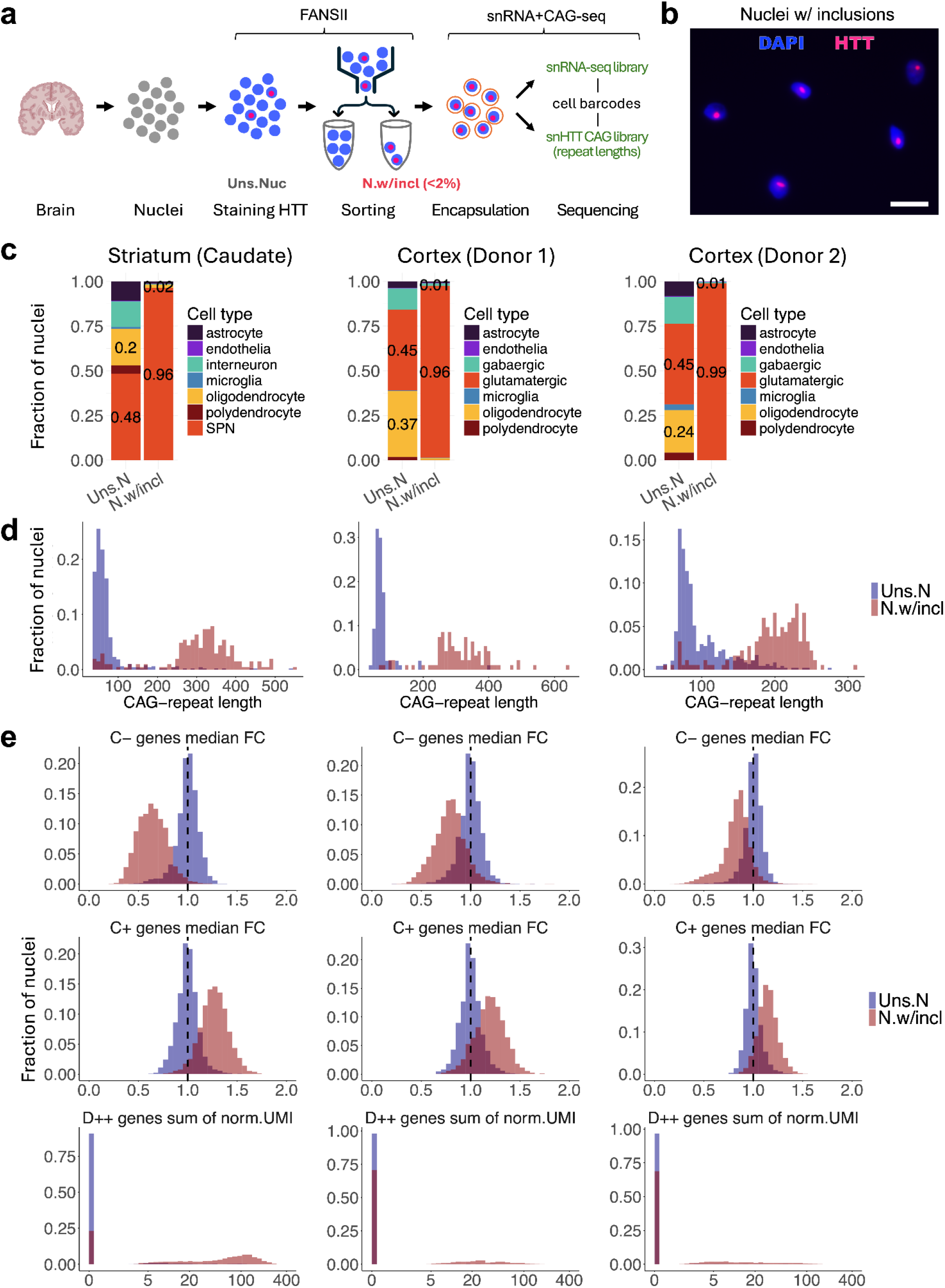
CAG-repeat expansions and phase C and D gene expression changes in nuclei with HTT inclusions. (**a**) Experimental approach for isolating nuclei with HTT inclusions by sorting. (**b**) Immunofluorescence microscopy analysis of nuclei from the sorted fraction (“nuclei with inclusions”, also labeled “N w/incl” in subsequent panels). Scale bar = 20 µm. (**c**) Cell types present among unsorted nuclei (“Uns N”) and nuclei in the sorted fraction “N w/ incl”. (**d**) CAG-repeat length distributions of SPNs (left) or glutamatergic neurons (middle and right) among unsorted nuclei (blue) and nuclei in the inclusion-sorted fractions (red). (**e**) Gene-expression changes in unsorted nuclei (blue histogram) and nuclei with inclusions (red histogram), for (from top to bottom) phase C- genes, phase C+ genes, and phase D++ genes. (Expression changes for C- and C+ genes are quantified in terms of median fold-change relative to baseline expression level; and for D++ genes, which are non-expressed or negligibly expressed at baseline, in terms of the total summed number of UMIs (per 100k) ascertained across these genes.) The black dashed line represents the baseline median fold change (x = 1) for C− and C+ genes. The fold-change calculations are described in Methods, and the genes used in the calculations are listed in Data S2. Additional analyses and visualizations are in **Fig. S15**.

Phase D also elicited (as in SPNs) greatly elevated expression of *CDKN2A* (more than 40-fold, **Fig. 5d**) and *CDKN2B* (8-fold), which promote senescence and apoptosis in many cellular contexts ^38–40^.

In adult mice, inactivation of the polycomb repressive complex 2 (PRC2) elicits (after several months) a similar set of de-repression events, including Cdkn2a, Cdkn2b, and many of the same transcription-factor genes; it also elicits progressive, fatal neurodegeneration ^41^. PRC has a well-established role in gene silencing in neurons. It is thus intriguing to consider the possibility that some aspect of polycomb function has been compromised in phase D neurons.

These results reveal shared principles for the gene-expression changes in striatal and cortical neurons in HD: phase C involves erosion of positive features of their respective identities, whereas phase D involves a later, more-rapid de-repression of embryonic development genes alongside increased expression of markers of imminent senescence or apoptosis. These functional themes are shared by affected cortical and striatal neurons with CAG-tract expansions beyond 150 CAGs.

### Neuronal loss (phase E) affects high-somatic-expansion pyramidal types

The variable somatic CAG-repeat expansion among different pyramidal neuron types (**Fig. 2,3**), the high repeat-length threshold for *HTT* toxicity in cortical neurons (∼150 CAGs, **Fig. 4a**), and our hypothesis that phase C/D transcriptional changes lead to neurodegeneration, together made a strong, testable prediction: across the types of pyramidal neurons, somatic expansion should have a strong negative relationship to neuronal survival in HD.

To estimate the survival rates of the various types of pyramidal neurons in HD, we analyzed two cortical areas – anterior cingulate cortex (BA32), and motor cortex (BA4) – in 100+ donors (including at least 50 persons with HD and 50 unaffected controls, with 95 donors in common between the two analyses). We then used the data to quantify loss of each type of neuron in each of the two cortical areas.

In both cortical areas, pyramidal neurons’ vulnerability to loss in HD was strongly related to their tendency to have expanded their CAG repeats somatically, with high-expansion types exhibiting much more severe loss than low-expansion types, and a clear relationship across the entire range (**Fig. 6**). L6b and L5/6NP neurons suffered particularly profound attrition, having declined by 40-60% in persons with HD relative to their abundance in unaffected controls (**Fig. 6**).

Given conflicting earlier reports^4–6,42,43^ about whether cortical interneurons suffer attrition in HD, we also analyzed the abundances of five populations of cortical interneurons (SST+, PVALB+, LAMP5+, VIP+, and SNCG+). Our analysis found no significant effect of HD on abundance of any of these interneuron types in either BA4 or BA32, despite the large number of donors in the analysis (50 with HD, 50 unaffected) (**Fig. S10a**). The same analyses detected substantial loss of SST+ and VIP+ interneurons with age, in both BA4 and BA32 – at similar rates in unaffected persons as in persons with HD (**Fig. S10b**) – an effect which causes their abundance to vary greatly from person to person. (This strong effect of age may, unless addressed in analyses, complicate case-control comparisons in HD and other disorders.)

Thus, in HD, the various types of cortical neurons appear to be vulnerable to neurodegeneration to an extent predicted by somatic expansion of their *HTT* CAG repeats.

### Non-cell-autonomous sequelae of neurodegeneration

Our earlier study of striatal neurons in HD ^31^ proposed that striatal pathology begins in those neurons that have expanded their CAG-repeat tracts beyond 150 CAGs – a model which the current results extend to the neocortex. Three studies ^7,8,44^ have challenged this model, based on analyses of case-vs-control “differentially expressed genes” (DEGs) whose expression levels are higher or lower (on average) in persons with HD than in unaffected persons. In particular, these studies report many DEGs that are not phase C or phase D genes, and that exhibit fold-changes sufficiently large as not to be readily explained by changes in the small fraction of neurons that have >150 CAGs at any moment in time ^7,8,44^ but have not yet been lost.

We used the data from our 100-donor motor cortex (BA4) analysis to further recognize and characterize case-vs-control DEGs. Analysis detected thousands of statistically significant DEGs in every cell type – as may be expected in a disease in which neurodegeneration has led to profound brain atrophy and declining health. Focusing on pyramidal neurons, these data also replicated the specific DEGs reported in the earlier work on human HD-impacted neocortex by Pressl et al. ^8^ (**Fig. S11**).

The key question is whether these case-vs-control DEGs are indeed cell-autonomous effects of the *HTT* CAG repeat – or, alternatively, are downstream effects of earlier neuronal loss and brain atrophy. The hypothesis that the DEGs are causes (rather than consequences) of neurodegeneration makes two predictions. The first prediction is that such changes would be specific to “vulnerable” neuronal types, i.e., to those cell types that are lost in HD. The data indicated that near-identical changes were in fact present in all pyramidal-neuron types, regardless of whether or not those types are lost in substantial numbers in HD (**Fig. S11, S12**). The second prediction is that, within any one person, such effects would be associated with the length of a neuron’s CAG-repeat tract. This prediction, too, found no support in the data: expression levels of the case-vs-control DEGs showed no relationship to CAG-repeat tract length across any part of the length range (**Fig. S13**). Thus, most case-vs-control DEG effects appear not to be related to cell-type-specific vulnerability nor to the length of a cell’s own CAG-repeat tract; such effects could instead be from system-wide responses to broad pathologies (such as atrophy, earlier neuronal loss, inflammation, or declining health).

The model we propose – that the pathology-initiating effects of the *HTT* CAG repeat arise in those sparse neurons with expansions beyond 150 CAGs – makes its own testable predictions. The first is that a neuronal type’s tendency to develop such long somatic expansions would predict its vulnerability to loss, a prediction confirmed above (**Fig. 6**).

Another prediction is that key pathological features specific to HD would arise in precisely those neurons with CAG-repeat tracts longer than 150 CAGs. We turned next to testing this prediction with a different kind of experiment.

### Nuclear inclusions of HTT protein

A long-noted finding of neuropathological studies of HD involves prominent inclusions of aggregated HTT protein, which are observed in persons with HD (but not in unaffected individuals)^45^ and in mouse models of HD (but not in wild-type mice)^46^.

CAG repeats encode polyglutamine (polyQ) tracts in proteins; long polyglutamine tracts can promote protein aggregation, leading to formation of inclusion bodies in HD and other

CAG-repeat expansion disorders, such as spinocerebellar ataxias (SCAs)^47^. The pathological significance of these inclusions has long been debated, with diverse views on whether inclusions are toxic, incidental, or even protective^48^. Interpretive uncertainty has been compounded by differences between endogenous and exogenous expression systems, full-length versus partial-length proteins, nuclear versus cytoplasmic inclusions, and human versus mouse models. A striking difference between human HD brains and knock-in mouse models has been central to this debate: HD mouse models (such as Q175) exhibit widespread inclusions but minimal neurodegeneration^49^, whereas human HD brains exhibit sparse inclusions but widespread neuronal loss^45,50^. It is natural that, in this context, the pathological significance of HTT inclusions has been uncertain.

We sought to recognize through experiments whether and how nuclear HTT inclusions in the human brain relate to (i) long CAG-repeat expansions and (ii) phase C and D gene-expression changes.

To do this, we developed a fluorescence-based sorting approach for isolating nuclei with HTT inclusions, FANSII (Fluorescence-Activated Nuclei Sorting with Intranuclear Inclusions). In this approach, nuclei are isolated from human post mortem tissue, stained with an antibody to the N-terminal region of HTT (with an epitope that has been mapped to the part of the HTT protein encoded by *HTT* exon 1, i.e. that is present on the HTT1a protein) ^50,51^, and then sorted based on their staining intensity (**Fig. 7a, Fig. S14a**). Isolating the small fraction of nuclei (generally 0-2% of all nuclei) with HTT inclusions made it possible to compare these sparse nuclei to the larger (unsorted) population of nuclei.

Immunofluorescence microscopy confirmed that 88-99% of the sorted nuclei contained visually prominent nuclear inclusions (**Fig. 7b, Fig. S14b**), in comparison to 0-2% of the unsorted nuclei.

We used this approach to sample, sort and prepare nuclei from the striatum and cortex of persons with HD, then analyzed the resulting populations of nuclei – unsorted nuclei, and nuclei with inclusions (88-99% confirmation rate by microscopy) – by our snRNA+CAG-seq approach. The genome-wide RNA data made it possible to associate each individual nucleus with its cell type and its transcriptional state.

To determine what cell population(s) harbor nuclear inclusions in HD, we analyzed RNA expression in the individual nuclei from the inclusion-positive nuclear fractions. In contrast to the diverse cell types represented among unsorted nuclei, almost all the inclusion-positive nuclei came from pyramidal neurons (in the cortex) and SPNs (in the striatum) (**Fig. 7c**) – the same types of neurons that degenerate in large numbers in HD. Among pyramidal neurons, the most slow-expanding type (L4IT) was under-represented (by 7- to 11-fold) among nuclei with inclusions (relative to unsorted neuronal nuclei), while fast- and moderately-expanding types (L6b, L6CT, L6IT) were over-represented (by approximately 2- to 8-fold) (**Fig. S15**); L5/6NP neurons were extremely sparse (<1% and <0.1% of the total in the two cortex donors) in both fractions, reflecting what is presumably decades of earlier loss (**Fig. S15**).

In every sample (cortex and striatum), unsorted nuclei and inclusion-positive nuclei exhibited dramatically different distributions of CAG-repeat lengths (**Fig. 7d**). In each experiment, the median CAG-repeat lengths among the inclusion-positive nuclei were >200 CAGs, a length obtained by less than 1% of the unsorted nuclei (**Fig. 7d**). These results suggest that the sparse nuclei with HTT inclusions and the sparse nuclei with long CAG-repeat expansions are the same nuclei.

The nuclei in the inclusion-positive fractions exhibited advanced phase C gene expression changes; the median (across inclusion-positive nuclei) of the median fold change (across phase C- genes) corresponded to the lower 5% tail of the distribution for the unsorted neuronal nuclei (**Fig. 7e**). A substantial fraction of these nuclei (>75% in striatum, >25% in cortex) also exhibited phase D de-repression changes(**Fig. 7e, Fig. S16abc**) – whereas only about 2% of the unsorted cortical nuclei exhibited such changes (**Fig. 7e**). Phase C and phase D gene-expression changes were the primary changes that distinguished nuclei with inclusions from unsorted neuronal nuclei (**Fig. S16def**).

These results associate nuclear inclusions, ultra-long CAG-repeat expansions, and gene-expression pathologies with the same sparse set of neuronal nuclei in human HD brain, and connect the formation of nuclear inclusions – in both cortex and striatum – with a late stage of the toxicity (advanced phase C and phase D) that arises among neurons with CAG-repeat expansions beyond 150 CAGs.

## Discussion

These results suggest that a single, unifying model explains the distinct HD pathologies that arise in two key brain areas, the striatum and the neocortex. They also associate nuclear HTT inclusions with the final stages (late phase C and phase D) of the pathological trajectories of individual neurons.

We found that cortical neurons in HD traverse a sequence of pathogenic phases analogous to those traversed by striatal neurons – but do so with highly variable kinetics and distinct biological sequelae. These phases involve decades of slow somatic expansion of the disease-causing CAG repeat tract (phase A); much-faster subsequent expansion (phase B); pervasive gene-expression changes that commence at about 150 CAGs, eroding the positive features of their respective identities (phase C); a de-repression crisis (phase D); and then elimination (phase E). Curiously, though, the specific gene-expression changes that arise in phase C (and thus distort the biology of these neurons while they are still living, and elicit downstream effects in tissues) are quite different in cortical and striatal neurons – potentially helping explain how the pathologies in these brain areas could have appeared to be so different.

We found that nuclear HTT inclusions – a historically hard-to-interpret feature of HD – appear in the late pathological phases (advanced phase C and phase D), in the same sparse neurons that also have extremely long somatic CAG-repeat expansions, in both striatum and cortex. One longstanding question about such inclusions has been why they are so sparse in HD-affected human brain tissue, yet appear in most-to-all cells (including glial cells) in the mouse Q175 model of HD. Our results suggest an explanation: in humans, nuclear inclusions appear only in specific cells – neurons that have acquired long somatic CAG-repeat expansions – in a toxicity phase through which such cells pass asynchronously and relatively quickly. In contrast, in the Q175 mouse, all cells begin life above the toxicity threshold, exposing cells of all types to synchronous toxicity and inclusion formation.

Our findings suggest that large nuclear HTT inclusions can serve as a cytological marker of neurons that have entered late phases of the CAG-repeat-driven toxicity. Whether nuclear inclusions are the direct cause of this toxicity remains to be determined.

These results offer simple, unifying explanations for the long-noted differences in the timing, severity, and gene-expression changes in HD pathologies in cortex vs. striatum. Differences in timing and severity appear to arise primarily from differing rates of somatic CAG-repeat expansion: only Layer 6b cortical pyramidal neurons appeared to acquire long somatic

CAG-repeat expansions to an extent comparable to SPNs, with other pyramidal types exhibiting much less expansion. The long-noted differences between striatal and cortical gene-expression changes in HD result in part from an intriguing phenomenon. Upon reaching the high threshold for toxicity (of about 150 CAGs) and entering phase C, cortical and striatal projection neurons undergo almost completely different gene expression changes (**Fig. 4d**) – changes that thus likely have different physiological sequelae and different downstream effects on other cells.

However, the striatal and cortical phase C changes share a surprising property: the genes that decline in expression are genes whose expression defines the identity of the affected neuronal type (**Fig. 4e**). An important focus of future work will be to identify a cellular mechanism for such progressive losses of neuronal identity.

The possibility that corticostriatal projections are fundamental to mechanism in HD – with repeat-length-dependent pathology in one brain area eliciting secondary pathology in the other – has long intrigued neuroscience, leading to ideas that the pathophysiological mechanism might involve retrograde spreading of aggregation-driving prions ^15,16^, excitotoxicity ^10–12^, or loss of synaptic connections (corticostriatal disconnection) ^8,13^. We believe that our results warrant reconsideration of this longstanding conception of HD pathology. First, our results suggest that the same fundamental, cell-autonomous dynamic propels degeneration of cortical and striatal neurons in HD – a dynamic in which expansion of each neuron’s own CAG-repeat tract beyond a threshold of about 150 CAGs initiates a pathological sequence of events. Second, we find that cortical pyramidal neurons of all types – regardless of molecular identity, connectivity, laminar location, or cortical area – are vulnerable to phase C pathology in those individual neurons whose CAG-repeat tracts have expanded beyond the 150-CAG threshold (**Fig. 4a**), and that the various types of pyramidal neurons suffer attrition in HD in a way that is predicted by their somatic expansion rates (**Fig. 6**). The appearance of “resilience” or “vulnerability” in specific cortical cell types (or layers) thus arises from variation in their average rates of somatic CAG-repeat expansion, which in turn determine what fraction of cells reach and cross the toxicity threshold during a human lifetime.

Single-cell genomics has been creating ever-more-elaborate taxonomies of pyramidal neurons based on their RNA expression patterns ^32,33^. However, the functional significance of these distinctions has been uncertain, as physiological differences among most of these neuronal types remain to be identified ^36^. Here we identified an important functional difference in HD: dramatic differences in their somatic DNA-repeat expansion (**Fig. 3**).

We propose that the apparent resilience (and vulnerability) of neuronal populations of various types comes not from connectivity to the striatum, nor from differential sensitivity to HTT toxicity, but instead from whether they have low (or high) rates of somatic DNA-repeat expansion – which determines whether a small (or large) fraction of neurons reach the biologically consequential threshold of about 150 CAGs during a human lifetime (**Fig. 4a, 6**). In principle this could also be generally true of other kinds of neurons throughout the brain.

Our results suggest that cortical neurons – even more than striatal neurons – may spend the longest period of their lives (perhaps 95-100%) with an innocuous *HTT* gene, then experience HTT toxicity in staggered temporal windows. Such a model suggests that in the neocortex, as in the striatum, there might be great therapeutic leverage in slowing somatic DNA-repeat expansion as a way to prevent HD in persons who have inherited HD-causing alleles. These results also suggest that a therapy that successfully slowed or stopped somatic DNA-repeat expansion would postpone or prevent neuronal loss in the neocortex as well as the striatum.

### Limitations of the study

Our analyses have focused on the cell-autonomous effects of the CAG repeat on a cell’s own biology as measured through its gene expression. Downstream of these effects, there are certain to be vast additional changes in the HD-transformed brain as this brain tissue responds to the effects of neuronal loss, atrophy, inflammation, circuit disconnection, and changed health circumstances. It is likely that many HD symptoms arise from these downstream sequelae rather than from the primary neurodegenerative events; making sense of this is an important direction for future work.

Our findings suggest that large nuclear HTT inclusions are present in the same sparse neurons that have entered late phases of the cell-autonomous toxicity process – but whether HTT aggregation directly causes this toxicity remains to be determined.

## Supporting information

Data S1 (supplemental data set)

Data S2 (supplemental data set)

Data S3 (supplemental data set)

## Acknowledgements

We are grateful to the patients and families whose donations of brain tissue enabled this work, and to the NIH NeuroBioBank for providing this tissue. This work was funded by CHDI Foundation, Inc., a nonprofit biomedical research organization exclusively dedicated to developing therapies that will substantially improve the lives of HD-affected individuals; the Ludwig Neurodegenerative Disease Seed Grants Program at Harvard Medical School; the Harvard Medical School Department of Genetics; and the National Human Genome Research Institute of the National Institutes of Health under award R01HG006855. We are grateful to Vahri Beaumont, Thomas Vogt, Jian Chen, and Cristina Sampaio for helpful advice and suggestions throughout this project’s conception and execution; and to Marta Florio, Nolan Kamitaki, Chris Patil, Vanessa Wheeler, Marcy MacDonald, James Gusella, Philip Reaper, and the people above for comments on manuscript drafts.

## Declaration of interests

The authors declare no competing interests.

## Resource availability

### Lead contact

Requests for further information and resources should be directed to and will be fulfilled by Steven McCarroll.

## Materials availability

Brain tissue samples from the donors in this study can be obtained by application to the NIH NeuroBioBank (https://neurobiobank.nih.gov/).

## Data availability

Datasets of a size suitable to be included as tabs of a spreadsheet (for example, the results of all the single-gene regression models identifying phase C and phase D genes) are included in Supplementary Data.

Raw datasets including snRNA-seq counts matrices, donor metadata, and raw sequencing data (including raw sequencing reads (FASTQ) and alignment files (BAM)) associated with this study, have been deposited to the NIH Neuroscience Multi-Omic Archive (NeMO; https://nemoarchive.org) under accession nemo:col-ei2o5ud. All non-identifying data sets are in the open-access part of the repository. Human data that are individually identifying (e.g. raw sequence reads containing brain donors’ allelic information) are available through NeMO’s controlled-access mechanism, for general research use.

Processed datasets and R scripts used to generate the figures have been deposited to Zenodo.

## Methods

### Experimental model and study participant details

We analyzed brain tissues from the Harvard Brain Tissue Resource Center/NIH NeuroBioBank (HBTRC/NBB), which were obtained by application (request #2496, “Neocortical cellular changes in Huntington’s Disease”).

Brain donors were recruited by the HBTRC/NBB in a community-based manner across the USA. The HBTRC procedures for informed consent by the donor’s legal next-of-kin and distribution of de-identified post-mortem tissue samples and demographic and clinical data for research purposes are approved by the Mass General Brigham Institutional Review Board. Post-mortem tissue collection followed the provisions of the United States Uniform Anatomical Gift Act of 2006 described in the California Health and Safety Code section 7150 and other applicable state and federal laws and regulations. Human brain tissue was obtained (for data generation) from the HBTRC/NBB (NBB request ID# 1835). Federal regulation 45 CFR 46 and associated guidance indicates that the generation of data from de-identified post-mortem specimens does not constitute human participant research that requires institutional review board review.

The HBTRC/NBB confirmed HD diagnosis and excluded clinical comorbidity and presence of unrelated pathological findings by reviewing medical records and by formal neuropathological assessment. The 1985 Vonsattel et al. grading of neostriatal pathology^54^ was used for diagnosis. Diagnosis on early cases is done using histological stainings and polyglutamine immunohistochemistry ^54–57^. Positivity in pontine gray neurons rules out HD-like-2 neuropathology ^58^, and cerebellar dentate neurons are mildly positive even in very early cases, while Purkinje cells are negative (unlike in cerebellar ataxia CAG expansion cases).

## Method details

### Isolation of nuclei from brain tissue

Nuclei were isolated from frozen brain tissue using a density-gradient method ^31^. Briefly, frozen tissue was homogenized in Nuclei EZ Lysis Buffer (MilliporeSigma, #NUC101) supplemented with NxGen® RNase Inhibitor (1 U/mL; Biosearch Technologies, #30281) using Dounce homogenizers. Homogenates were filtered through 70 µm cell strainers (Bel-Art, #H13680-0070) and centrifuged at 500 × g for 5 min at 4 °C. Supernatants were discarded and pellets were resuspended in 300 µL of G30 buffer (30% iodixanol, 3.4% sucrose, 20 mM tricine, 25 mM KCl, 5 mM MgCl₂; pH 7.8). An additional 1 mL of G30 was gently underlaid to form a cushion, and the sample was layered above it. After centrifugation at 8,000 × g for 10 min at 4 °C, supernatants were removed. Nuclei pellets were washed once in 1 mL wash buffer (1% BSA in 1× PBS with 1 U/mL NxGen RNase Inhibitor), followed by 500 × g for 5 min at 4 °C. Nuclei were resuspended in 50 µL wash buffer and counted on a LUNA-FL Dual Fluorescence Cell Counter (Logos Biosystems).

### Preparation of snRNA-seq libraries

Nuclei encapsulation and library preparation were performed with Chromium Next GEM Single Cell 3′ Reagent Kits v3.1 (10X Genomics, PN-1000121) or Chromium GEM-X Single Cell 3′ Reagent Kits v4 (10X Genomics, PN-1000691) following the manufacturer’s protocol, with minor modifications during cDNA amplification (see Single-cell measurement of CAG-repeat length in snRNA-seq experiments below). Libraries were sequenced on Illumina NovaSeq 2000, 6000 or NovaSeq X platforms.

### Single-cell measurement of CAG-repeat length in snRNA-seq experiments

We previously developed an approach to simultaneously measure HTT CAG-repeat length and transcriptomes in individual nuclei ^31^. In brief, a whole-transcriptome library and an HTT-CAG targeted library were generated and linked by shared 10X cell barcodes.

During the cDNA amplification step (step 2.2a of the standard 3′ scRNA-seq protocol from 10X), we added two spike-in primers targeting the 5′ region upstream of the HTT CAG repeat to 1 µM each:

Spike-in A: 5′-CCCAGAGCCCCATTCATTGCC-3′

Spike-in B: 5′-GGCGACCCTGGAAAAGCTGATG-3′

In step 2.2d, the extension time was increased to 2 min. The amplified cDNA was split to generate (1) the conventional snRNA-seq library (10 µL) by continuing the standard 10X workflow and (2) an HTT-CAG library (4 µL) by targeted PCR using the UltraRun LongRange PCR Kit (QIAGEN, #206444; total reaction volume 20 µL) with following primers and PCR program:

Biotin IllumR-HTT-C:

5′-/5Bio/GTCTCGTGGGCTCGGAGATGTGTATAAGAGACAGCCTTCGAGTCCCTCAAGTCCTTC-3′

Illum_f_10X_3p_B:

5′-TCGTCGGCAGCGTCAGATGTGTATAAGAGACAGCTACACGACGCTCTTCCGATCT-3′

PCR program: 93 °C 3 min; 18–19 cycles of 93 °C 30 s, 60 °C 30 s, 68 °C 8 min; 72 °C 10 min; hold 4 °C.

The amplified HTT-CAG library was size-fractionated with SPRI beads (Beckman Coulter, #B23319) using 0.4X (Long, “L”) and 1.0X (Short, “S”) selections. DNA was eluted in 10 µL water. “L” and “S” fractions (either both or “L” only) were mixed with 10 µL of Dynabeads MyOne Streptavidin C1 (Thermo Fisher Scientific, #65002) pre-washed in 2X Wash & Binding buffer (2 M NaCl, 1 mM EDTA, 10 mM Tris-HCl, pH 7.5), incubated for 30 min at RT with rotation, washed 3 times in 1X Wash & Binding buffer, and resuspended in 10 µL water.

Bead-bound templates were then amplified for 12–14 cycles with unique pairs of standard Nextera i7/i5 indexing primers (the same index pair was used for “L” and “S” libraries when both were sequenced together) using the same kit and cycling conditions (total PCR volume 40 µL). PCR products were purified by 1.0X SPRI (eluted in 10 µL) and prepared for long-read sequencing.

For PacBio libraries, multiple samples were pooled equimolarly and prepared using SMRTbell Express Template Prep Kit 2.0 (Pacific Biosciences, #100-938-900) following PN 101-892-000 v01 (Jan 2020). “L” and “S” libraries were loaded on different flow cells and sequenced on Sequel IIe with a 100 pM loading concentration.

For Oxford Nanopore libraries, unpooled samples were barcoded and pooled with Native Barcoding Kit 24 V14 (ONT, SQK-NBD114.24; protocol V_NBA_9168_v114_revR_30Jan2025). The pooled libraries (35–50 fmol) were loaded on FLO-PRO114M flow cells and sequenced on PromethION 2 Integrated (P2i) instruments for 72 h with Super-accurate (SUP) base-calling.

### FANSII (fluorescent-activated nuclei sorting with intranuclear inclusions)

Freshly isolated nuclei from frozen human brain tissue were incubated overnight at 4 °C with an anti-HTT antibody (mEM48; MilliporeSigma, #MAB5374; 1:250) at a density of 10,000 nuclei/µL in PBSAi buffer (3% BSA in 1× PBS supplemented with 0.5 U/mL NxGen RNase Inhibitor), supplemented with 0.025% Tween-20 and 0.2× protease inhibitor cocktail (Thermo Fisher Scientific, #78429).

The following day, nuclei were pelleted by centrifugation (4 °C, 500 × g, 5 min), and the supernatant was removed. Nuclei were washed three times with PBSAi buffer by gentle resuspension in twice the original staining volume, followed by centrifugation and removal of the supernatant.

Nuclei were then incubated for 1 h at 4 °C with a fluorophore-conjugated secondary antibody (Thermo Fisher Scientific, #A-21240; 1:1000) in PBSAi buffer supplemented with 0.025% Tween-20 and 0.2× protease inhibitor cocktail, maintaining the same nuclear density used for primary antibody staining. Following incubation, nuclei were pelleted (4 °C, 500 × g, 5 min), the supernatant was removed, and nuclei were washed once with PBSAi buffer at 4 °C.

To label residual endoplasmic reticulum contiguous with the nuclear membrane, nuclei were incubated with ER-Tracker Green (Thermo Fisher Scientific, #E34251; final concentration 1 µM) in PBSAi buffer at room temperature for 15 min. After an additional wash in PBSAi buffer at room temperature for 10 min, nuclei were resuspended in DAPI (0.5 µg/mL), filtered through a 35 µm cell-strainer cap into 5 mL FACS tubes (Fisher Scientific, #08-771-23), and sorted using a BD FACSAria™ cell sorter equipped with a 70-µm nozzle.

Nuclei containing HTT inclusions were gated as HTT-high and ER-Tracker-low to minimize potential autofluorescence and non-specific fluorophore binding associated with residual endoplasmic reticulum content (see **Fig. S14**).

### Fluorescence imaging of sorted nuclei with HTT inclusions

Sorted nuclei (1.5 µL) were mixed with an equal volume of mounting medium (VECTASHIELD® HardSet™ Antifade Mounting Medium; Vector Laboratories, #H-1400-10). A volume of 2.5 µL of the mixture was mounted onto a glass slide and covered with a 12-mm round coverslip (Carolina Biological Supply, #633029). Samples were imaged within 10 min of mounting using a Zeiss Observer.D1 fluorescence microscope equipped with a Zeiss Axiocam HRm camera.

### Quantification and statistical analysis

#### Analysis of HTT-CAG library data

After sequencing, basecalling and alignment of the PacBio libraries was performed as previously reported ^31^. Alignment was performed using “pbmm2” either version 1.4.0 or version 1.10.0 with —preset ISOSEQ and using the GRCh38 “no alt” reference genome (ftp://ftp.ncbi.nlm.nih.gov/genomes/all/GCA/000/001/405/GCA_000001405.15_GRCh38/seqs_for_alignment_pipelines.ucsc_ids/GCA_000001405.15_GRCh38_no_alt_analysis_set.fna.gz).

For the Oxford Nanopore data, demultiplexing by barcode and read alignment was performed on the sequencing instrument using minknow version 6.4.8 and the same reference genome used for the PacBio sequencing.

After alignment, we further processed the long-read data from each flowcell using a custom analysis pipeline we developed (https://github.com/broadinstitute/HTT-CAG-Software). For the PacBio data, we used version 1.0 of this pipeline, the same version used in ^31^. For the Oxford Nanopore data, we used version 1.1 of the pipeline, which had only a few changes compared to the earlier version. The two principal differences are (a) we increased the search window for recognizing landmark sequences during read decoding and (b) we re-implemented parts of the consensus algorithms to improve efficiency and consume less memory to accommodate deeper sequencing (more reads per UMI).

#### Mapping of snRNA-seq profiles to cortical cell taxonomies

We sought to classify each of the hundreds of thousands of neuronal gene-expression profiles (from snRNA-seq experiments) into multiple cortical taxonomies, including the Siletti taxonomy (the most detailed and current taxonomy, as of this work), and the Hodge/Tasic taxonomy (to which data on axonal projections and laminar locations have been mapped by earlier studies).

In this project, we generated 10x snRNA-seq data in either village or single donor experiments. To all of these nuclei, we applied an scPred model that classifies each nucleus as one of 7 main cell types as well as the MapMyCells algorithm that assigns each nucleus to a subcluster_name / cluster_name / supercluster_name (see the Siletti taxonomy).

To analyze glutamatergic and GABAergic neurons, we selected the nuclei that were classified as either “glutamatergic” or “GABAergic” by scPred (with the assignment probability max.prob > 0.8) and also were assigned by MapMyCells to one of 4 excitatory or 3 inhibitory neurons’ superclusters (in the Siletti taxonomy).

For these glutamatergic and GABAergic sets of neurons, we further ran the corresponding scPred glutamatergic or GABAergic subclass / subtype models that were trained on the data from Hodge et. al

#### Mapping MapMyCells cluster_name to glutamatergic subclasses

To map each MapMyCells “cluster_name” (which in the Siletti taxonomy are children of 4 excitatory neuron superclusters ) to scPred’s glutamatergic subclasses, we used the village data from BA4 and (separately) BA32 to build confusion matrices between “cluster_name” and glutamatergic subclass assignments. For each cluster_name with at least 10 neurons in it, we computed subclass frequencies for the neurons labelled with this cluster_name and then assigned this cluster_name to the highest frequency subclass. We determined that for each of the cluster_names more than 60% (most often, more than 90%) of all the neurons belonged to a single glutamatergic subclass, to which we then associated it. We performed this mapping independently for the BA4 and BA32 data and confirmed that the mapping results derived from these two analyses/regions were identical.

#### Negative binomial regression analyses of somatic CAG-repeat expansion

For a cell *c* with an HD-causing allele length measurement, we define its phase A somatic expansion value, *phaseA_expansion*, as the difference between its CAG-repeat length (capped at 100 CAGs) in c and the inherited CAG-repeat length for the donor from whom cell *c* was sampled. In more formal terms,

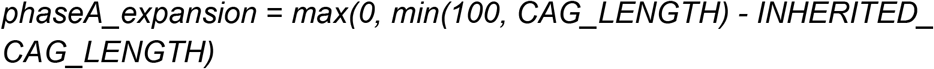

The reason for capping *CAG_LENGTH* at 100 is that at about this CAG-repeat length there is a transition from phase A to a far-faster phase B, and the CAG-repeat expansion dynamic accelerates significantly, and could potentially be governed by different cell-type-specific dynamics. So we focus on phase A expansion kinetics.

We model the phase A CAG-repeat expansion using a Negative binomial regression of this form:

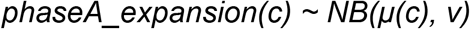

where the log mean of this negative binomial model, log(*μ*), is a linear function *L* of cell level covariates for a cell *c*. *L* is defined with one of these two formulas, depending on which subset of the data the model is fitted on.

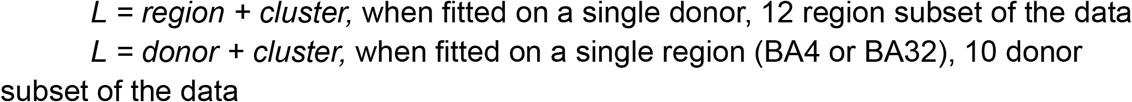

***region*** is the brain region (cortical area) from which cell *c* was sampled.

***donor*** is the donor from whom cell *c* was sampled. Donor-level effects implicitly include any effects of age, genetic background (e.g. modifiers), and disease progression.

***cluster*** is the cluster (in the Siletti taxonomy) to which cell *c* was assigned by MapMyCells.

**Fig. 3** reports values of the *region* and *cluster* coefficients, for 44 clusters with 10 or more neurons in at least one of 12 regions.

#### Negative binomial regression analyses of gene-expression changes

In order to identify genes whose expression changes as the HD-causing *HTT* allele’s

CAG-repeat tract expands, in our recent study we fitted per-gene Negative Binomial Regression models that used various covariate selections as well as their functional forms ^31^.

In the current study, we adopted the same approach to modeling gene expression as a function of CAG-repeat length (CAG_LENGTH) and additional covariates, with a slight modification of the covariates used in the current model. The NBR models for gene expression in the cortex are as follows:

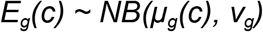

where the log mean of this negative binomial model, log(*μ_g_*), is a linear function *L* of the following cell level covariates for a cell *c*:

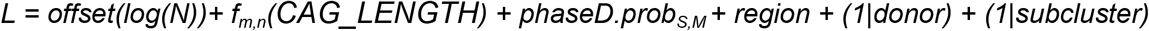

***N*** is the total count of UMIs (RNA transcripts from any gene) in cell *c*

***f_m,n_(CAG_LENGTH)*** is a function family (parameterized by positive integers *m* and *n such that m < n*) of the CAG-repeat length of the HD allele in cell *c*. *f* can be a simple function of CAG_LENGTH (for example, it can be CAG_LENGTH itself), or it can be defined, for example, as a “hinge function” (in which CAG_LENGTH has no effect until a threshold m is reached, then has a continuously escalating effect with further increase in CAG_LENGTH):

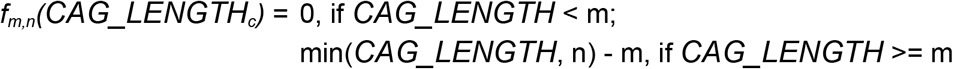

***phaseD.prob_S,M_*** is a covariate used to measure the probability of cell c having entered phase D. It is modeled as the following logistic function of the total normalized count of the phase D++ gene transcripts (computed as the sum of the UMI counts normalized to 100k over all 59 phase D++ genes):

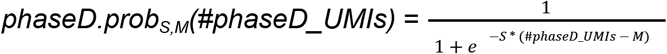

where S and M are the steepness and the midpoint of the function respectively.

***region*** is the brain region (cortical area) from which cell *c* was sampled.

***donor*** is the donor from whom cell *c* was sampled. Donor-level effects implicitly include any effects of age, genetic background (e.g. modifiers), and disease progression. ***subcluster*** is the subcluster (in the Siletti taxonomy) to which cell *c* was assigned.

Note that *log(N)* was modeled as an offset, *CAG_LENGTH, phaseD.prob* and *region* as fixed effects covariates, and *donor* and *subcluster* as random effects covariates.

The R function we used to fit the per-gene Negative Binomial Regression models (*glmmTMB* from the eponymous R package) is very slow when run on a data set of this size and a model definition containing a random effects covariate (in this case, it’s *subcluster*) with more than 150 levels. Preliminary analyses revealed that essentially all of the highly significant results were arising from the small fraction of cells with repeat-tracts longer than 150. To make the recognition of phase C and phase D genes more computationally tractable as the data set expanded, we downsampled (to 25%) the data in the interval of (35-100] CAG repeats, while retaining all the cells with CAG-repeat length > 100. All NBR models were thus fitted to data from 16,793 pyramidal neurons, of which 13,381 (or 80%) had CAG-repeat length <= 100 and 3,412 (or 20%) had CAG-repeat length > 100.

To compare gene expression changes across cortical layers and areas, we also fitted per-gene NBR models on four data subsets: 12,034 upper-layer, 4,867 deep-layer, 4,273 BA4, and 3,359 BA32 neurons. These comparisons confirmed that gene expression changes associated with increasing CAG-repeat length are highly correlated both between upper- and deep-layer neurons, and between BA4 and BA32 neurons.The statistical significance values of the *f(CAG_LENGTH)* and *phaseD.prob* model terms were used to recognize (respectively) phase C and phase D genes, which are enumerated in **Supplementary Data.** The Supplementary Data spreadsheet contains the per-gene statistics for all genes, from both the regression models fitted on 16,793 pyramidal neurons as well as the models fitted on four subsets of the data (described before), all stored as the sheets named “cortex”, “cortex_UL”, “cortex_DL”, “BA4” and “BA32” respectively.

#### Quantification of phase C changes at the single-cell level

Expression changes for C+ and C- genes in individual cells were quantified in terms of their fold-change relative to baseline expression level: i.e. relative to the average expression level in the cells of the same pyramidal subtype which had CAG-repeat tracts shorter than 100 CAGs – generally, the large majority of the cells of that subtype.

To visualize the phase C+ or phase C- changes in individual cells, we calculate each individual cell’s “median fold-change” across these genes. Because snRNA-seq data can be sparse, in order for a fold-change to be well-estimated at the individual-cell level, we generally need to focus on the subset of phase C genes that are most highly expressed, in the sense that they are detected robustly across the vast majority of the individual cells. While using more phase C- and C+ genes in the median fold-change calculations may reduce the noise and make underlying relationships more clear, including genes with too-low expression levels and thus unstable fold-change estimates will result in the noisier median fold-change estimates as well.

We found that an inclusion threshold in which a gene must be detected in at least 90% of the cells in the analysis appeared to optimize the stability of the results and the clarity of the underlying relationships. The number of genes that reach this threshold can vary from experiment to experiment, depending mainly on the RNA quality of individual post mortem specimens (a higher per-nucleus UMI ascertainment allows a larger number of genes to reach this threshold and contribute to the calculation of the median fold-change). So, although the same high-expression phase C genes are included in all analyses, a variable number of medium-expression-level genes are included in each. A Supplementary Data Set lists the genes included in each individual figure panel.

#### Quantification of phase D changes at the single-cell level

For phase D++ (de-repressed) genes, which are negligibly expressed at baseline, we cannot effectively use a fold-change metric, since the denominator (expression level at baseline) is zero or extremely low, making the ratio infinite or overly sensitive to single-UMI changes. So, since the UMI counts for these genes are so low, we instead sum the UMI counts (in a single cell) across all the D++ genes, then divide by the total number of UMIs (across all genes) ascertained in that cell, and then rescale (multiplying this small decimal number by 100,000) so that the units are “UMIs per 100k”; this normalization corrects for the way that some cells have more UMI ascertainment than other cells do. (We use 100k because it is the power of 10 that is closest to the average number of UMIs ascertained per pyramidal neuron nucleus.) We used the same 59 D++ genes in all analyses.

#### Analyses of cell loss

For snRNA-seq analysis of pyramidal neuron loss across 100 donors, we processed sets (“villages”) of 20 brain donors (including HD and control donors) en bloc as single pooled samples, in a manner we previously described ^31,59^. Each 20-donor village was pooled and handled together from nuclei extraction through sequencing, and each village was partitioned across eight encapsulation reactions. Donor-of-origin for each nucleus was assigned computationally from combinations of transcribed SNPs detected in the nucleus’s RNA-seq reads ^60^.

We first normalize the abundance of each pyramidal neuron type relative to that of layer 5 intra-telencephalic-projecting (L5IT) neurons, a high-abundance pyramidal population that undergoes just modest amounts of somatic expansion. We found quantification in this manner to yield estimates that were more consistent (less variable) across unaffected control donors – superior in this regard to simply quantifying abundance as a fraction of all nuclei sampled – potentially because numbers of certain other populations (especially OPCs and interneurons) change with age.

These quantifications were made in each donor individually. For each pyramidal type, we then calculated the ratio of (median abundance among persons with HD) to (median abundance among unaffected donors).

#### Case-control DEG analysis

We identified case-vs-control differentially expressed genes (DEGs) in the 50+ brain donors with HD to the 50+ unaffected control donors, utilizing the BA4 data set and identifying DEGs using the R library DESeq2. To select the genes for **Fig. S11**, we compared to the data from the Pressl et al. study (whose Table S3 contains ATAC-filtered significantly differentially expressed genes for HD vs. NC for cell types they described as L5a, L6a, and L6b) and focused on genes that reached significance in both studies, without regard to direction of effect. We identified all the genes that were significant in at least one of our subclass DEG analyses (for **Fig. S12**, we further filtered these genes to be in the gene set “GOCC_SYNAPSE” from the Gene Ontology Consortium), and were also present among the genes listed in the Pressl DEG analysis for L5a pyramidal neurons.

---

**Table S1.**
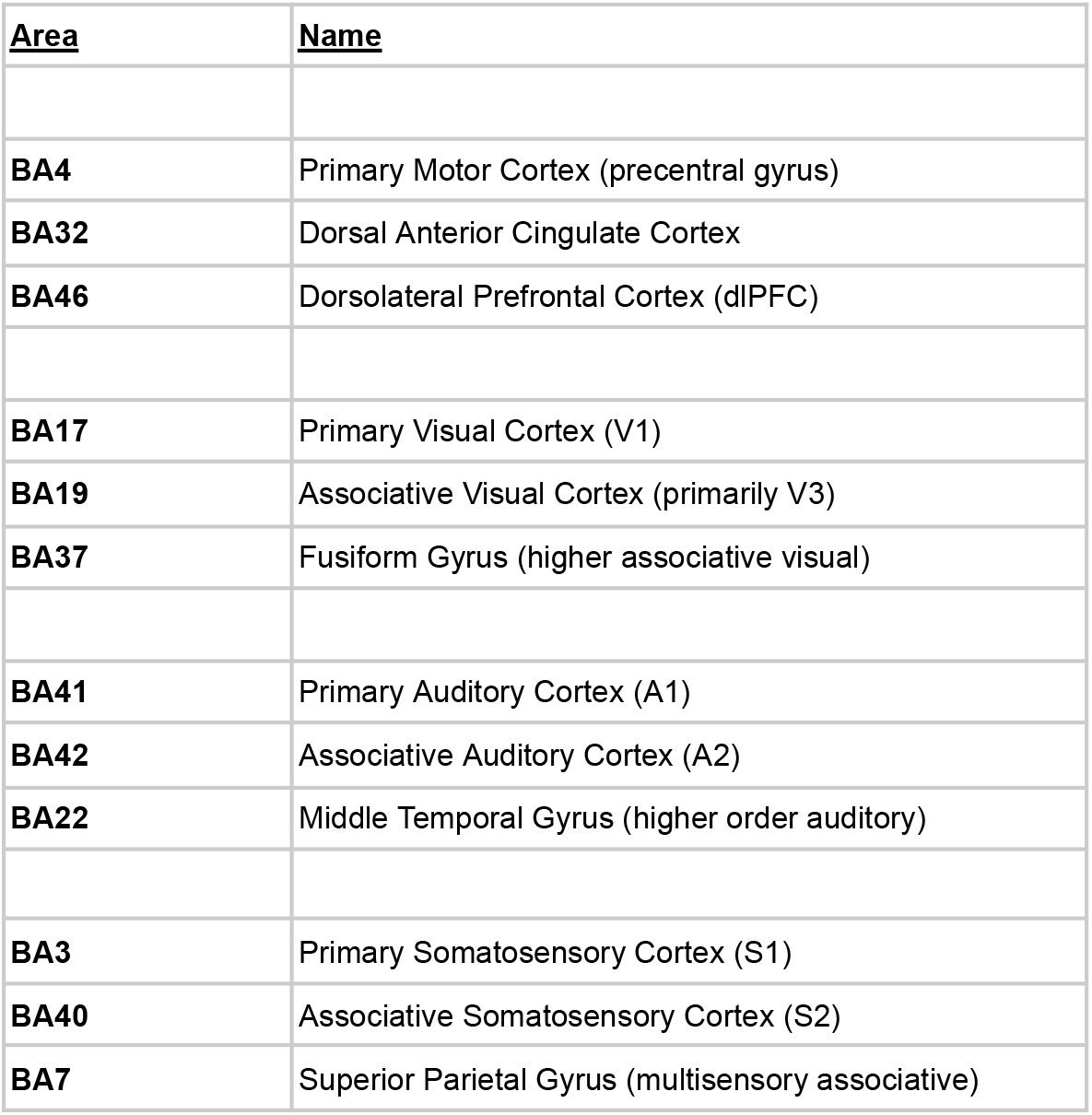
Cortical areas analyzed by snRNA-seq. Related to Figures 2 and 3.

**Figure S1.**
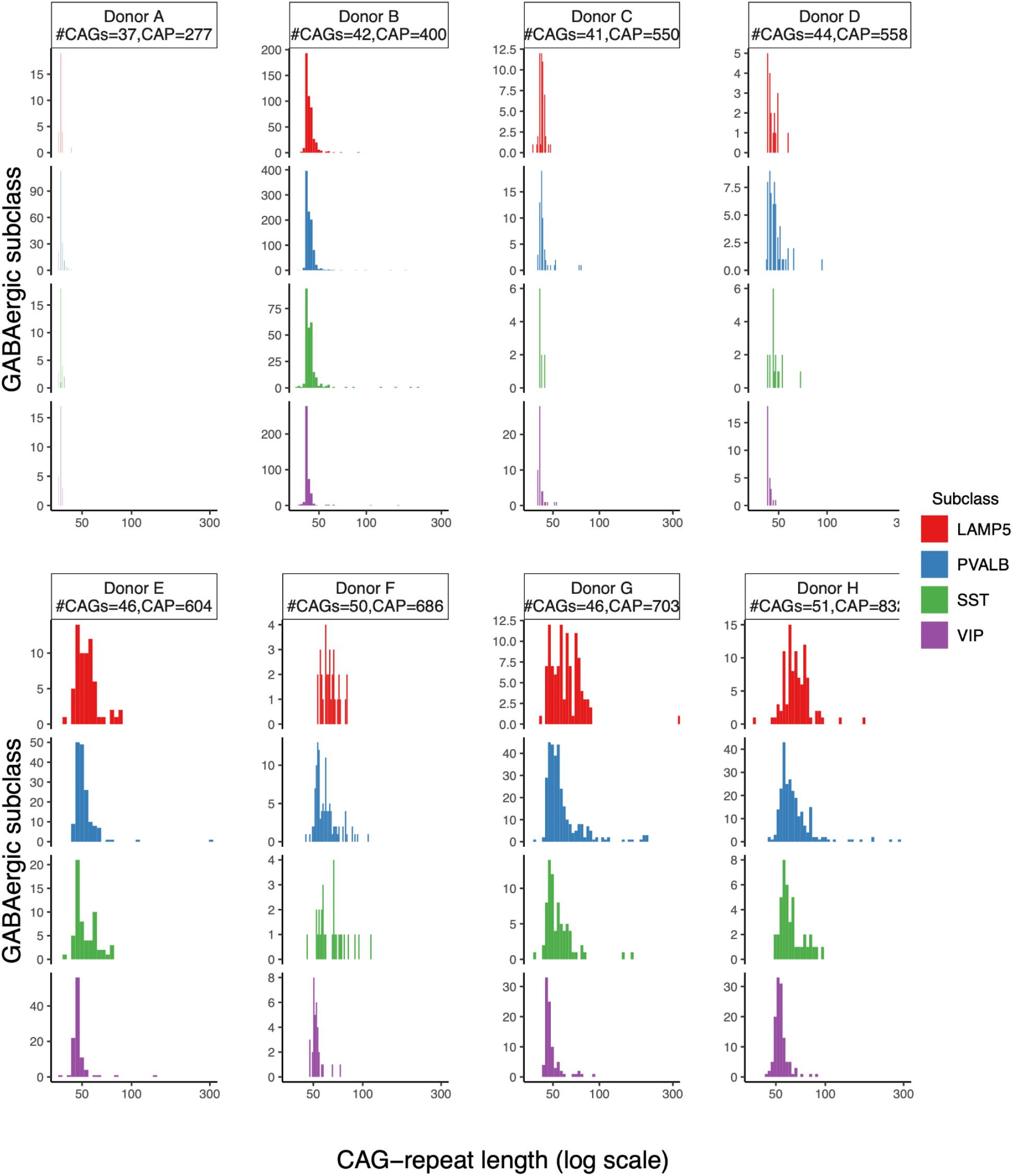
Somatic CAG-repeat expansion in specific types of cortical GABAergic neurons. Related to Fig. 1, here focusing on specific types of interneurons. Each row/color corresponds to a different type of GABAergic neurons. Data for eight donors (ordered by CAP score, an estimate of disease progression) are shown. LAMP5+ interneurons consistently show the most somatic expansion, though few of them have expanded their CAG-repeat tracts beyond 100 CAGs.

**Figure S2.**
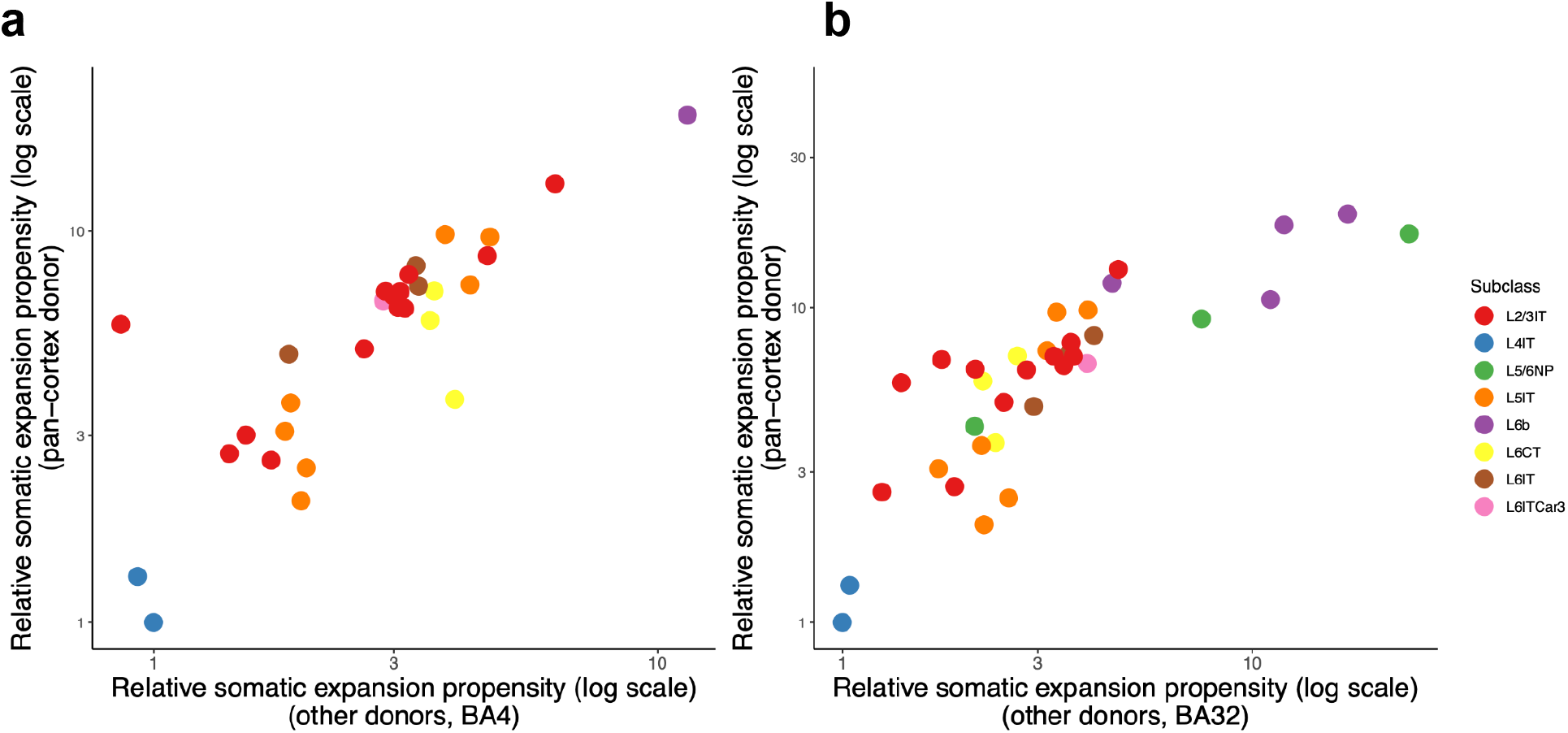
Replication across brain donors of estimates of the relative amounts of somatic *HTT* CAG-repeat expansion in different pyramidal neuron subtypes. Related to Fig. 3, which focused on data from an individual brain donor (the “pan-cortex donor, included again here on the y-axes for comparison to the other donors). Each circle corresponds to a pyramidal neuron subtype (from the high-resolution Siletti taxonomy) for which we have quantified somatic CAG-repeat expansion. These 20+ pyramidal subtypes are each indicated with a circle; the colors of the circles reflect supertypes (L2/3IT, etc.) from the earlier, lower-resolution (but more extensively annotated) Hodge taxonomy. On the y-axes are estimates from a deeply sampled brain donor from whose neocortex we sampled twelve cortical areas. On the x-axes are estimates from analyzing other brain donors, focusing on estimates from motor cortex (BA4, panel **a**) or anterior cingulate cortex (BA32, panel **b**). Note that, for L6b and L5/6NP pyramidal subtypes (purple and green circles), loss of the neurons with the most somatic expansion during the course of HD (see Fig. 6) would cause us to under-estimate their somatic expansion, by an amount that varies from donor to donor; this may be why these purple and green points diverge from the otherwise-linear replication relationship formed by the other pyramidal subtypes. Also note that, because some donors have more somatic expansion than others (due e.g. to age and genetic modifiers), the cardinal values of these estimates are donor-specific – it is their *relative* (not absolute) values that appear to be consistent across people.

**Figure S3.**
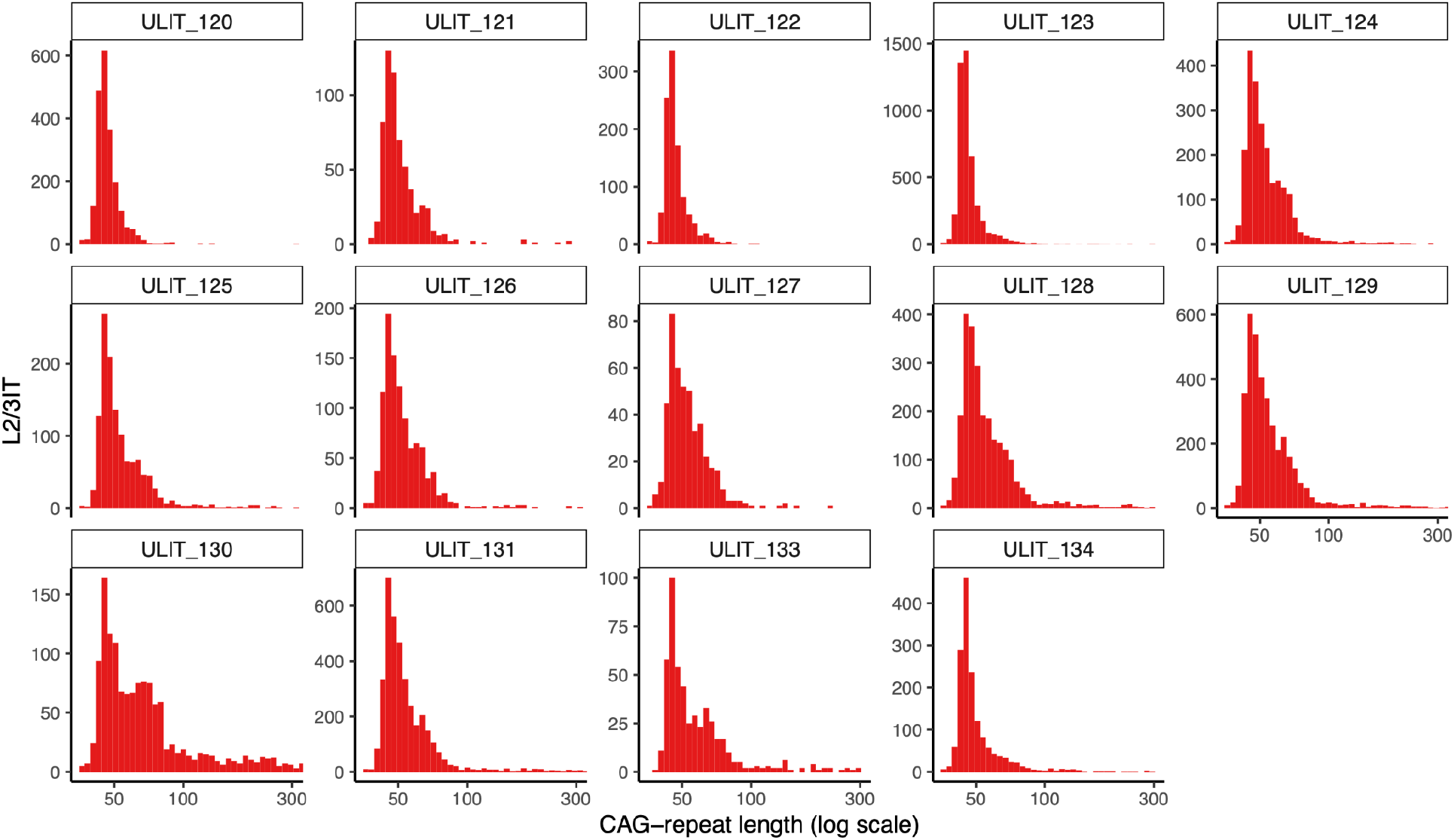
Distributions of somatic CAG-repeat tract lengths, focusing on the HD-causing allele, in subtypes of upper-layer pyramidal neurons. Related to Fig. 3. The nuclei were sampled from a person with HD (48yo, inherited 42-CAG allele, Vonsattel grade HD-1, “Donor B” in Fig. 1).

**Figure S4.**
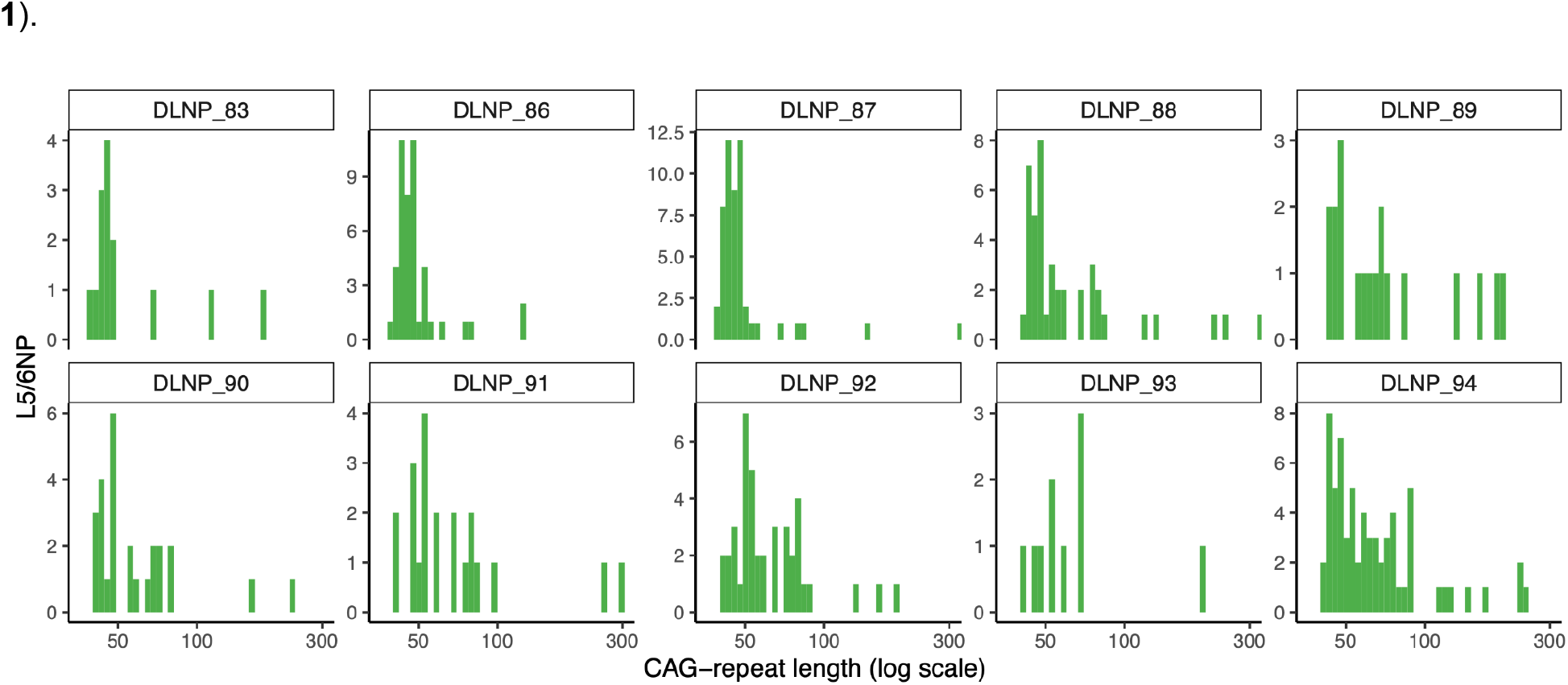
Distributions of somatic CAG-repeat tract lengths, focusing on the HD-causing allele, in subtypes of deep-layer near-projecting (L5/6NP) pyramidal neurons. Related to Fig. 3. The nuclei were sampled from a person with HD (48yo, inherited 42-CAG allele, Vonsattel grade HD-1, “Donor B” in Fig. 1).

**Figure S5.**
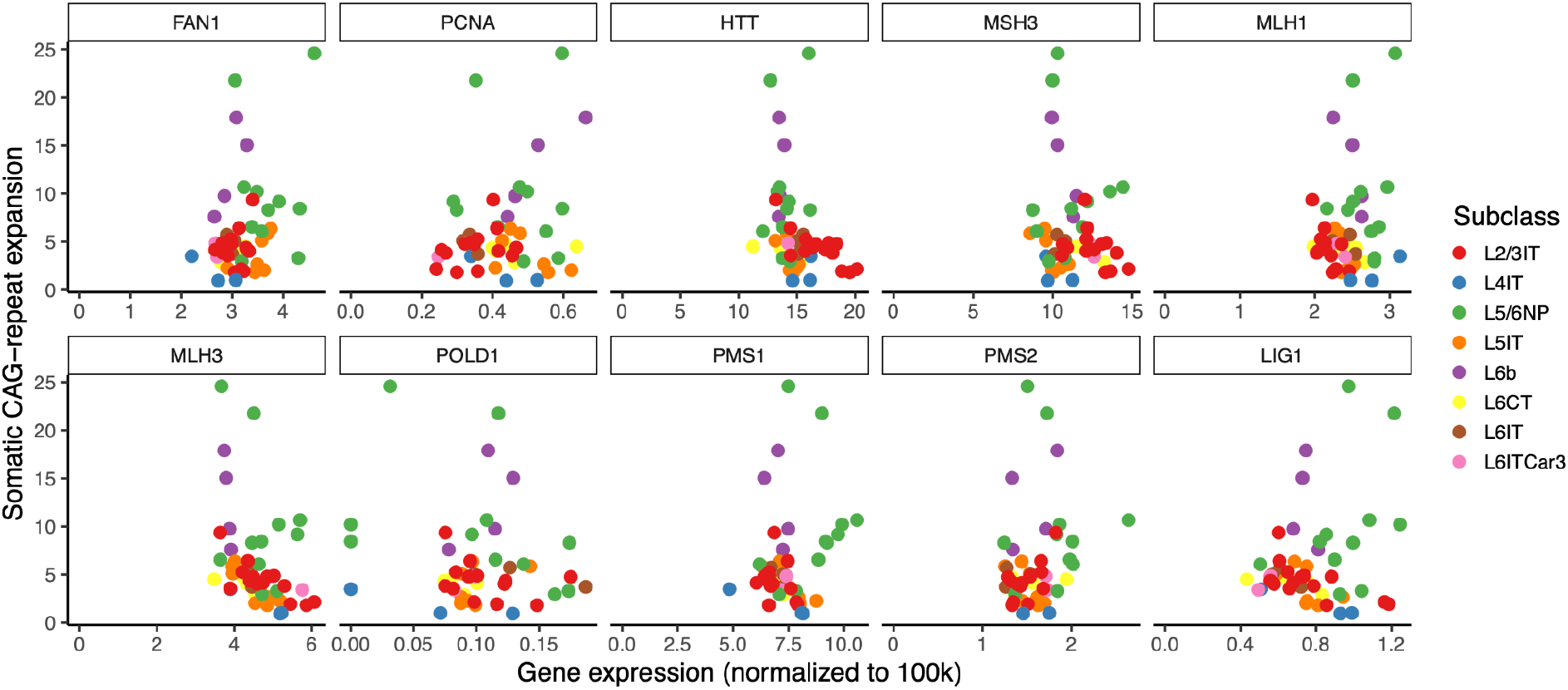
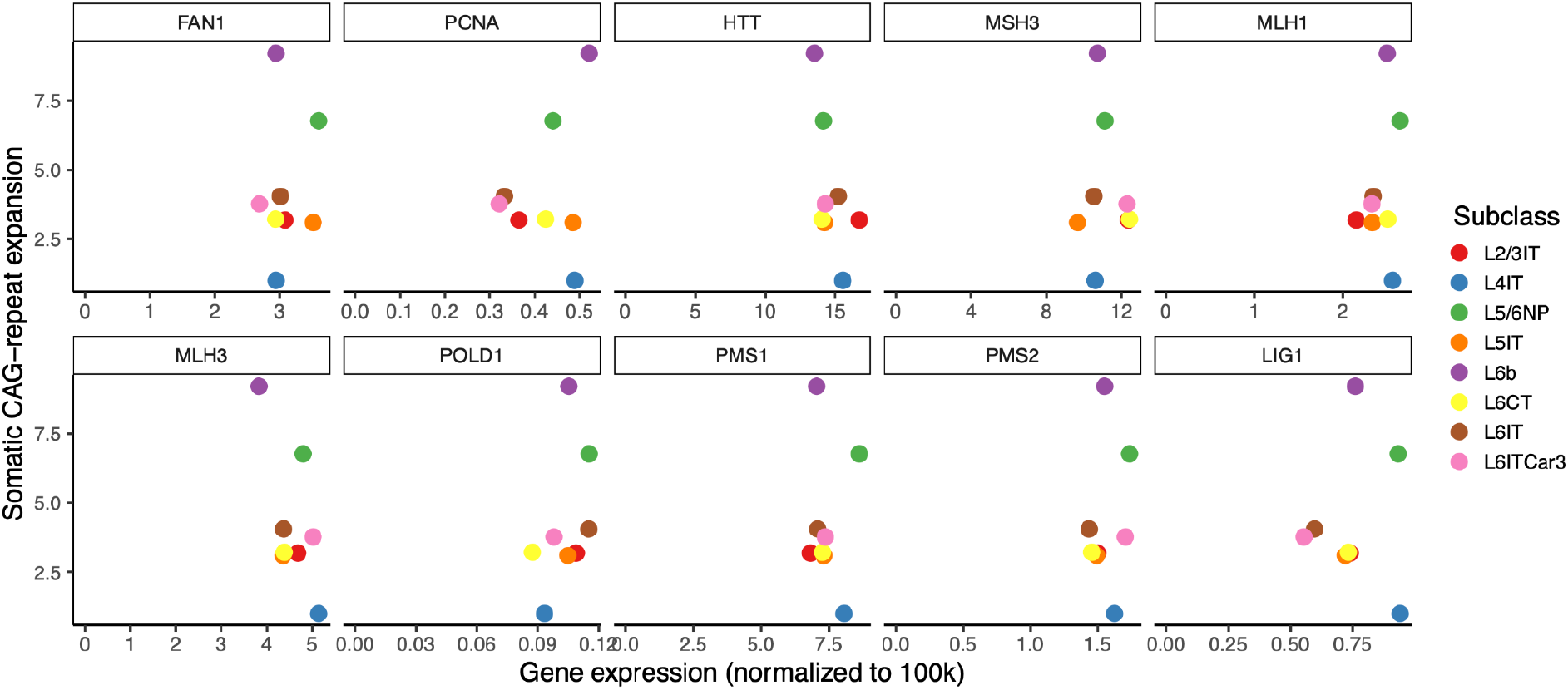
Relative rates of somatic *HTT* CAG-repeat expansion across pyramidal neuron types and subtypes, in relationship to nuclear RNA expression levels of *HTT* and a variety of genes known or proposed to regulate the stability of the *HTT* CAG repeat. Each point represents a type or subtype of pyramidal neuron; the points’ y-axis locations reflect relative amounts of somatic expansion, and their x-axis locations reflect nuclear RNA expression levels (measured in UMIs per 100k UMIs samples) of the gene shown. (a) Analysis of high-level pyramidal neuron types. Note that rigorous statistical analysis of such relationships is challenging, because the individual points (cell types) are non-independent: cell types have a complex, structured, and partially hierarchical set of similarity/dissimilarity relationships, and cannot be considered independent in the way that statistical tests of correlation assume. We have not reported p-values because p-values from statistical tests that do not incorporate these complex relationships are misleading, but can report that, even in a naive analysis, none reached nominal significance in parametric or non-parametric correlation tests. (b) As in (a), but dis-aggregated into pyramidal subtypes. Note that here, the statistical challenge posed by the hierarchical similarity-dissimilarity relationships of the cell types is particularly acute: for example, *PMS1* may have hints at an upward-sloping relationship, though these are driven by subtypes of L5/6NP neurons (which are similar to one another and thus non-independent observations). Some L5/6NP subtypes (green points) with particularly high relative somatic expansion (>8) tended to have relatively high expression of several mismatch repair genes (e.g. *MLH3, MLH1, PMS1, LIG1*), though again, these closely related neuronal subtypes cannot be considered independent observations. It is important to remember that individual genes tend to change in expression in the context of large gene-expression programs that involve concerted changes in the expression of very many other genes – often hundreds of genes – together. Thus, expression measurements for a single gene are also indirect proxies for many other genes’ expression levels, and for unmeasured cellular properties (such as neuronal activity patterns or metabolic states) that may also affect somatic instability. In this context, one should be reluctant to attribute causality to individual genes based on observational/correlational analyses.

**Figure S6.**
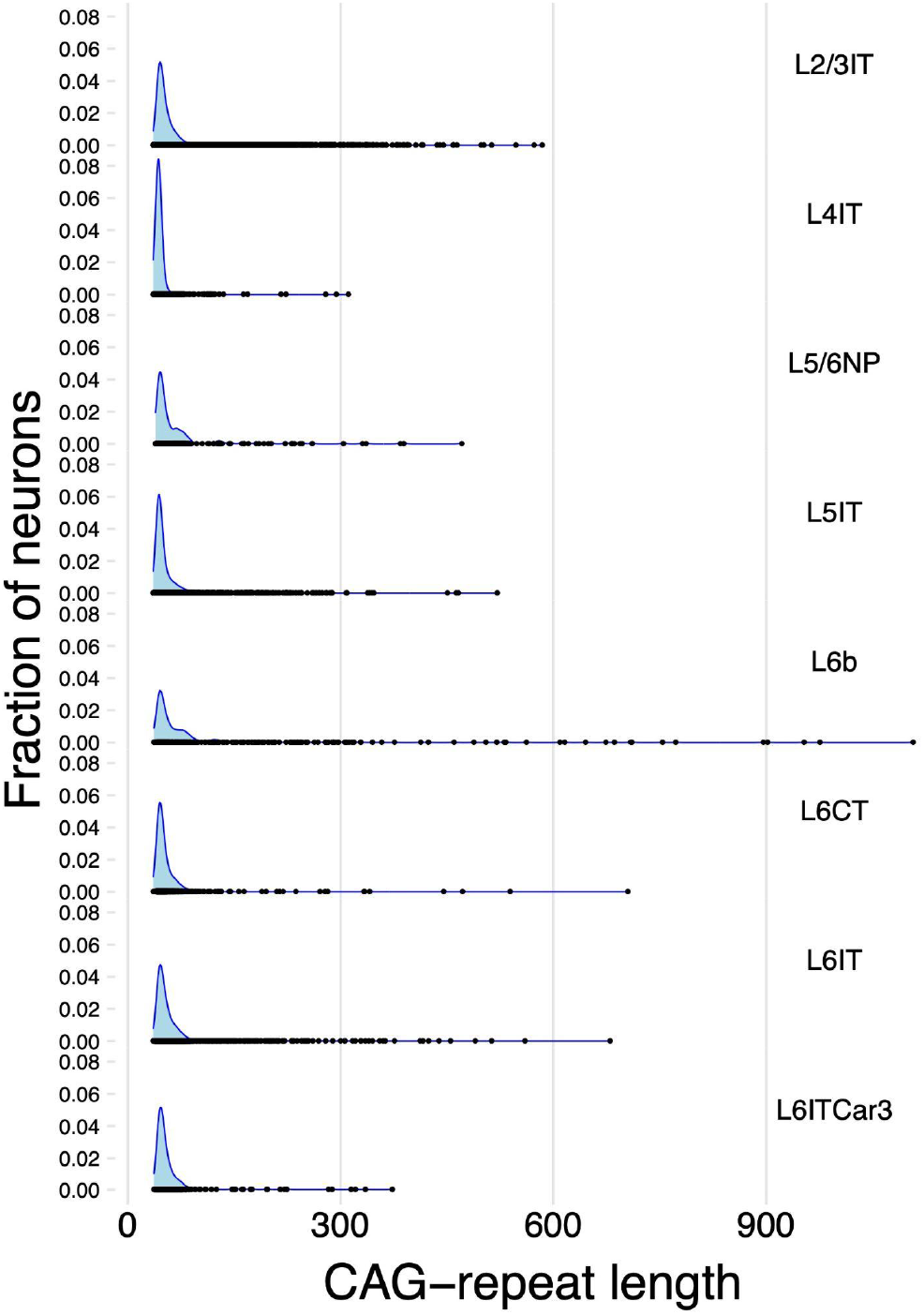
Distributions of CAG-repeat lengths in eight types of cortical pyramidal neurons from the same person with HD. Blue shaded areas are smoothed density estimates of the repeat-length distribution. Overplotted black points show the CAG-repeat length measurements in individual glutamatergic neurons. In all eight glutamatergic neuron subclasses, the repeat-length distribution exhibits an armadillo-like shape, in which the DNA-repeat tract in most neurons has undergone modest expansion (up to about 90 CAGs) but in a small fraction of neurons it has undergone far greater expansion (up to 500+ CAGs).

**Figure S7.**
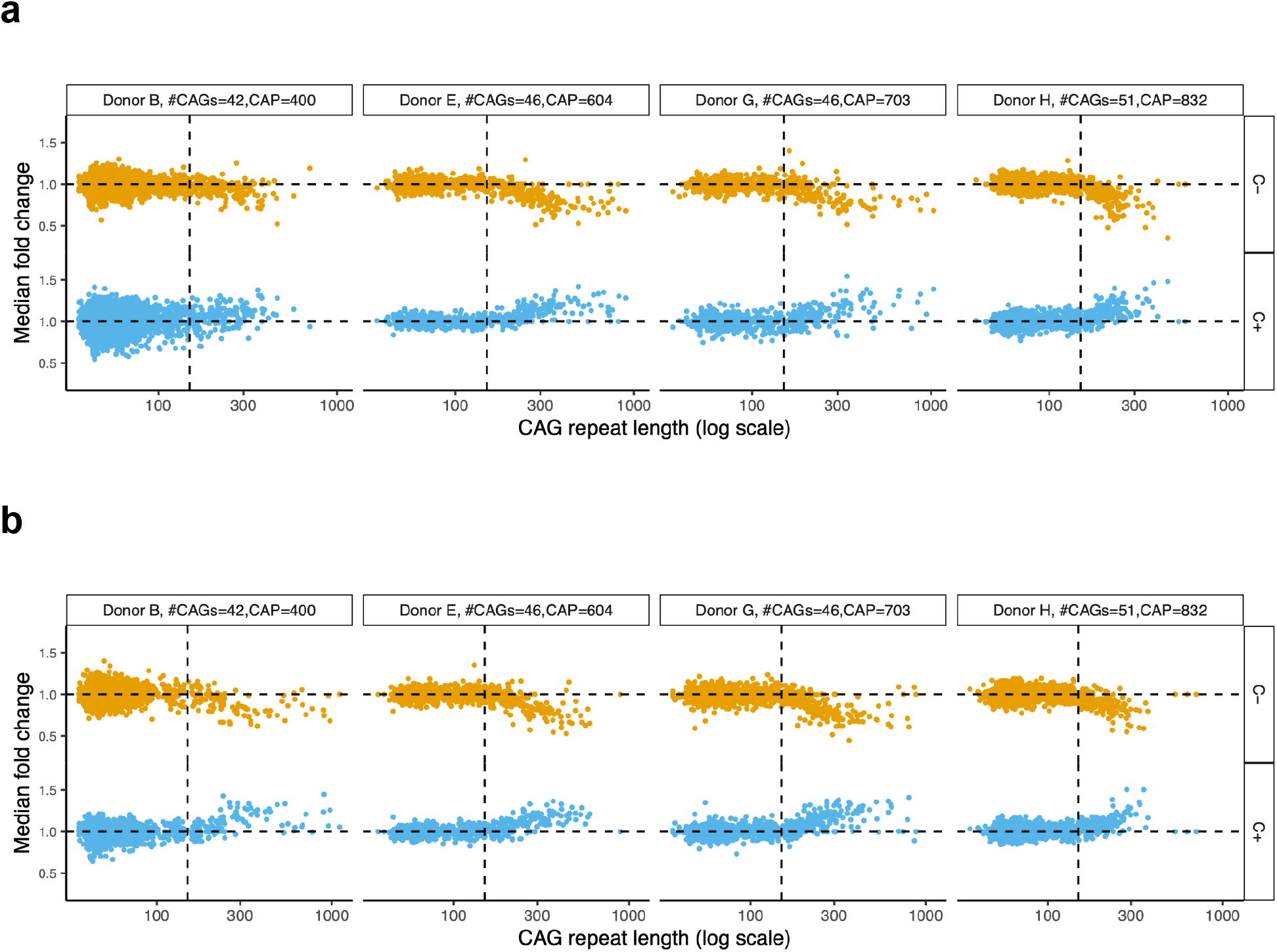
Phase C gene-expression changes appear in cortical neurons with somatic CAG-repeat expansion beyond 150 CAGs. Related to Fig. 4, and extending analysis to additional persons with HD. Analyses of (**a**) motor cortex (BA4) and (**b**) anterior cingulate cortex (BA32). On each pair of panels, each pyramidal neuron is represented by both a blue point (lower plot) and an orange point (upper plot): blue points show the median fold-change of a set of 115 genes that decrease in expression with CAG-repeat expansion (C- genes); orange points show the median fold-change of a set of 142 genes that increase in expression with CAG-repeat expansion (C+ genes).

**Figure S8.**
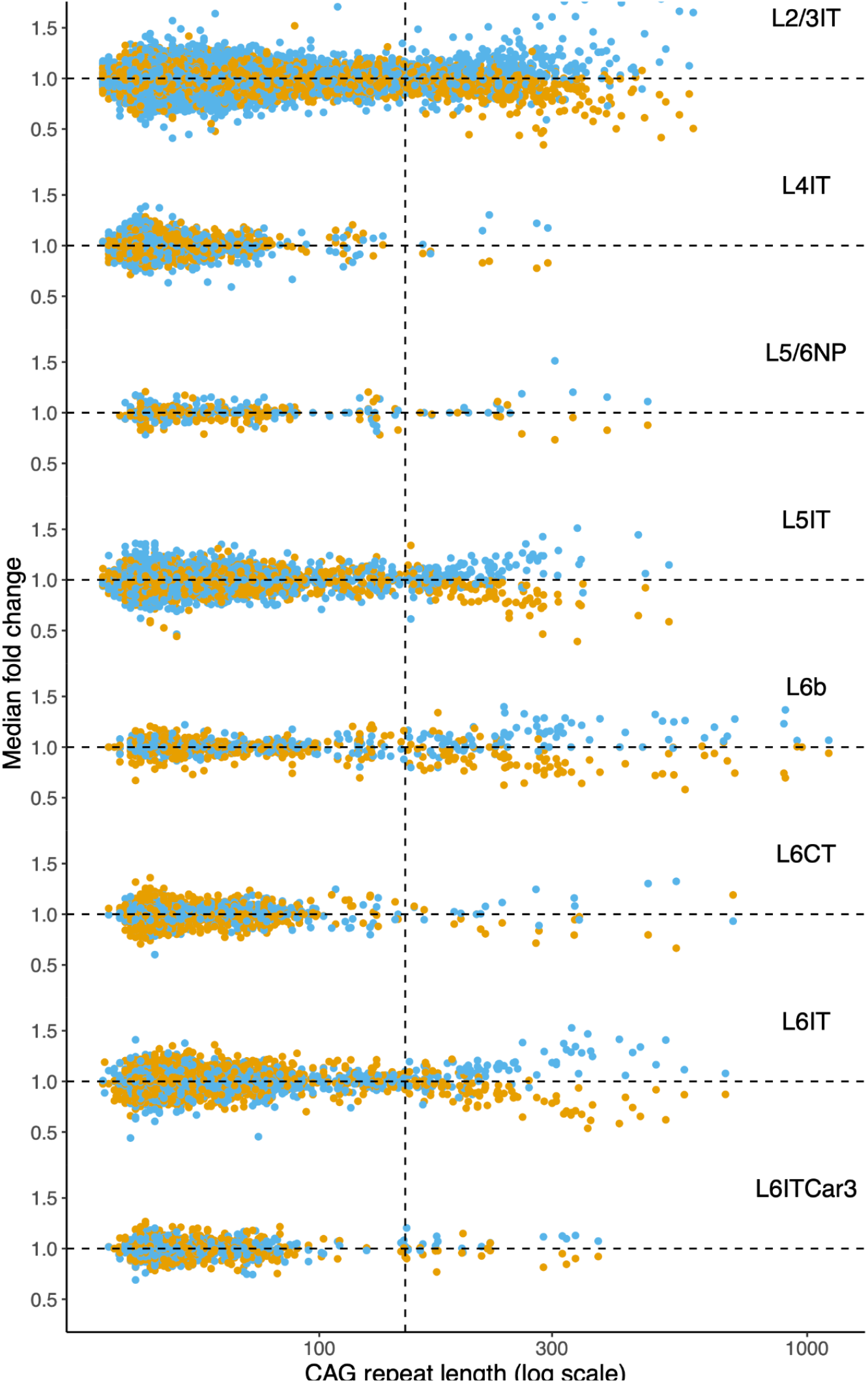
Gene-expression changes with somatic CAG-repeat expansion beyond 150 CAGs in each type of pyramidal neuron. Related to Fig. 4, but here drawing upon the 12 cortical areas sampled in one of the donors. Each of the eight plots shows the data for a specific type of cortical pyramidal neuron. On each plot, each neuron is represented by both a blue point and an orange point: orange points show the median fold-change of a set of 144 genes that decrease in expression with CAG-repeat expansion (C- genes); blue points show the median fold-change of a set of 179 genes that increase in expression with CAG-repeat expansion (C+ genes). The same genes were used for the calculation in each panel, and the 12 sampled cortical areas were used together. Fig. 4a contains analogous analyses of nuclei sampled from other persons with HD.

**Figure S9.**
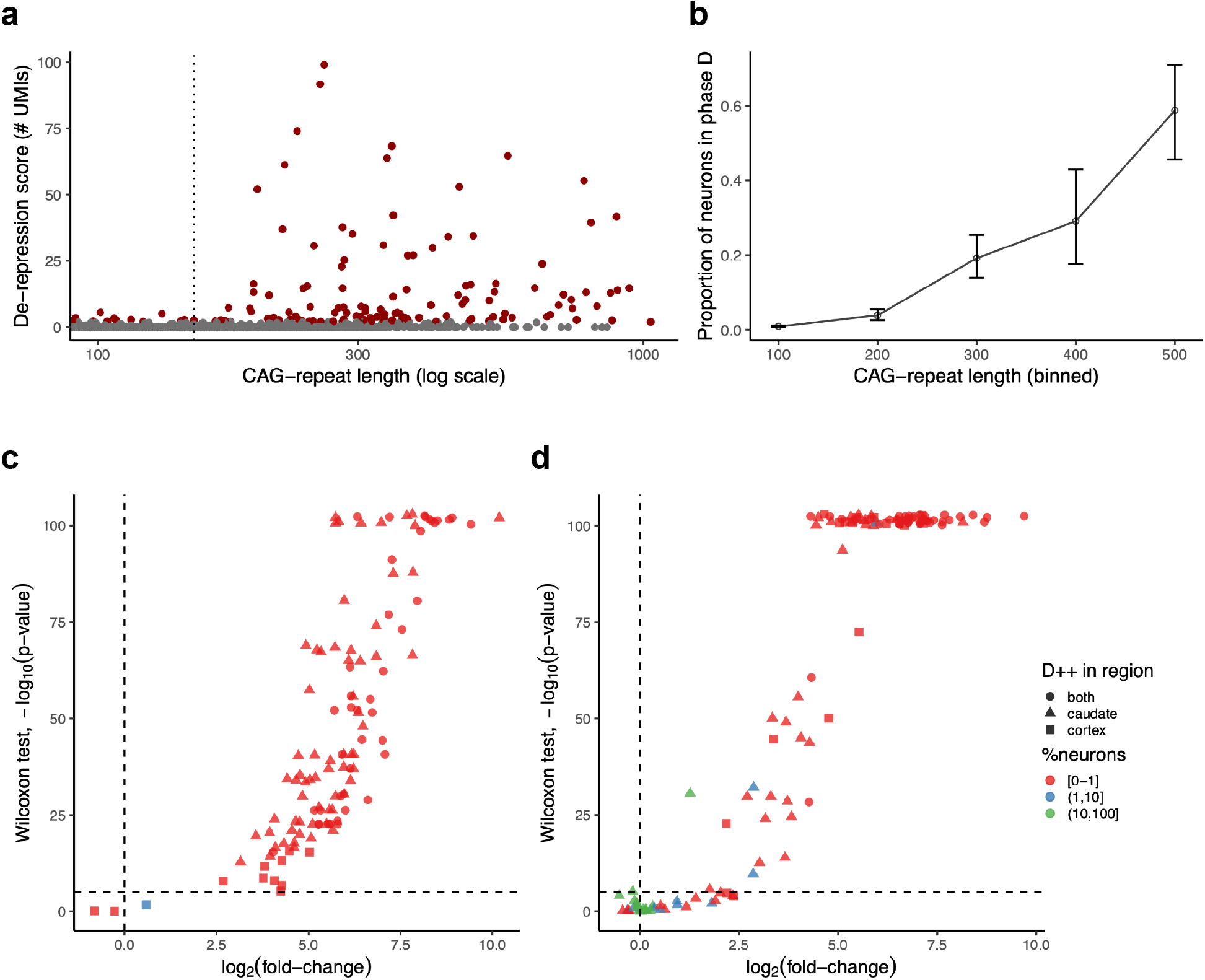
Phase D gene-expression changes. Related to Fig. 5, and drawing upon additional brain donors with HD. (**a**) Phase D changes in additional brain donors (shown all together). Points represent individual neuronal nuclei. y-axis: de-repression score, the number of transcripts (UMIs) detected from 59 phase D genes that are normally silent in cortical projection neurons. (**b**) Fraction of pyramidal neurons that have entered phase D, in relationship to CAG-repeat length (binned). (**c,d**) Sharing of Phase D de-repressed (D++) genes between striatal and cortical neurons. There were 124 phase D++ genes that had (in either cortex or striatum) the properties of being (i) silent at baseline (i.e. in neurons with <150 CAGs), and (ii) expressed in phase D neurons. Plots in panels c and d show how these 124 genes changed in expression in separate analyses of phase D striatal neurons (panel c) and of phase D cortical neurons (panel d). For each gene (point), location on the x-axis shows the log-fold-change when comparing phase D neurons to neurons at baseline (<150 CAGs); the y-axis shows the statistical significance of the comparison (Wilcoxon test). The color of the points reflects the fraction of neurons (with <150 CAGs) in which any RNA from the gene was detected. Note that the genes that fail to show de-repression in cortical neurons (the points below the dashed line in panel d) are for the most part genes that are already expressed in cortical neurons at baseline (as shown by the colors of the points). These include genes encoding the cortical transcription factors NFIX, LHX2, TBR1, TLE3, and TLX2, as well as genes with other neuron-type-specific functions (SLC17A7, KCNC2, ALDOC, PMS3, ARL4C).

**Figure S10.**
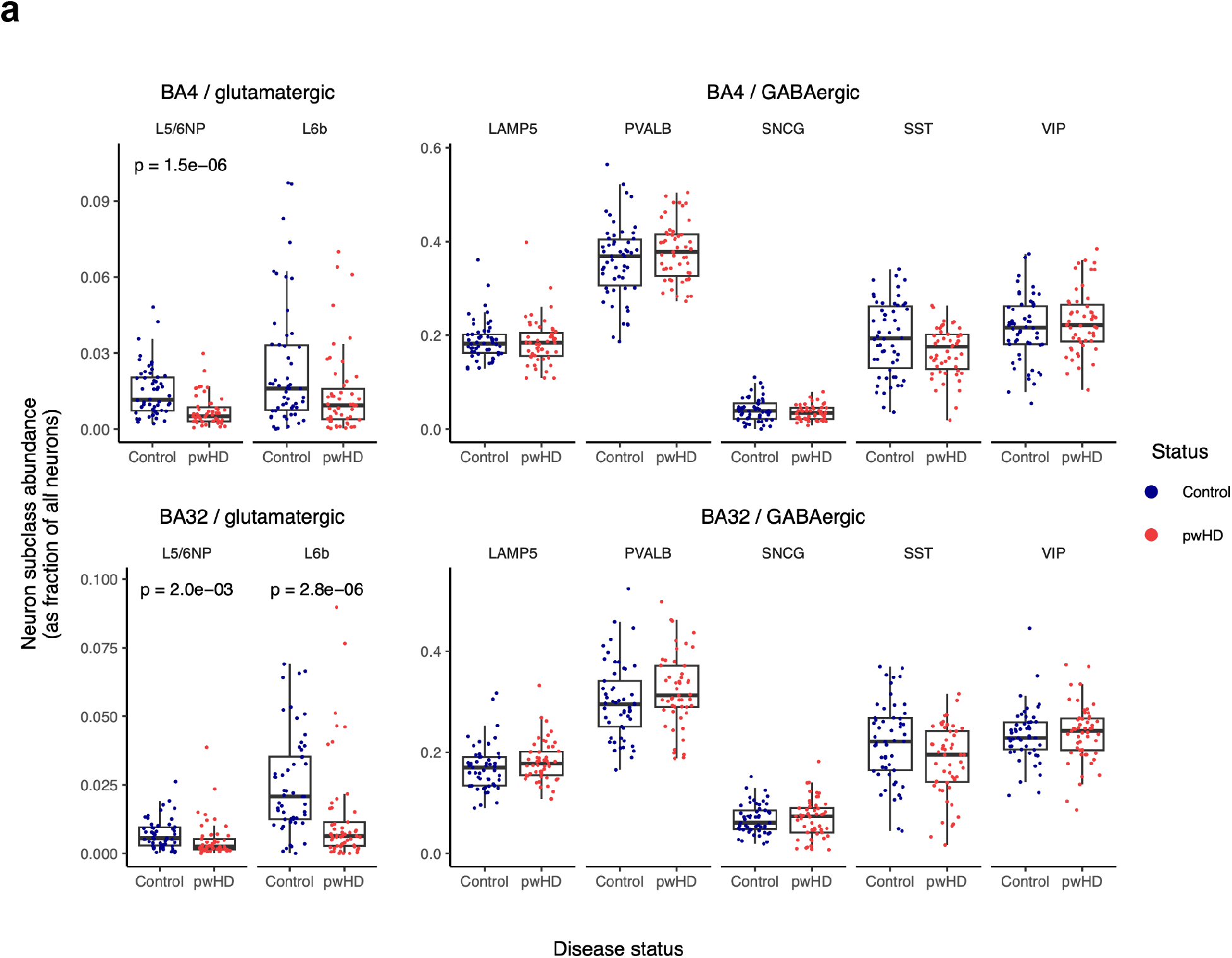
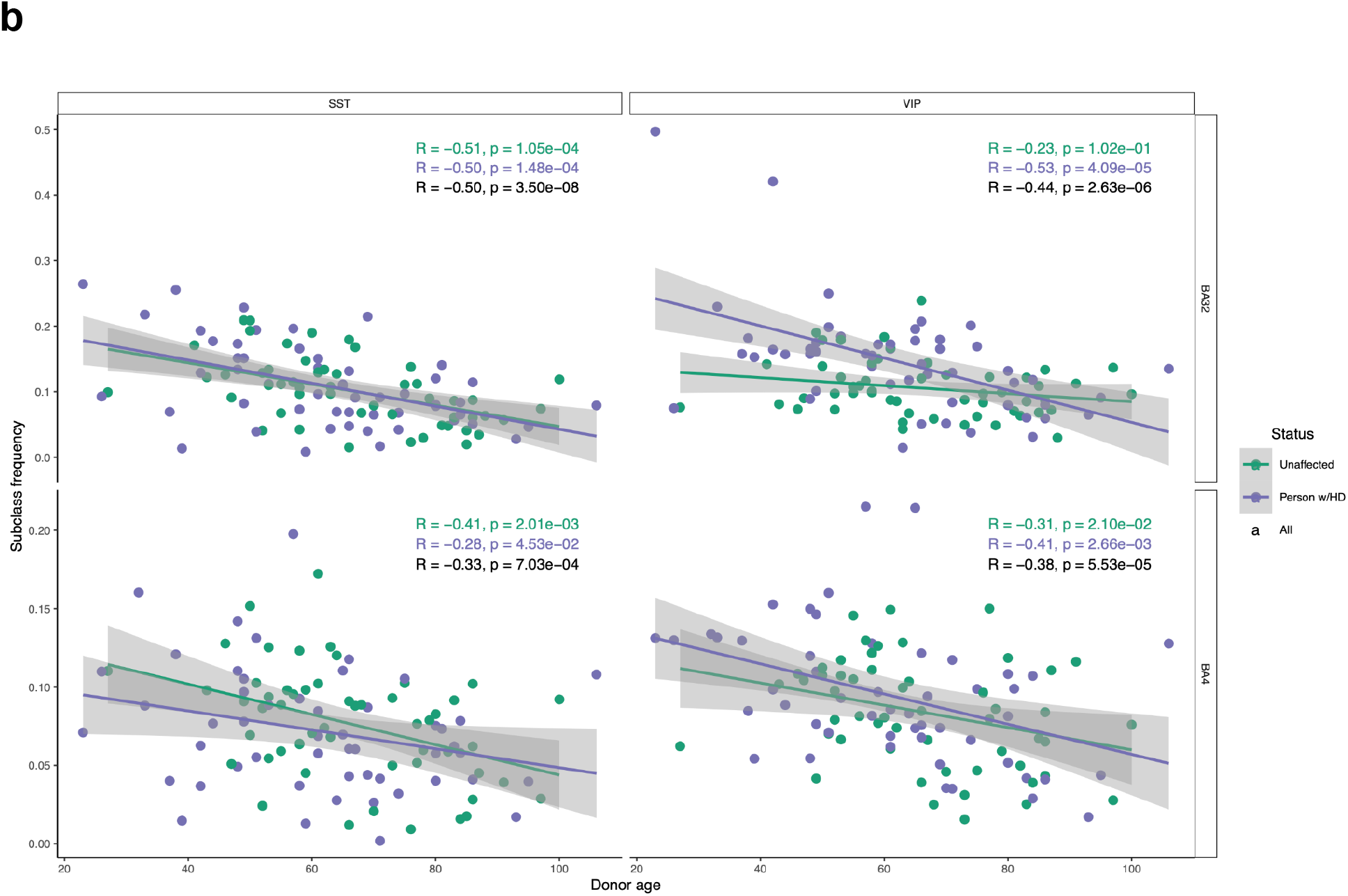
Attrition of cortical interneurons due to age but not HD. Related to Fig. 6. (a) Cortical interneurons (GABAergic neurons) exhibited no consistent difference in abundance between 50+ brain donors with HD and 50+ unaffected brain donors – in contrast to L5/6NP and L6b pyramidal neurons (leftmost panels). Results are from motor cortex (BA4, upper set of panels) and anterior cingulate cortex (BA32, lower set of panels). The p-values are calculated from the Wilcoxon (non-parametric) test so as to be resistant to outliers and are nominal p-values (not corrected for the number of cell types tested). Note that the abundances of several types of GABAergic interneurons exhibit striking inter-individual variation, but that (in this 50 HD vs. 50 unaffected analysis) the differences between HD-affected and unaffected individuals were not even nominally significant in either BA4 or BA32. (b) Attrition of SST and VIP interneurons with age, in both BA32 (upper panels) and BA4 (lower panels), among both unaffected persons (green points) and persons with HD (purple points). The green and purple regression lines (and correlation statistics) were separately fit to the data from unaffected persons (green) and persons with HD (purple).

**Figure S11.**
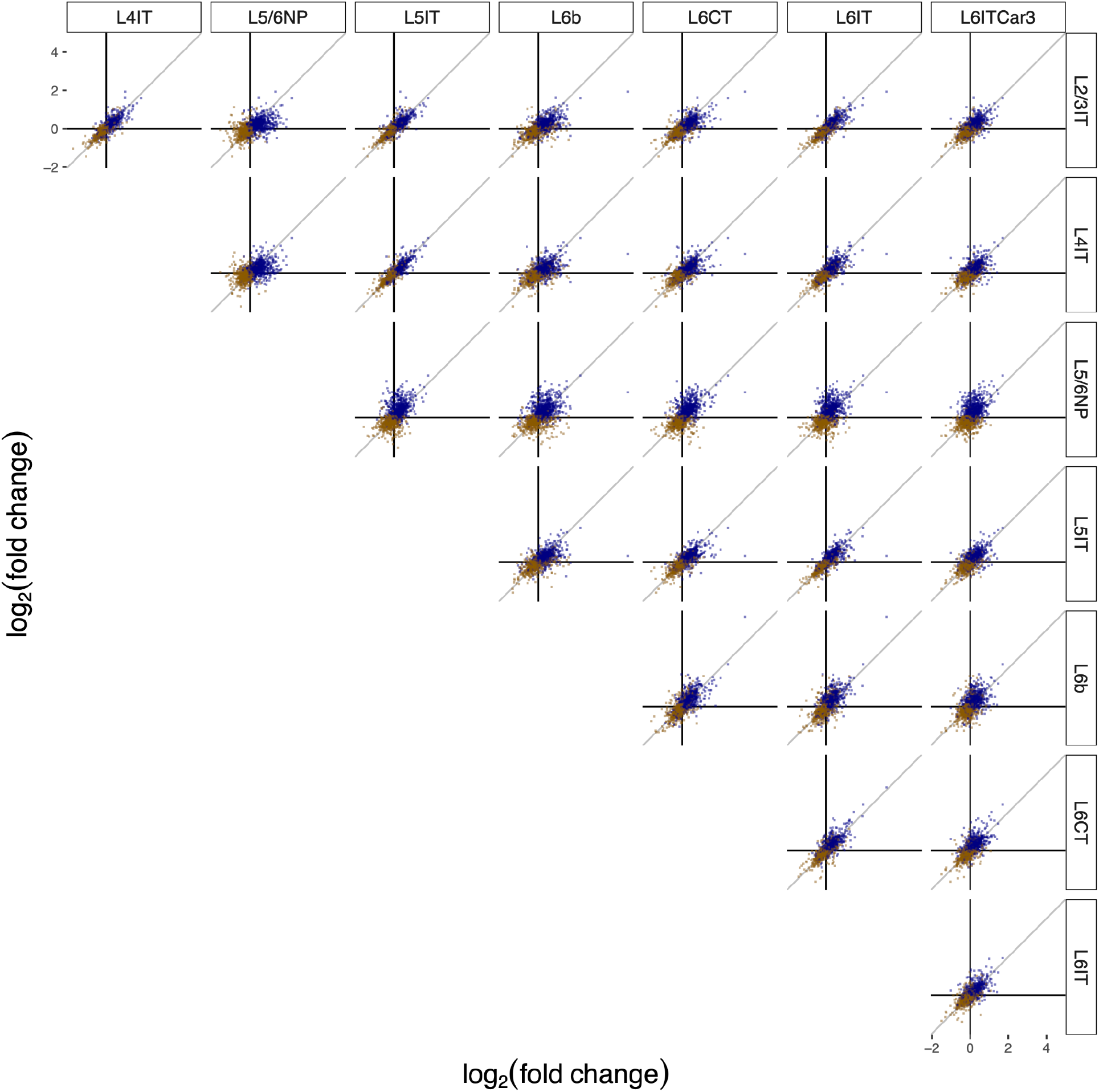
Replication of case-control differentially expressed genes (DEGs) across all types of pyramidal neurons, regardless of their vulnerability in HD. Related to **Results** section on non-cell-autonomous sequelae of neurodegeneration. DEGs were identified by comparing gene expression in persons with HD vs. unaffected individuals in each of the pyramidal neuron subclasses. Each point corresponds to a gene found to be differentially expressed in the current work’s differential-expression analyses of persons with HD (relative to unaffected individuals) in at least two pyramidal neuron subclasses and is also present in the list of differential expressed genes for L5a in the study by Pressl et al; expression fold-changes (HD/unaffected) are shown for pairs of pyramidal neuron types. The locations of points (genes) on the plot are based on fold-change in the current data (comparing HD-affected to unaffected brain donors). Points are colored blue or orange based on whether they were found to be up- or down-regulated in HD-impacted neocortex in the study by Pressl et al.

**Figure S12.**
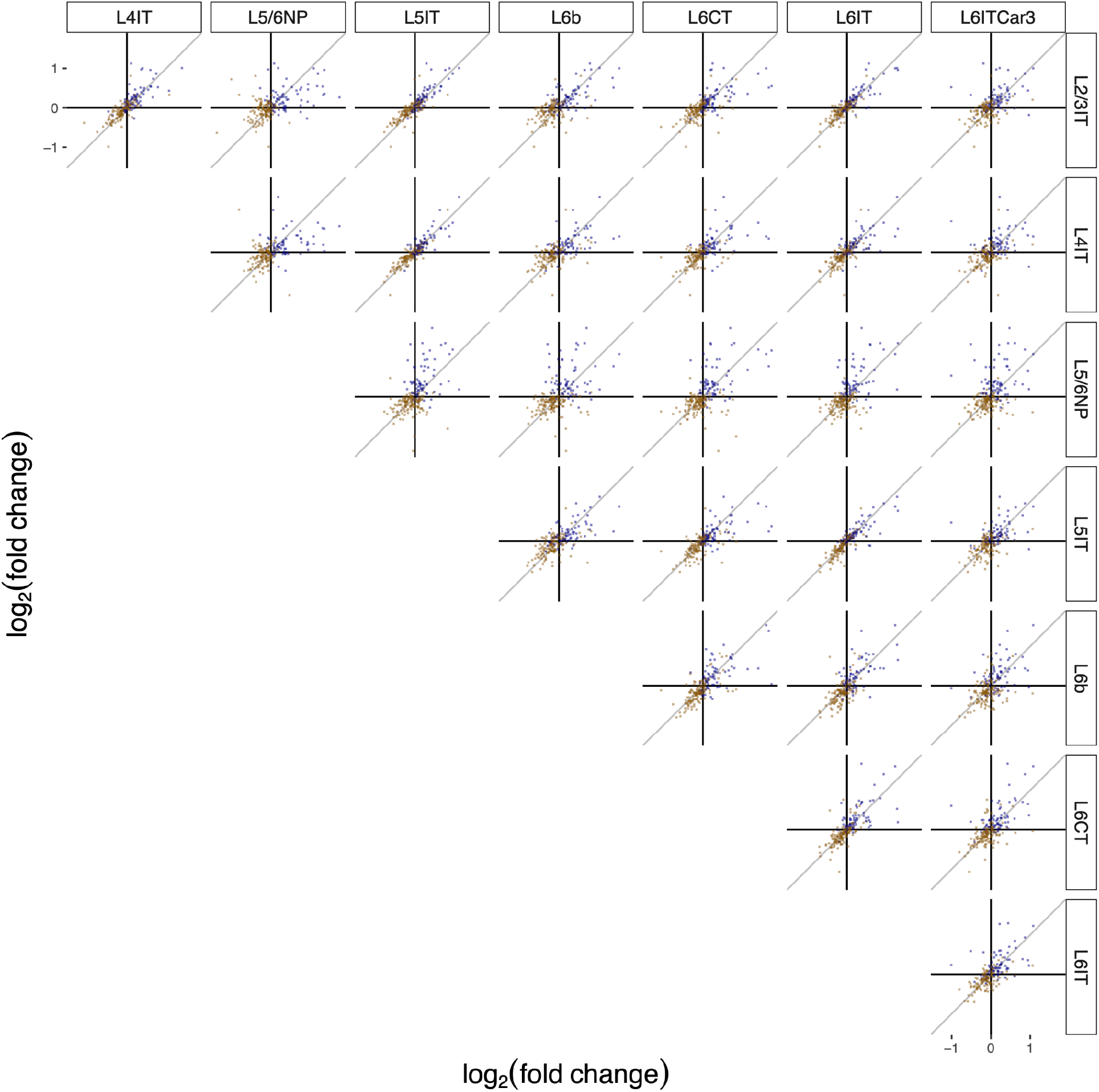
Similar to **Fig. S11**, but subsetting to the set of genes in the “synapse” Gene Ontology cellular component category. The locations of points (genes) on the plot are based on fold-change in our own data (comparing HD-affected to unaffected brain donors). Points are colored blue or orange based on whether they were found to be up- or down-regulated in HD-impacted neocortex in the study by Pressl et al.

**Figure S13.**
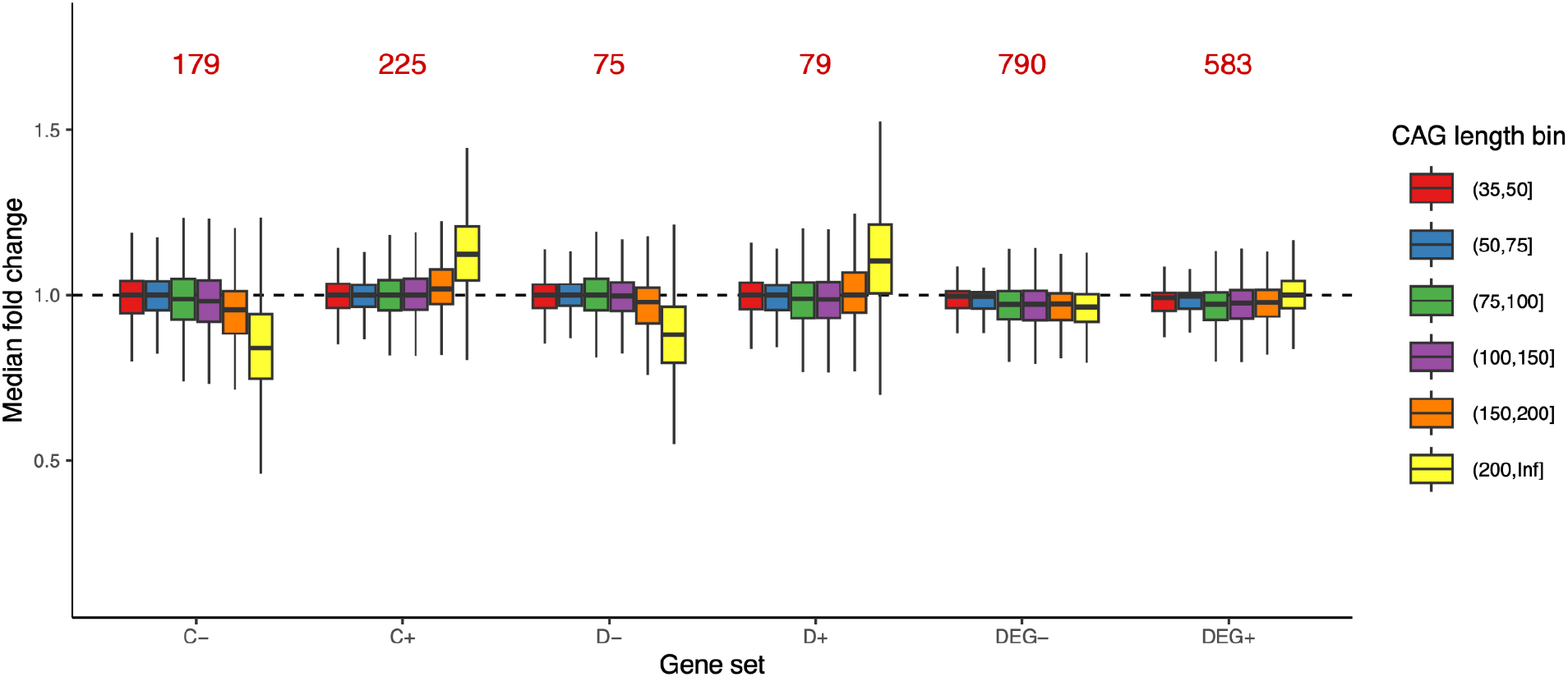
Relationship of six types of gene-expression changes to length of a pyramidal neuron’s CAG-repeat tract. Related to **Results** section on non-cell-autonomous sequelae of neurodegeneration. Gene sets considered, from left to right: phase C genes (C- and C+ denote down- and up-regulated genes); phase D genes (D- and D+); and “differentially expressed genes” in L5a pyramidal neurons identified in the study by Pressl et al (DEGs, divided into DEG- and DEG+ gene sets based on whether expression tended to increase or decrease in HD-affected relative to unaffected neocortex). Each of these gene sets was filtered to contain genes expressed in at least 50% of all pyramidal neurons. The quantity plotted is the median fold-change among those genes in the set. The numbers in red above the bar plots show the numbers of genes in each filtered gene set.

**Figure S14.**
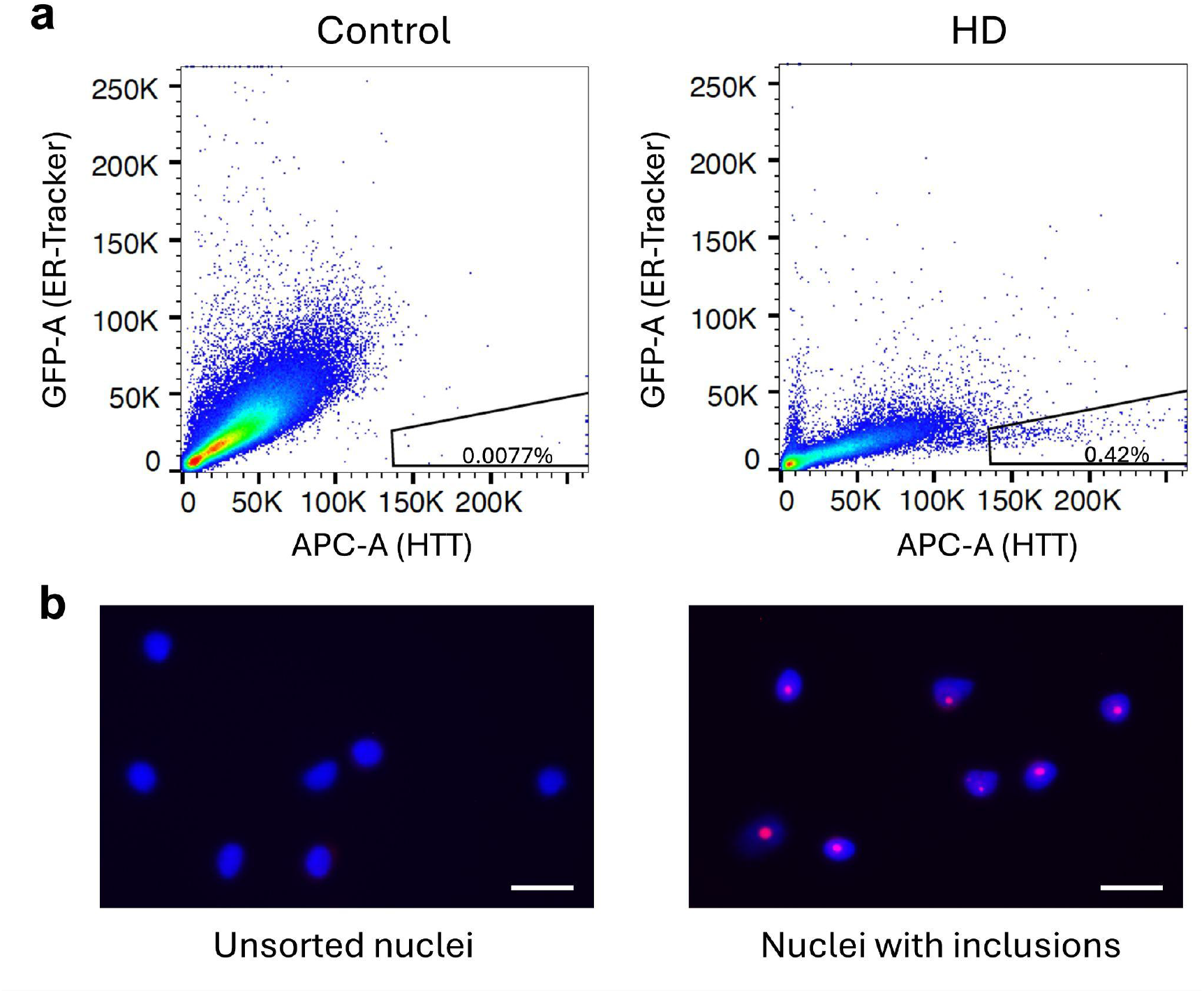
Sorting gating strategy for isolating nuclei with HTT inclusions. Related to Fig. 7. (**a**) Representative FANSII gating of nuclei isolated from a control donor (left) and a donor with HD (right). Nuclei containing HTT inclusions were gated as HTT-high and ER-Tracker-low to minimize potential autofluorescence and non-specific fluorophore binding associated with residual endoplasmic reticulum content. Notably, few to no nuclei fall within this gate in the sample from the unaffected (control) brain donor. (**b**) Representative immunofluorescence images of nuclei from unsorted (left) and sorted (right) fractions from an HD donor using this gating strategy. Scale bar = 25 µm.

**Figure S15.**
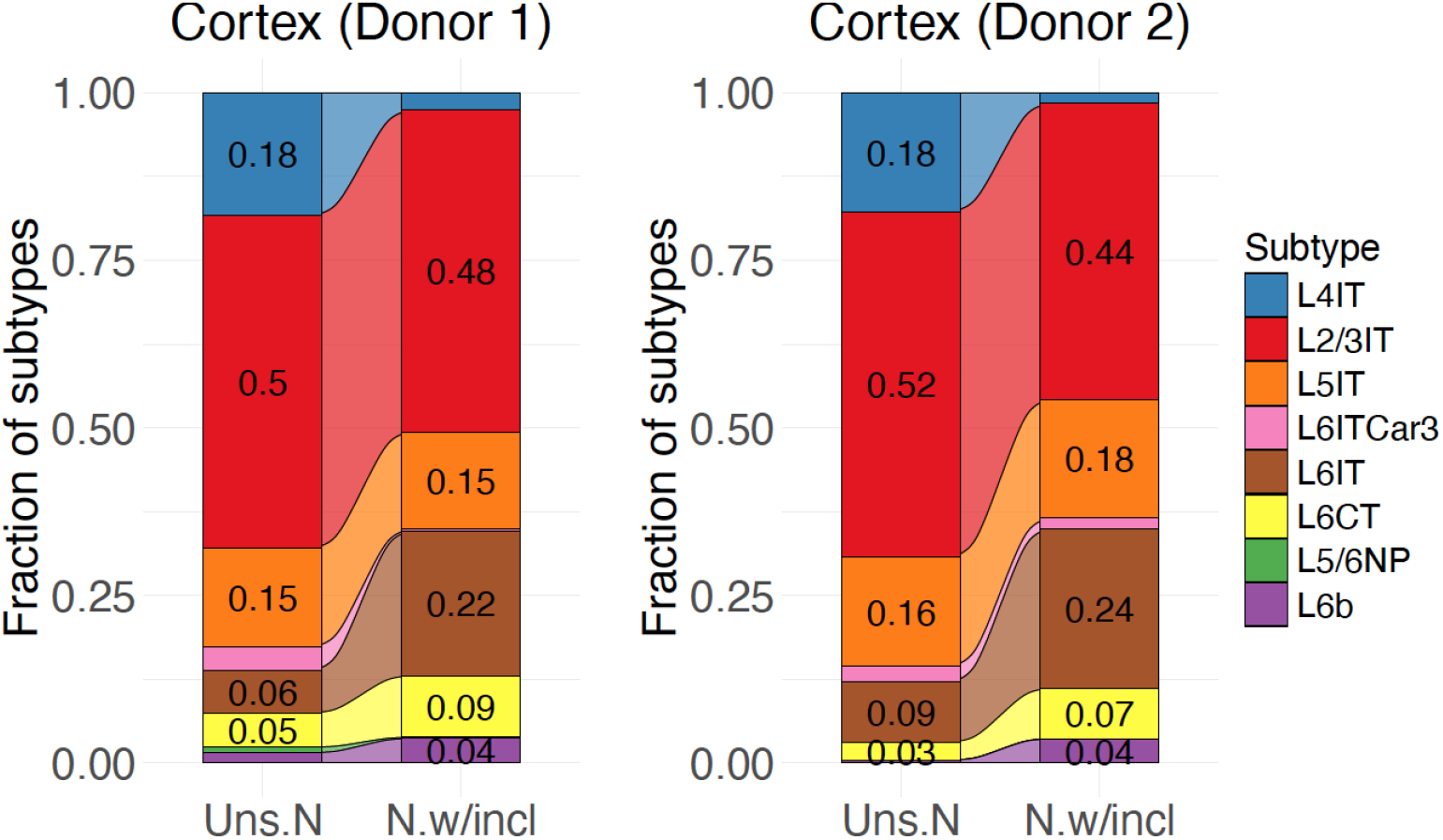
Representation of pyramidal neuron types among nuclei with HTT inclusions, in comparison to all (unsorted) neuronal nuclei. Related to Fig. 7c, here expanding to consider the various types of pyramidal neurons. The sorted and unsorted nuclei were analyzed by snRNA-seq; subtype identities were inferred from the genome-wide RNA data in each individual nucleus. Here the pyramidal neuron types have been ordered by their relative amounts of somatic expansion (inferred from earlier analyses in other donors, shown in Fig. 2c), from least expansive (L4IT, blue, top) to most expansive (L6IT, L6CT, L5/6 NP, L6b; brown, yellow, green, purple, bottom). Note that this analysis reflects subtype composition at a specific time point (end of life) and therefore has an indirect and potentially complex relationship to cumulative cell loss through the course of illness; for example, L5/6 NP neurons were already so depleted as to be less than 1% and 0.1% of pyramidal neurons in the two donors (in unsorted nuclei as well as sorted nuclei with HTT inclusions).

**Figure S16.**
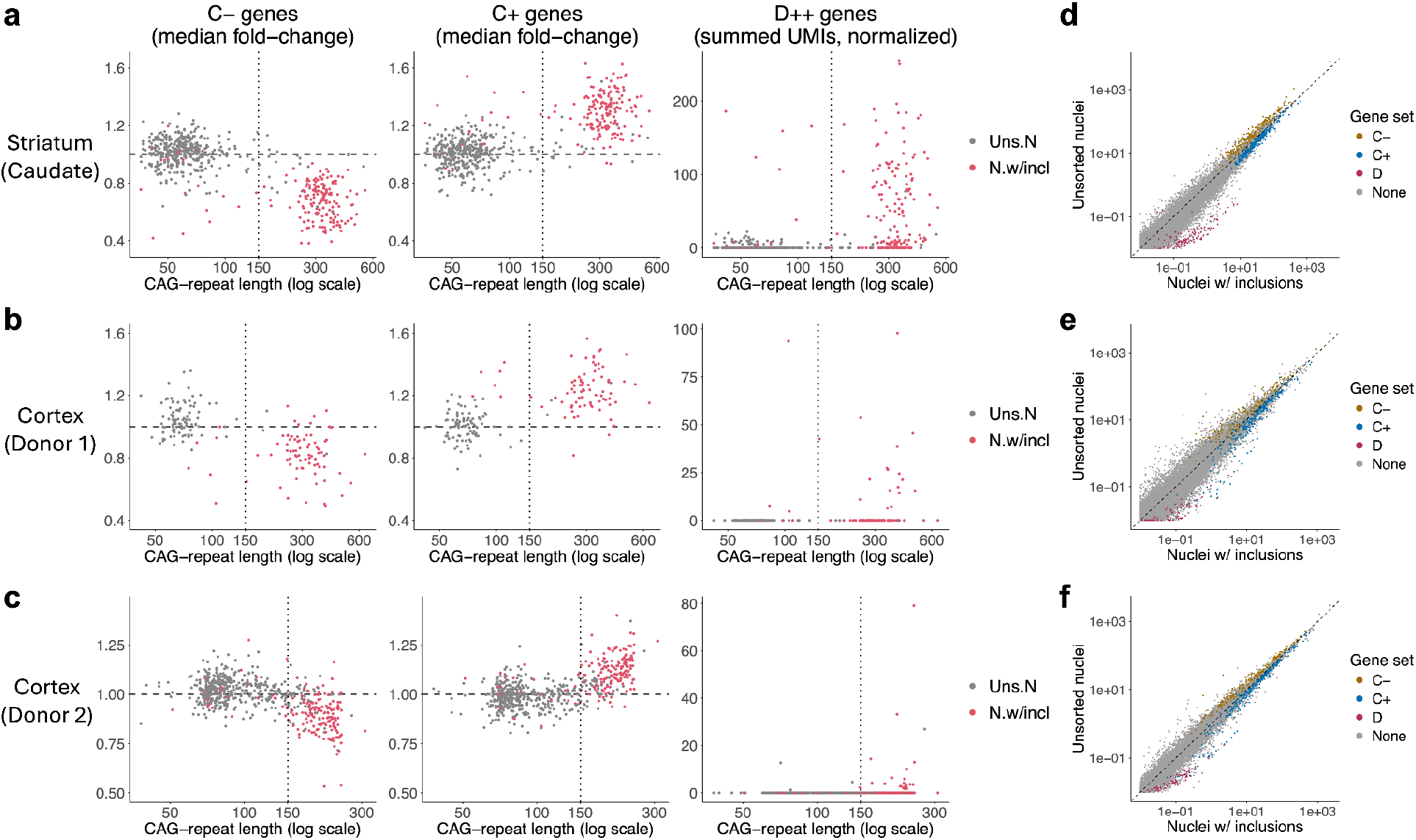
Gene expression changes in neuronal nuclei with inclusions. Related to Fig. 7. (**a-c**) Phase C and D gene-expression changes in nuclei with inclusions (red points) in comparison to unsorted nuclei (grey points). From left to right: expression changes in individual nuclei for phase C- genes , phase C+ genes, and phase D++ genes. (Expression changes for C- and C+ genes are quantified in terms of median fold-change relative to baseline expression level; and for D++ genes, which are non-expressed or negligibly expressed at baseline, in terms of the total summed number of UMIs (per 100k) ascertained across these 59 genes.) Data are from the same experiments as in Fig. 7, but here including only those nuclei with CAG-repeat measurements for the HD-causing allele. The fold-change calculations are described in Methods, and the genes used in the calculations are listed in Supplementary Data. (**d-f**) Genome-wide comparisons of gene-expression levels between nuclei with inclusions (x-axis) and unsorted neuronal nuclei (y-axis). Phase C-, C+, and D++ genes are shown with orange, blue and red points respectively; all other genes are shown with gray points.

## Notes

https://www.nemoarchive.org/

### Competing Interest Statement

The authors have declared no competing interest.

### Summary of Updates

The revised manuscript includes additional experiments and results. Figures have also been consolidated and include additional computational analyses.

