## Supplemental figures (S1-S16) and table (S1) for "Somatic DNA-repeat expansions initiate distinct yet analogous changes in Huntington’s disease cortical and striatal neurons"

Contains Table S1 and Figures S1-S16

**Table S1**. Cortical areas analyzed by snRNA-seq. Related to Figures 2 and 3.

| **Area** | **Name** |
|---|---|
| **BA4** | Primary Motor Cortex (precentral gyrus) |
| **BA32** | Dorsal Anterior Cingulate Cortex |
| **BA46** | Dorsolateral Prefrontal Cortex (dlPFC) |
| **BA17** | Primary Visual Cortex (V1) |
| **BA19** | Associative Visual Cortex (primarily V3) |
| **BA37** | Fusiform Gyrus (higher associative visual) |
| **BA41** | Primary Auditory Cortex (A1) |
| **BA42** | Associative Auditory Cortex (A2) |
| **BA22** | Middle Temporal Gyrus (higher order auditory) |
| **BA3** | Primary Somatosensory Cortex (S1) |
| **BA40** | Associative Somatosensory Cortex (S2) |
| **BA7** | Superior Parietal Gyrus (multisensory associative) |

**Figure S1**. Somatic CAG-repeat expansion in specific types of cortical GABAergic neurons. Related to **Fig. 1,** here focusing on specific types of interneurons. Each row/color corresponds to a different type of GABAergic neurons. Data for eight donors (ordered by CAP score, an estimate of disease progression) are shown. LAMP5+ interneurons consistently show the most somatic expansion, though few of them have expanded their CAG-repeat tracts beyond 100 CAGs.

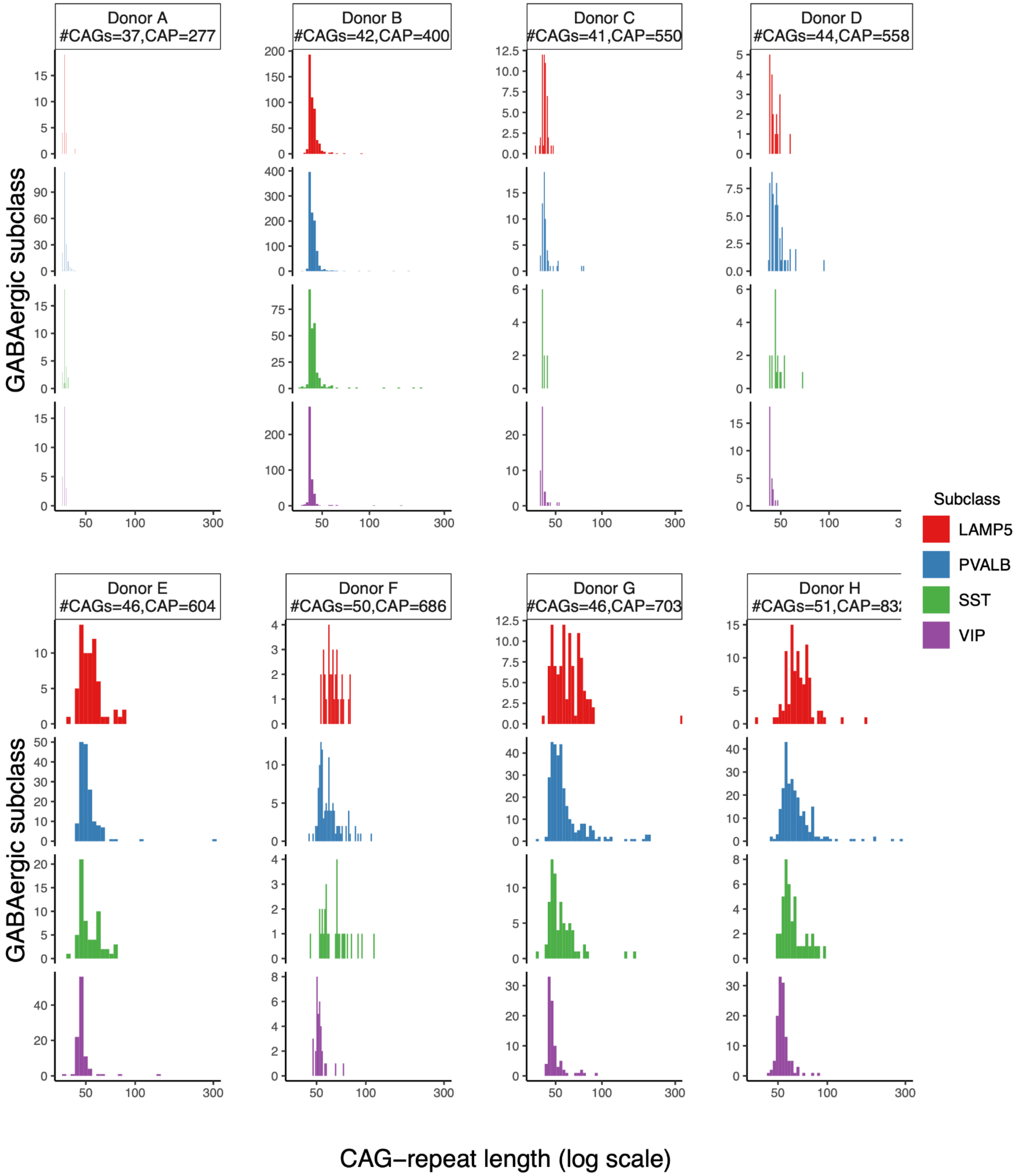

**Figure S2**. Replication across brain donors of estimates of the relative amounts of somatic *HTT* CAG-repeat expansion in different pyramidal neuron subtypes. Related to **Fig. 3**, which focused on data from an individual brain donor (the "pan-cortex donor, included again here on the y-axes for comparison to the other donors). Each circle corresponds to a pyramidal neuron subtype (from the high-resolution Siletti taxonomy) for which we have quantified somatic CAG-repeat expansion. These 20+ pyramidal subtypes are each indicated with a circle; the colors of the circles reflect supertypes (L2/3IT, etc.) from the earlier, lower-resolution (but more extensively annotated) Hodge taxonomy. On the y-axes are estimates from a deeply sampled brain donor from whose neocortex we sampled twelve cortical areas. On the x-axes are estimates from analyzing other brain donors, focusing on estimates from motor cortex (BA4, panel **a**) or anterior cingulate cortex (BA32, panel **b**). Note that, for L6b and L5/6NP pyramidal subtypes (purple and green circles), loss of the neurons with the most somatic expansion during the course of HD (see **Fig. 6**) would cause us to under-estimate their somatic expansion, by an amount that varies from donor to donor; this may be why these purple and green points diverge from the otherwise-linear replication relationship formed by the other pyramidal subtypes. Also note that, because some donors have more somatic expansion than others (due e.g. to age and genetic modifiers), the cardinal values of these estimates are donor-specific – it is their *relative* (not absolute) values that appear to be consistent across people.

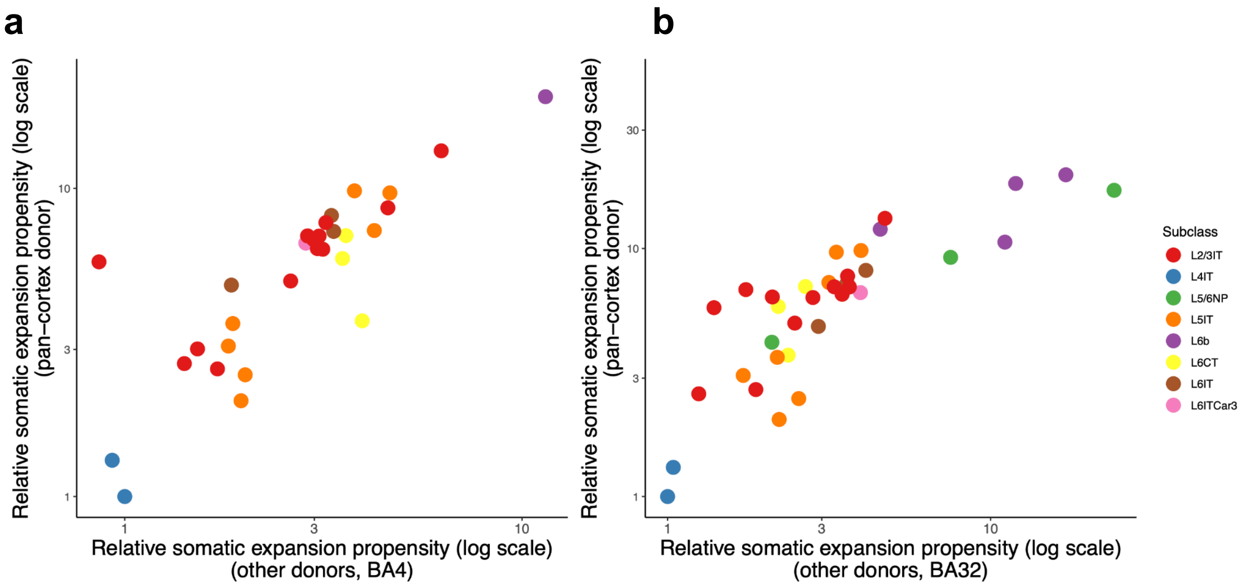

**Figure S3**. Distributions of somatic CAG-repeat tract lengths, focusing on the HD-causing allele, in subtypes of upper-layer pyramidal neurons. Related to **Fig. 3**. The nuclei were sampled from a person with HD (48yo, inherited 42-CAG allele, Vonsattel grade HD-1, "Donor B" in **Fig. 1**).

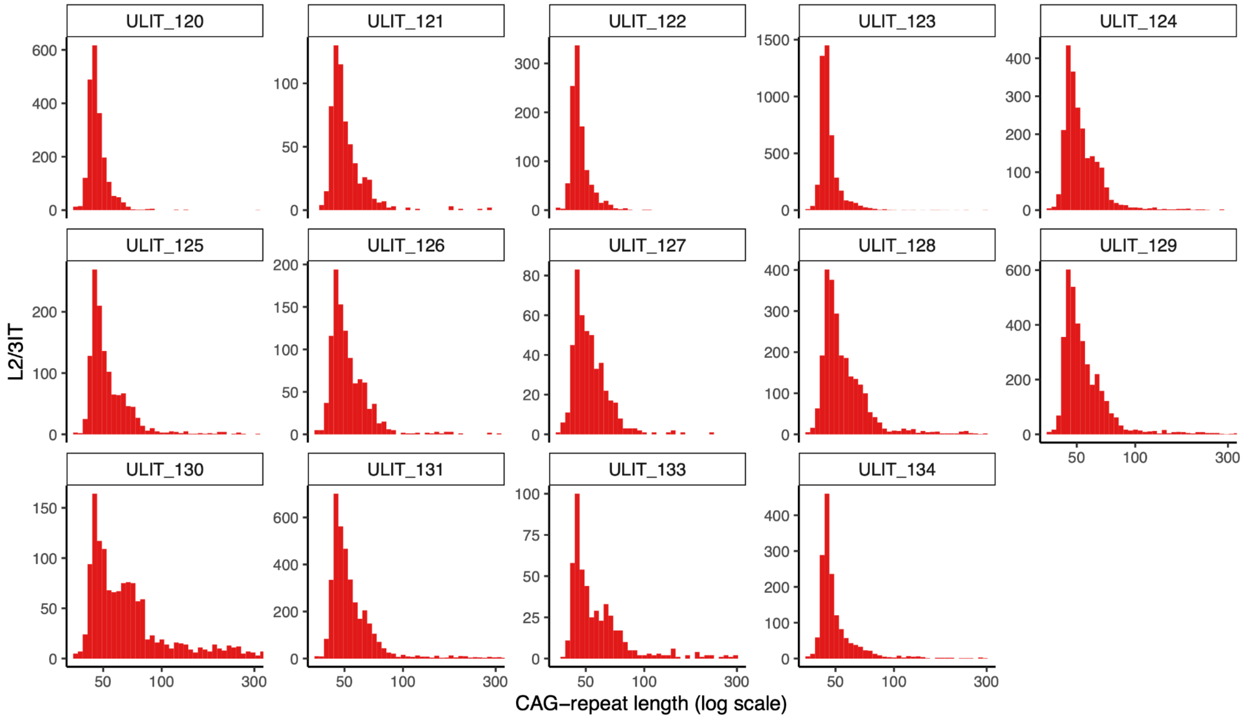

**Figure S4**. Distributions of somatic CAG-repeat tract lengths, focusing on the HD-causing allele, in subtypes of deep-layer near-projecting (L5/6NP) pyramidal neurons. Related to **Fig. 3**. The nuclei were sampled from a person with HD (48yo, inherited 42-CAG allele, Vonsattel grade HD-1, "Donor B" in **Fig. 1**).

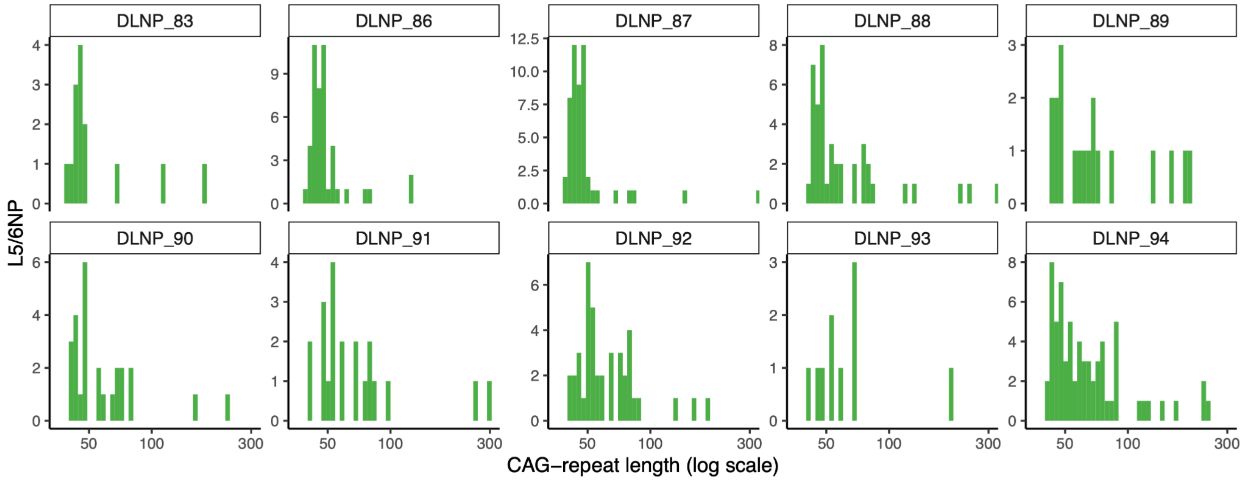

**Figure S5.** Relative rates of somatic *HTT* CAG-repeat expansion across pyramidal neuron types and subtypes, in relationship to nuclear RNA expression levels of *HTT* and a variety of genes known or proposed to regulate the stability of the *HTT* CAG repeat. Each point represents a type or subtype of pyramidal neuron; the points' y-axis locations reflect relative amounts of somatic expansion, and their x-axis locations reflect nuclear RNA expression levels (measured in UMIs per 100k UMIs samples) of the gene shown.

(**a**) Analysis of high-level pyramidal neuron types. Note that rigorous statistical analysis of such relationships is challenging, because the individual points (cell types) are non-independent: cell types have a complex, structured, and partially hierarchical set of similarity/dissimilarity relationships, and cannot be considered independent in the way that statistical tests of correlation assume. We have not reported p-values because p-values from statistical tests that do not incorporate these complex relationships are misleading, but can report that, even in a naive analysis, none reached nominal significance in parametric or non-parametric correlation tests.

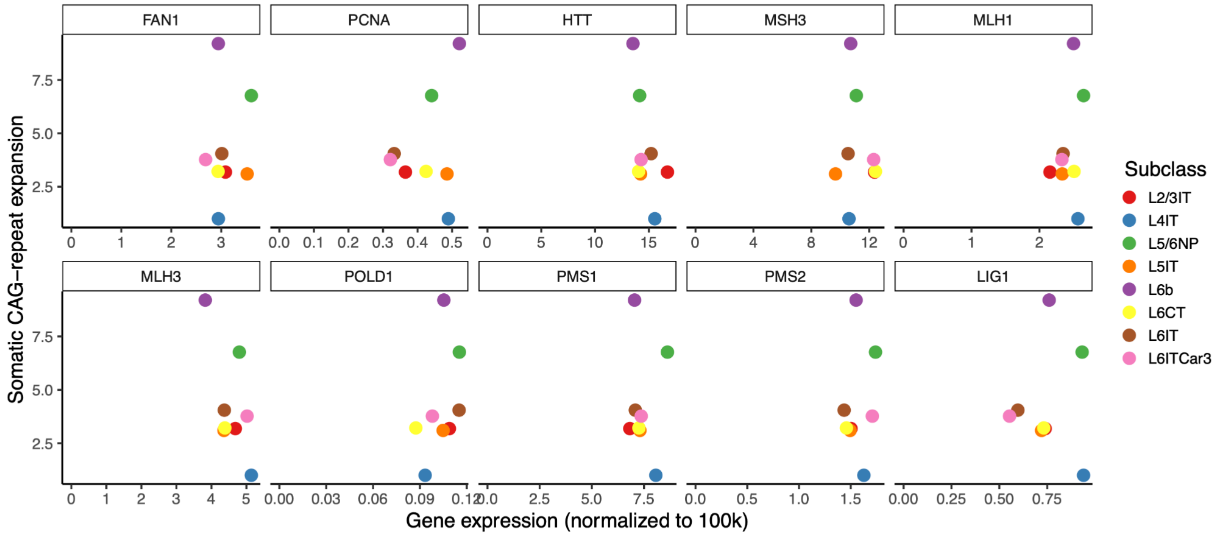

(**b**) As in (a), but dis-aggregated into pyramidal subtypes. Note that here, the statistical challenge posed by the hierarchical similarity-dissimilarity relationships of the cell types is particularly acute: for example, *PMS1* may have hints at an upward-sloping relationship, though these are driven by subtypes of L5/6NP neurons (which are similar to one another and thus non-independent observations). Some L5/6NP subtypes (green points) with particularly high relative somatic expansion (>8) tended to have relatively high expression of several mismatch repair genes (e.g. *MLH3, MLH1, PMS1, LIG1*), though again, these closely related neuronal subtypes cannot be considered independent observations.

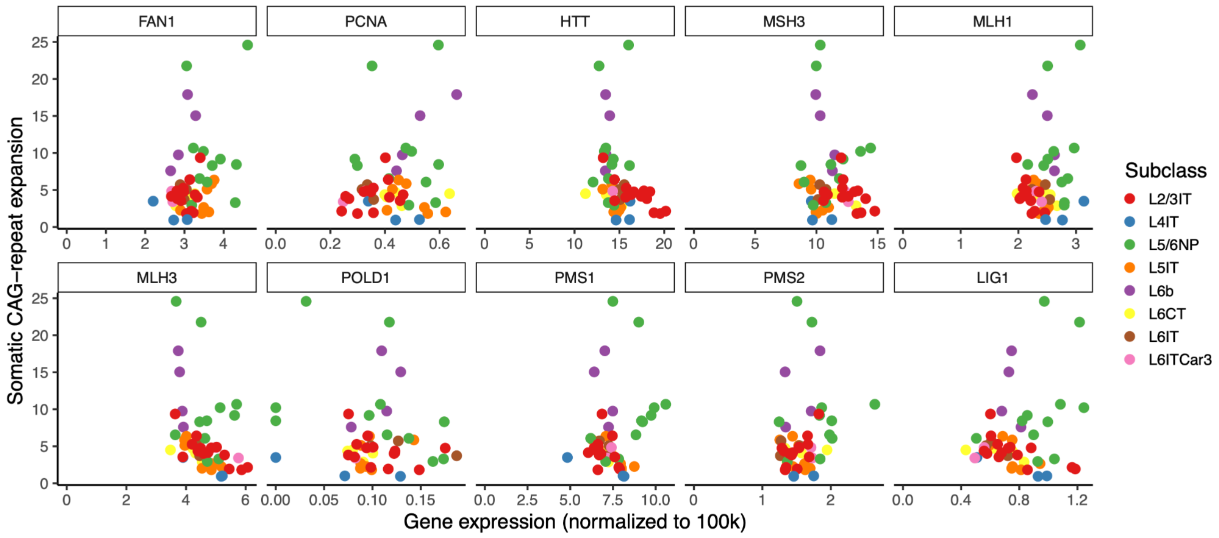

It is important to remember that individual genes tend to change in expression in the context of large gene-expression programs that involve concerted changes in the expression of very many other genes – often hundreds of genes – together. Thus, expression measurements for a single gene are also indirect proxies for many other genes' expression levels, and for unmeasured cellular properties (such as neuronal activity patterns or metabolic states) that may also affect somatic instability. In this context, one should be reluctant to attribute causality to individual genes based on observational/correlational analyses.

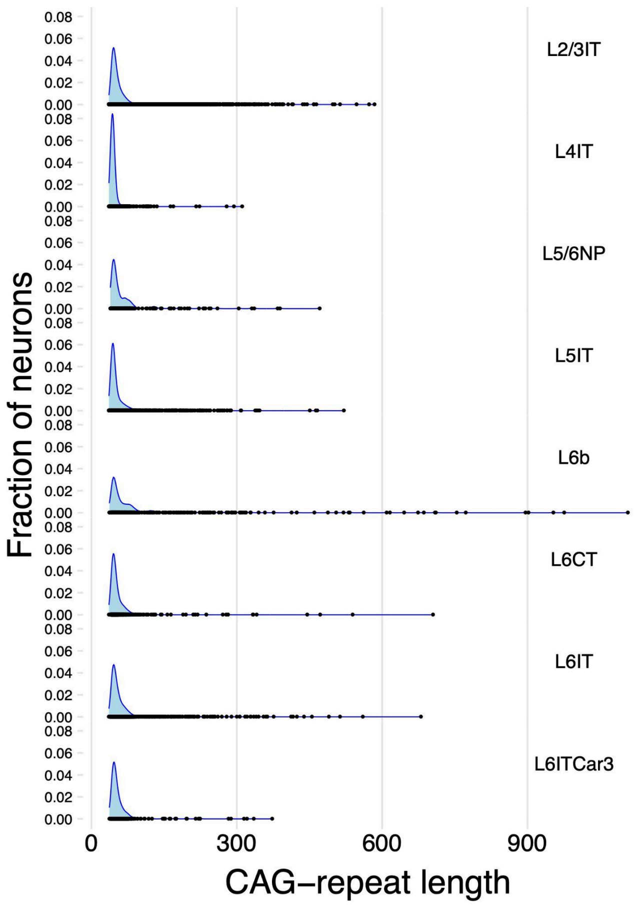

**Figure S6**. Distributions of CAG-repeat lengths in eight types of cortical pyramidal neurons from the same person with HD. Blue shaded areas are smoothed density estimates of the repeat-length distribution. Overplotted black points show the CAG-repeat length measurements in individual glutamatergic neurons. In all eight glutamatergic neuron subclasses, the repeat-length distribution exhibits an armadillo-like shape, in which the DNA-repeat tract in most neurons has undergone modest expansion (up to about 90 CAGs) but in a small fraction of neurons it has undergone far greater expansion (up to 500+ CAGs).

**Figure S7**. Phase C gene-expression changes appear in cortical neurons with somatic CAG-repeat expansion beyond 150 CAGs. Related to **Fig. 4**, and extending analysis to additional persons with HD. Analyses of (**a**) motor cortex (BA4) and (**b**) anterior cingulate cortex (BA32). On each pair of panels, each pyramidal neuron is represented by both a blue point (lower plot) and an orange point (upper plot): blue points show the median fold-change of a set of 115 genes that decrease in expression with CAG-repeat expansion (C- genes); orange points show the median fold-change of a set of 142 genes that increase in expression with CAG-repeat expansion (C+ genes).

**a**

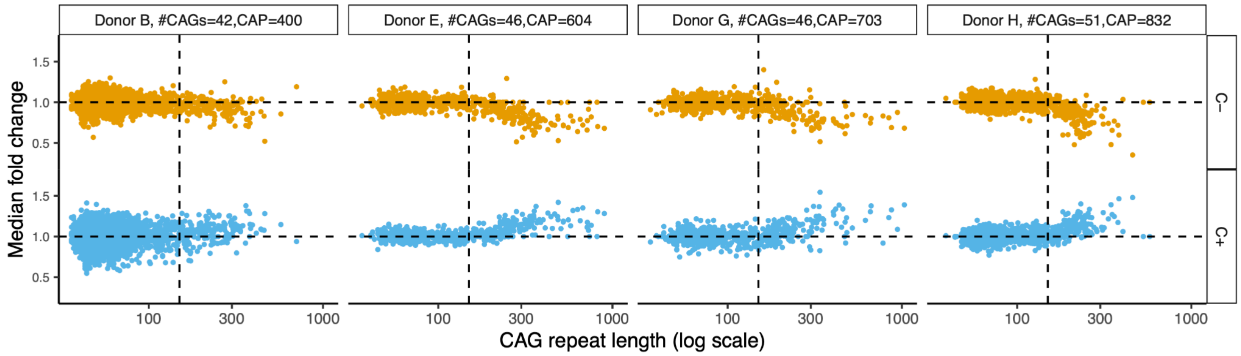

**b**

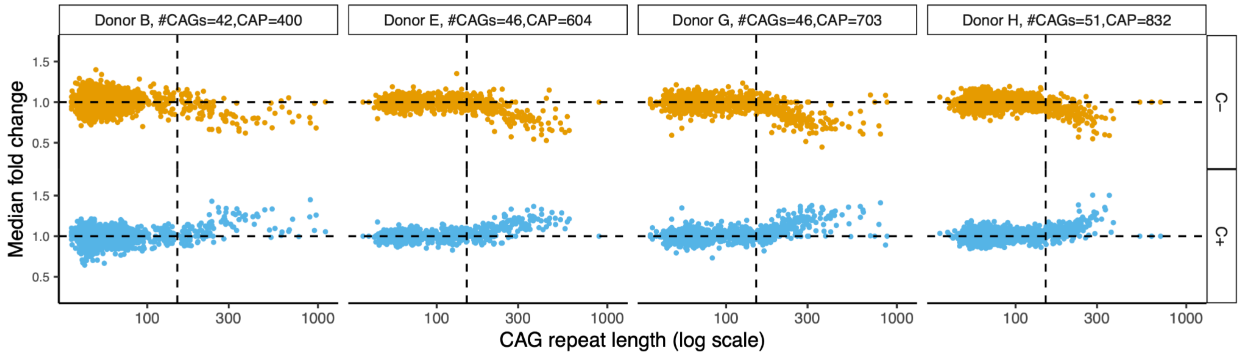

**Figure S8**. Gene-expression changes with somatic CAG-repeat expansion beyond 150 CAGs in each type of pyramidal neuron. Related to **Fig. 4**, but here drawing upon the 12 cortical areas sampled in one of the donors. Each of the eight plots shows the data for a specific type of cortical pyramidal neuron. On each plot, each neuron is represented by both a blue point and an orange point: orange points show the median fold-change of a set of 144 genes that decrease in expression with CAG-repeat expansion (C- genes); blue points show the median fold-change of a set of 179 genes that increase in expression with CAG-repeat expansion (C+ genes). The same genes were used for the calculation in each panel, and the 12 sampled cortical areas were used together. **Fig. 4a** contains analogous analyses of nuclei sampled from other persons with HD.

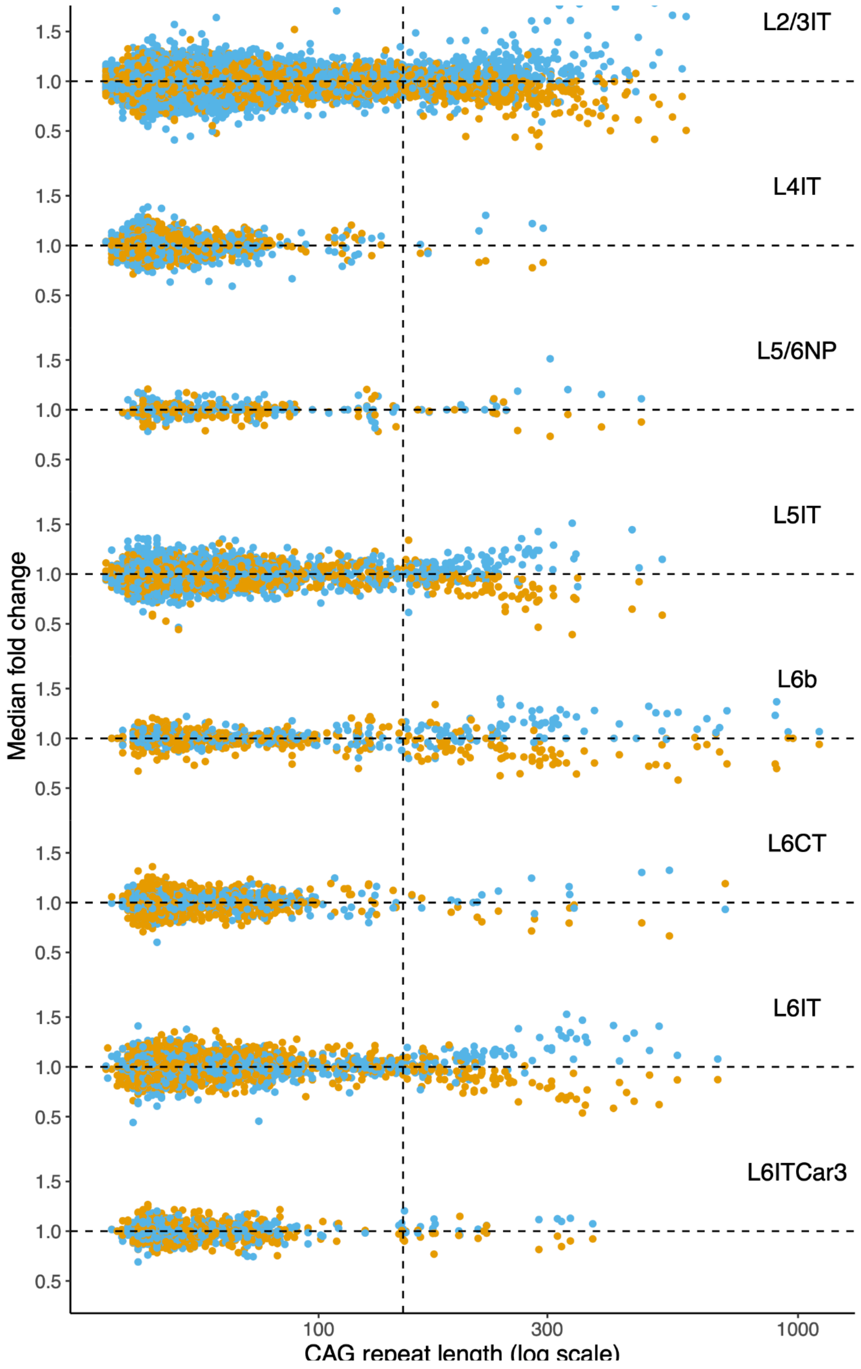

**Figure S9**. Phase D gene-expression changes. Related to **Fig. 5**, and drawing upon additional brain donors with HD. (**a**) Phase D changes in additional brain donors (shown all together). Points represent individual neuronal nuclei. y-axis: de-repression score, the number of transcripts (UMIs) detected from 59 phase D genes that are normally silent in cortical projection neurons. (**b**) Fraction of pyramidal neurons that have entered phase D, in relationship to CAG-repeat length (binned). (**c,d**) Sharing of Phase D de-repressed (D++) genes between striatal and cortical neurons. There were 124 phase D++ genes that had (in either cortex or striatum) the properties of being (i) silent at baseline (i.e. in neurons with <150 CAGs), and (ii) expressed in phase D neurons. Plots in panels c and d show how these 124 genes changed in expression in separate analyses of phase D striatal neurons (panel c) and of phase D cortical neurons (panel d). For each gene (point), location on the x-axis shows the log-fold-change when comparing phase D neurons to neurons at baseline (<150 CAGs); the y-axis shows the statistical significance of the comparison (Wilcoxon test). The color of the points reflects the fraction of neurons (with <150 CAGs) in which any RNA from the gene was detected. Note that the genes that fail to show de-repression in cortical neurons (the points below the dashed line in panel d) are for the most part genes that are already expressed in cortical neurons at baseline (as shown by the colors of the points). These include genes encoding the cortical transcription factors NFIX, LHX2, TBR1, TLE3, and TLX2, as well as genes with other neuron-type-specific functions (SLC17A7, KCNC2, ALDOC, PMS3, ARL4C).

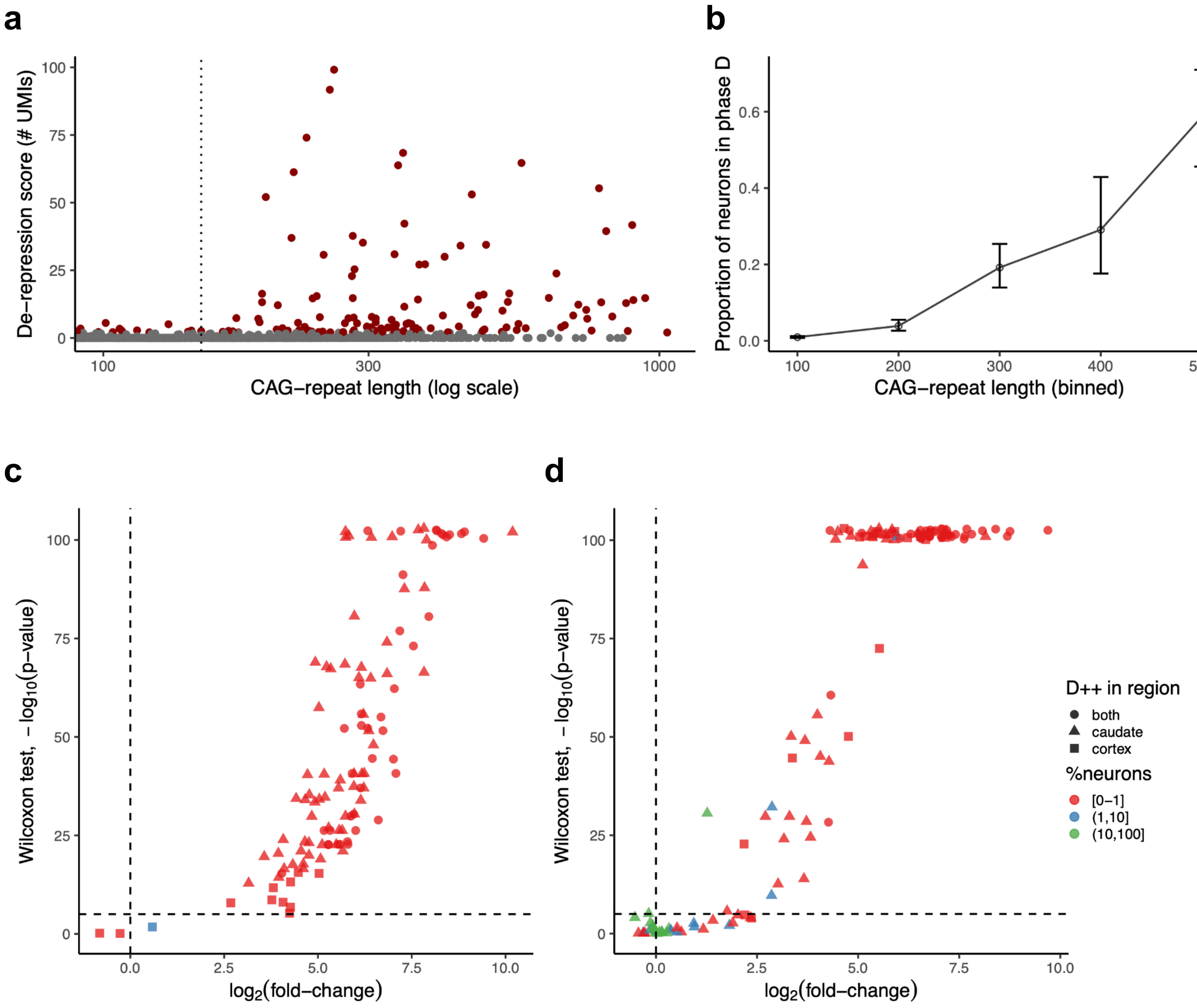

**Figure S10.** Attrition of cortical interneurons due to age but not HD. Related to **Fig. 6**.

(**a**) Cortical interneurons (GABAergic neurons) exhibited no consistent difference in abundance between 50+ brain donors with HD and 50+ unaffected brain donors – in contrast to L5/6NP and L6b pyramidal neurons (leftmost panels). Results are from motor cortex (BA4, upper set of panels) and anterior cingulate cortex (BA32, lower set of panels). The p-values are calculated from the Wilcoxon (non-parametric) test so as to be resistant to outliers and are nominal p-values (not corrected for the number of cell types tested). Note that the abundances of several types of GABAergic interneurons exhibit striking inter-individual variation, but that (in this 50 HD vs. 50 unaffected analysis) the differences between HD-affected and unaffected individuals were not even nominally significant in either BA4 or BA32.

**a**

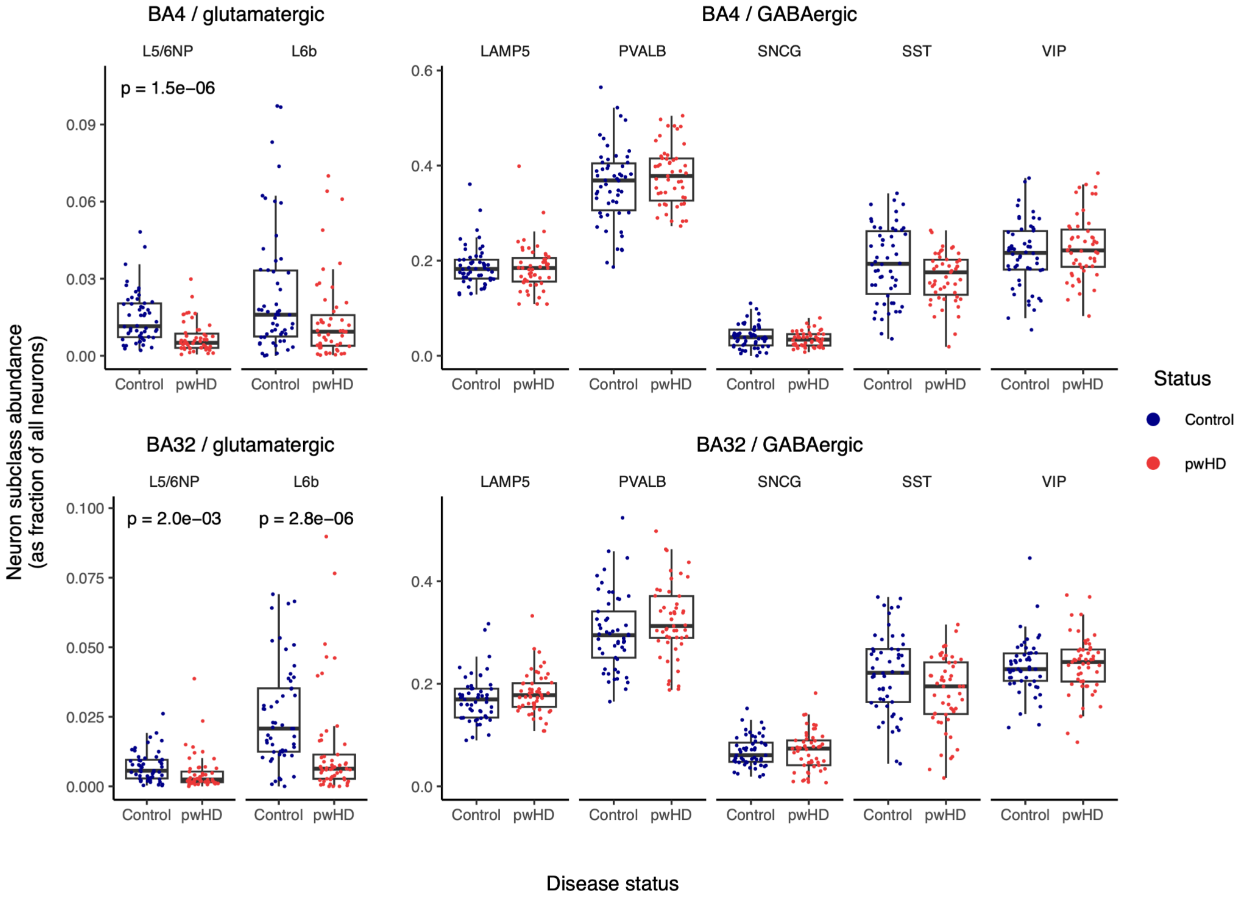

(**b**) Attrition of SST and VIP interneurons with age, in both BA32 (upper panels) and BA4 (lower panels), among both unaffected persons (green points) and persons with HD (purple points). The green and purple regression lines (and correlation statistics) were separately fit to the data from unaffected persons (green) and persons with HD (purple).

**b**

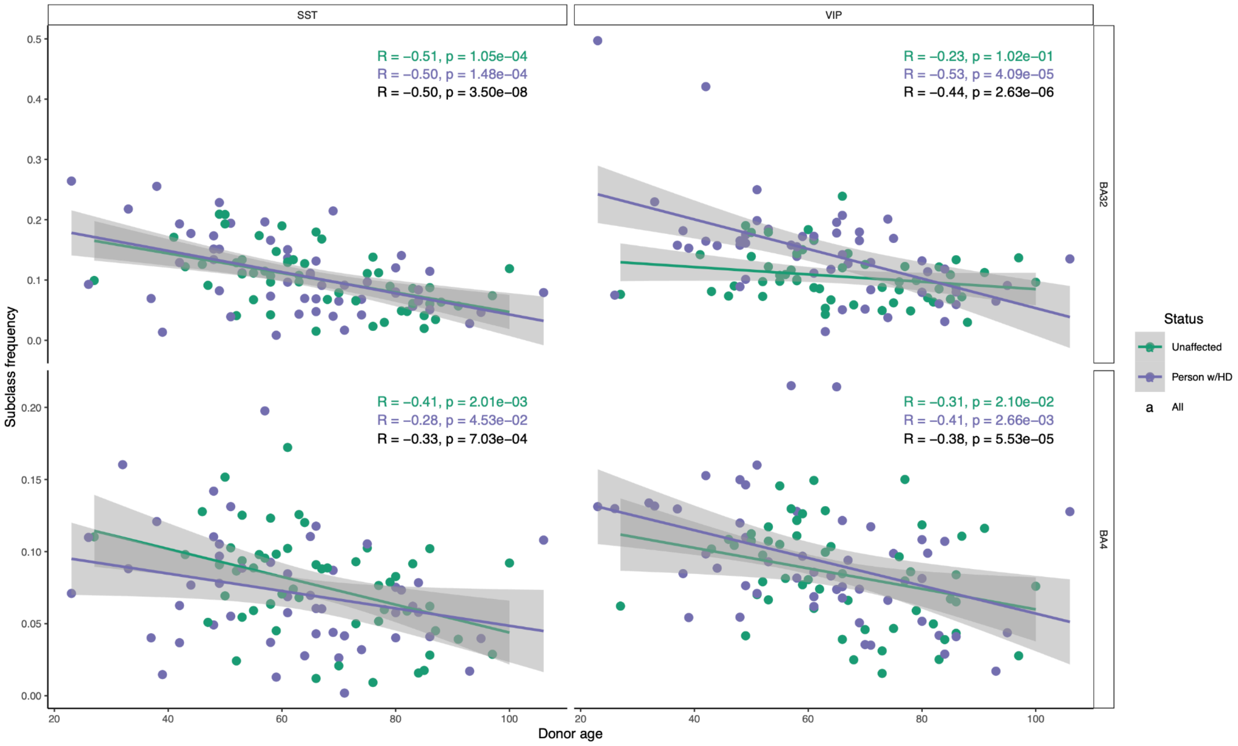

**Figure S11.** Replication of case-control differentially expressed genes (DEGs) across all types of pyramidal neurons, regardless of their vulnerability in HD. Related to **Results** section on non-cell-autonomous sequelae of neurodegeneration. DEGs were identified by comparing gene expression in persons with HD vs. unaffected individuals in each of the pyramidal neuron subclasses. Each point corresponds to a gene found to be differentially expressed in the current work's differential-expression analyses of persons with HD (relative to unaffected individuals) in at least two pyramidal neuron subclasses and is also present in the list of differential expressed genes for L5a in the study by Pressl et al; expression fold-changes (HD/unaffected) are shown for pairs of pyramidal neuron types. The locations of points (genes) on the plot are based on fold-change in the current data (comparing HD-affected to unaffected brain donors). Points are colored blue or orange based on whether they were found to be up- or down-regulated in HD-impacted neocortex in the study by Pressl et al.

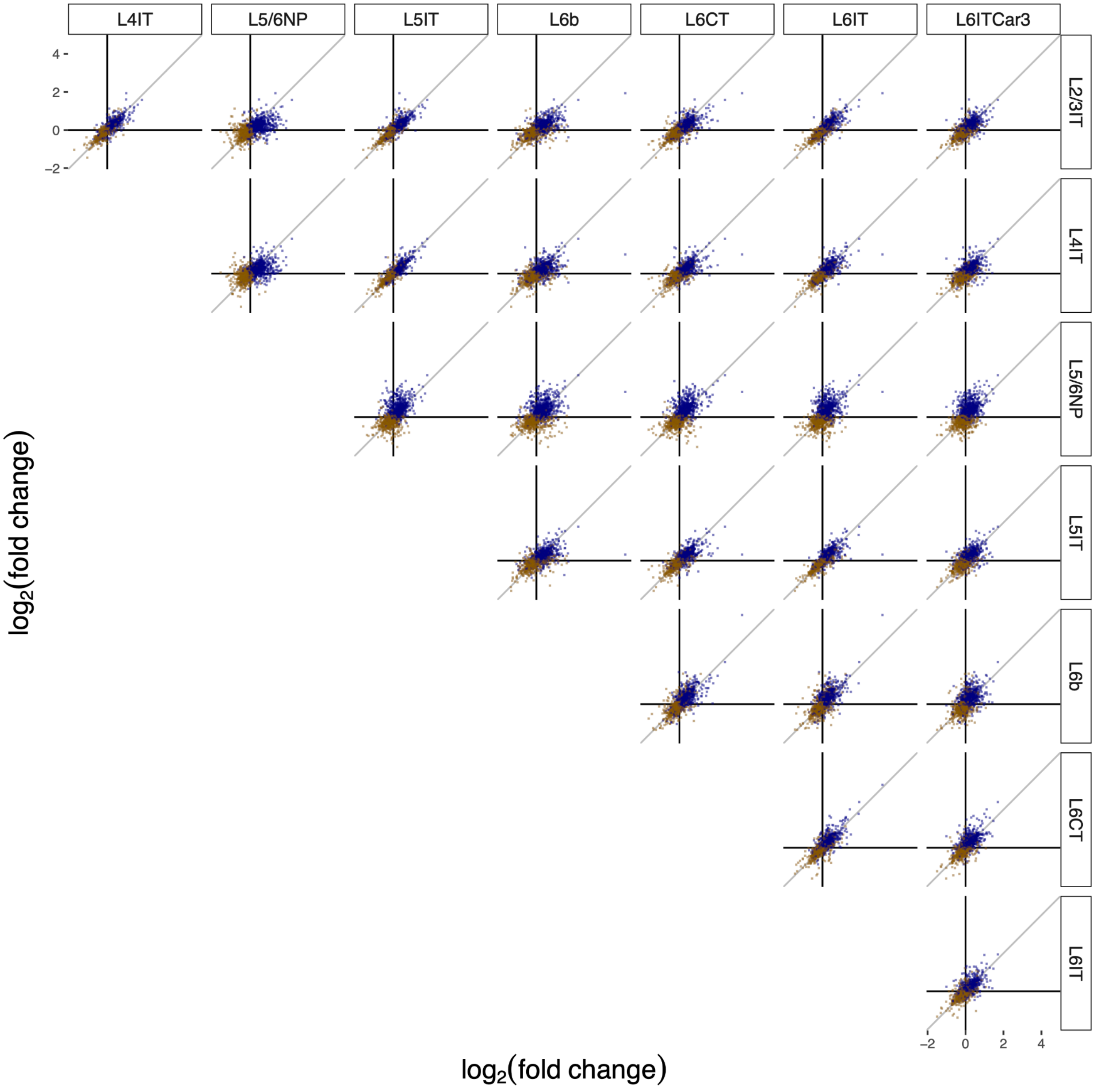

**Figure S12.** Similar to **Fig. S11**, but subsetting to the set of genes in the "synapse" Gene Ontology cellular component category. The locations of points (genes) on the plot are based on fold-change in our own data (comparing HD-affected to unaffected brain donors). Points are colored blue or orange based on whether they were found to be up- or down-regulated in HD-impacted neocortex in the study by Pressl et al.

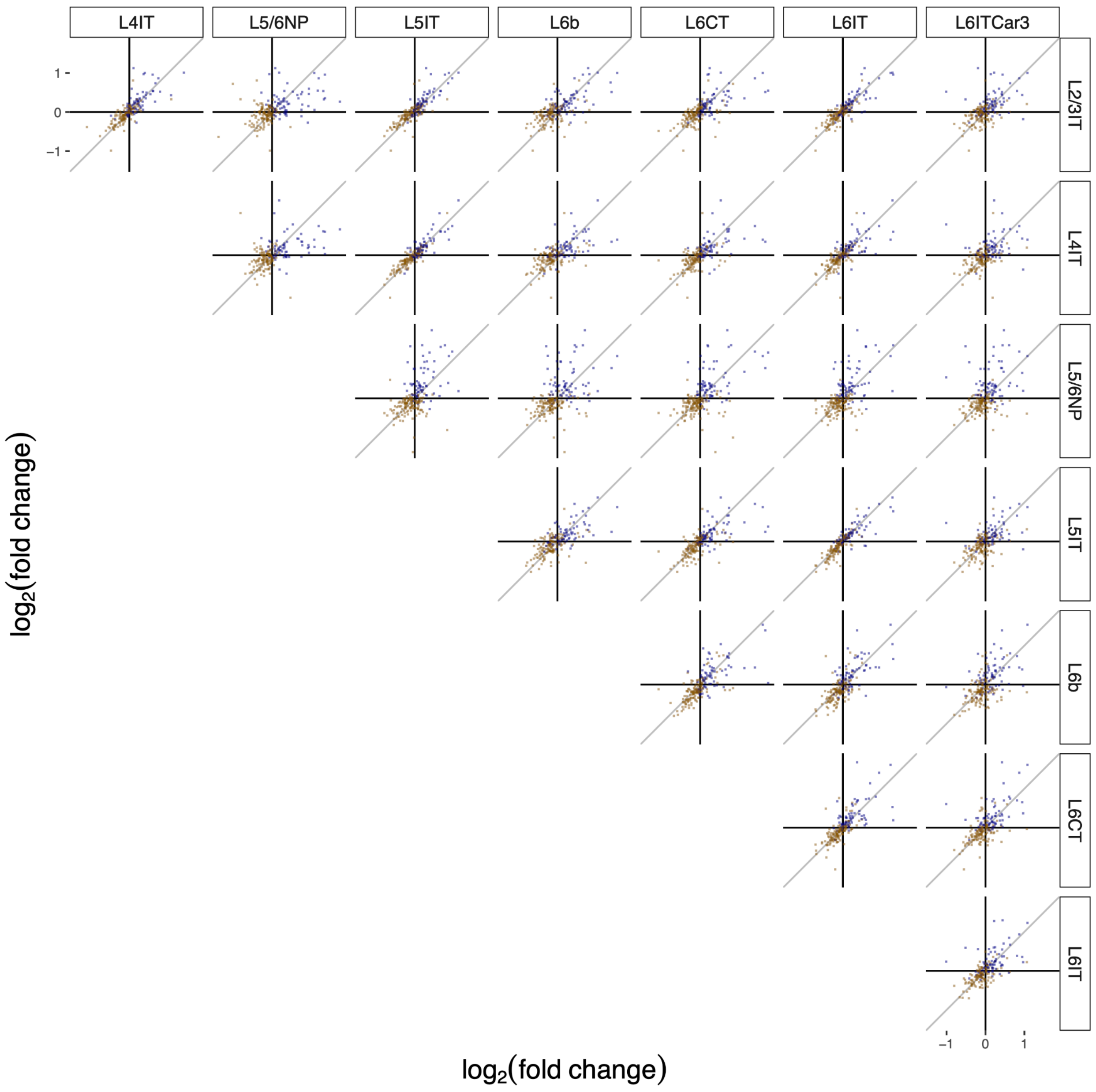

**Figure S13.** Relationship of six types of gene-expression changes to length of a pyramidal neuron's CAG-repeat tract. Related to **Results** section on non-cell-autonomous sequelae of neurodegeneration. Gene sets considered, from left to right: phase C genes (C- and C+ denote down- and up-regulated genes); phase D genes (D- and D+); and "differentially expressed genes" in L5a pyramidal neurons identified in the study by Pressl et al (DEGs, divided into DEG- and DEG+ gene sets based on whether expression tended to increase or decrease in HD-affected relative to unaffected neocortex). Each of these gene sets was filtered to contain genes expressed in at least 50% of all pyramidal neurons. The quantity plotted is the median fold-change among those genes in the set. The numbers in red above the bar plots show the numbers of genes in each filtered gene set.

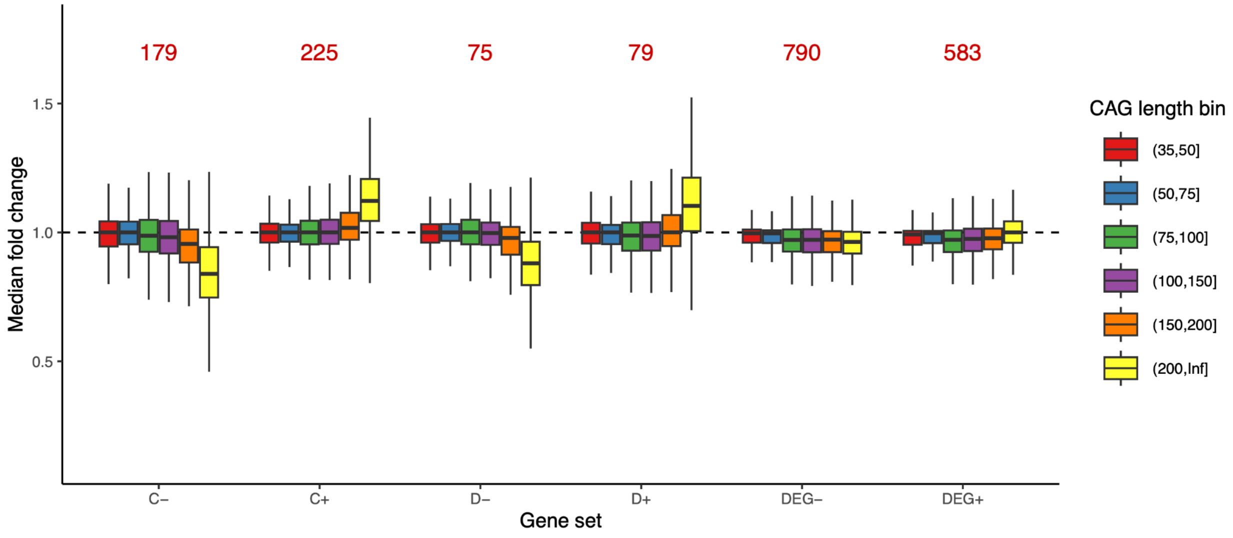

**Figure S14.** Sorting gating strategy for isolating nuclei with HTT inclusions. Related to **Fig. 7**. (**a**) Representative FANSII gating of nuclei isolated from a control donor (left) and a donor with HD (right). Nuclei containing HTT inclusions were gated as HTT-high and ER-Tracker-low to minimize potential autofluorescence and non-specific fluorophore binding associated with residual endoplasmic reticulum content. Notably, few to no nuclei fall within this gate in the sample from the unaffected (control) brain donor. (**b**) Representative immunofluorescence images of nuclei from unsorted (left) and sorted (right) fractions from an HD donor using this gating strategy. Scale bar = 25 μm.

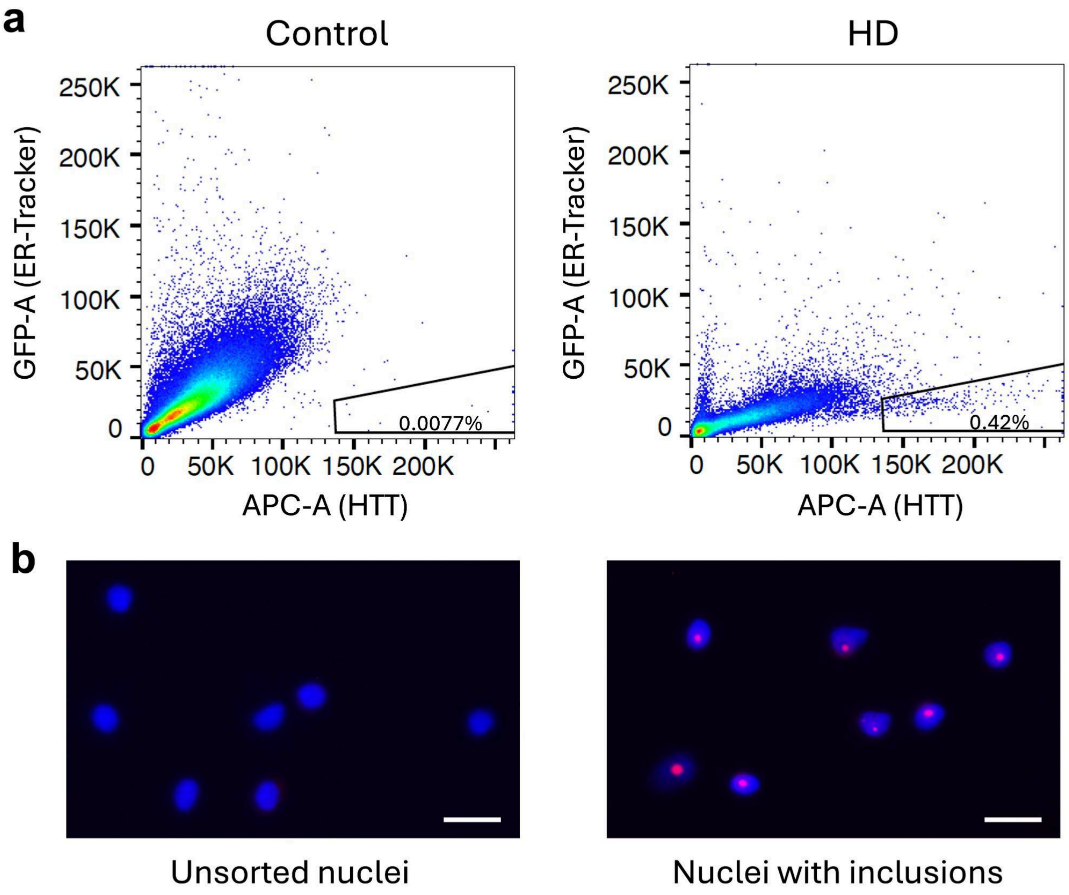

**Figure S15.** Representation of pyramidal neuron types among nuclei with HTT inclusions, in comparison to all (unsorted) neuronal nuclei. Related to **Fig. 7c**, here expanding to consider the various types of pyramidal neurons. The sorted and unsorted nuclei were analyzed by snRNA-seq; subtype identities were inferred from the genome-wide RNA data in each individual nucleus. Here the pyramidal neuron types have been ordered by their relative amounts of somatic expansion (inferred from earlier analyses in other donors, shown in **Fig. 2c**), from least expansive (L4IT, blue, top) to most expansive (L6IT, L6CT, L5/6 NP, L6b; brown, yellow, green, purple, bottom). Note that this analysis reflects subtype composition at a specific time point (end of life) and therefore has an indirect and potentially complex relationship to cumulative cell loss through the course of illness; for example, L5/6 NP neurons were already so depleted as to be less than 1% and 0.1% of pyramidal neurons in the two donors (in unsorted nuclei as well as sorted nuclei with HTT inclusions).

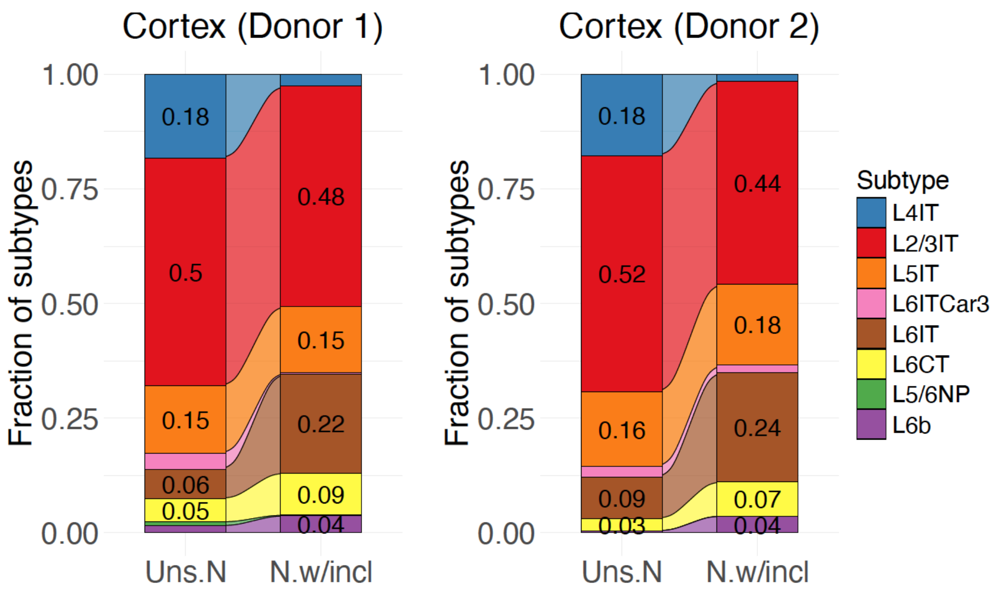

**Figure S16**. Gene expression changes in neuronal nuclei with inclusions. Related to **Fig. 7**. (**a-c**) Phase C and D gene-expression changes in nuclei with inclusions (red points) in comparison to unsorted nuclei (grey points). From left to right: expression changes in individual nuclei for phase C- genes , phase C+ genes, and phase D++ genes. (Expression changes for C- and C+ genes are quantified in terms of median fold-change relative to baseline expression level; and for D++ genes, which are non-expressed or negligibly expressed at baseline, in terms of the total summed number of UMIs (per 100k) ascertained across these 59 genes.) Data are from the same experiments as in **Fig. 7**, but here including only those nuclei with CAG-repeat measurements for the HD-causing allele. The fold-change calculations are described in Methods, and the genes used in the calculations are listed in Supplementary Data. (**d-f**) Genome-wide comparisons of gene-expression levels between nuclei with inclusions (x-axis) and unsorted neuronal nuclei (y-axis). Phase C-, C+, and D++ genes are shown with orange, blue and red points respectively; all other genes are shown with gray points.

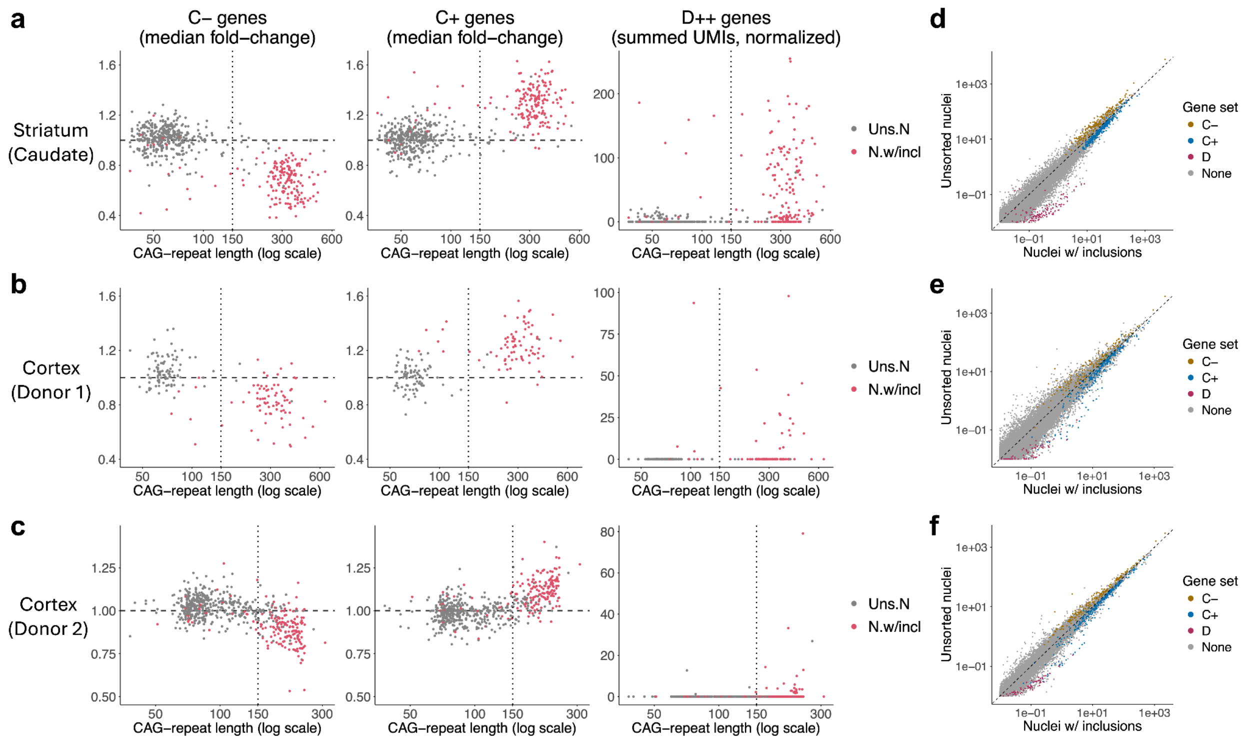
